# Analysis of Mammalian Succinate Dehydrogenase Kinetics and Reactive Oxygen Species Production

**DOI:** 10.1101/870501

**Authors:** Neeraj Manhas, Quynh V. Duong, Pilhwa Lee, Jason N. Bazil

## Abstract

Succinate dehydrogenase is an inner mitochondrial membrane protein complex that links the tricarboxylic acid cycle to the electron transport system. It catalyzes the reaction between succinate and ubiquinone to produce fumarate and ubiquinol. In addition, it can produce significant amounts of superoxide and hydrogen peroxide under the right conditions. While the flavin adenine dinucleotide (FAD) is the putative site of reactive oxygen species production, free radical production from other sites are less certain. Herein, we developed a computational model to analyze free radical production data from complex II and identify the mechanism of superoxide and hydrogen peroxide production. The model includes the major redox centers consisting of the FAD, three iron-sulfur clusters, and a transiently catalytic bound semi quinone. The model consists of five-states that represent oxidation status of the enzyme complex. Each step in the reaction scheme is thermodynamically constrained, and transitions between each state involve either one-electron or two-electron redox reactions. The model parameters were simultaneously fit using data consisting of enzyme kinetics and free radical production rates under a range of conditions. In the absence of respiratory chain inhibitors, model analysis revealed that the 3Fe-4S iron-sulfur cluster is the primary source of superoxide production followed by the FAD radical. However, when the quinone reductase site of complex II is inhibited or the quinone pool is highly reduced, superoxide production from the FAD site dominates at low succinate concentrations. In addition, hydrogen peroxide formation from the complex is only significant when these one of these conditions is met and the fumarate concentrations is in the low micromolar range. From the model simulations, the redox state of the quinone pool was found to be the primary determinant of free radical production from complex II. This study highlights the importance of evaluating enzyme kinetics and associated side-reactions in a consistent, quantitative and biophysical detailed manner. By incorporating the results from a diverse set of experiments, this computational approach can be used to interpret and explain key differences among the observations from a single, unified perspective.

## Introduction

Succinate dehydrogenase (SDH) is a heterotetrametric protein attached to the inner membrane of mammalian mitochondria. It links the tricarboxylic acid cycle (TCA) to the electron transport system (ETS) [1, 2] and is comprised of four subunits. The flavoprotein subunit SDHA contains a flavin adenine dinucleotide (FAD) covalently bound to the active site. The iron-sulfur protein subunit SDHB contains three iron-sulfur clusters: [*2Fe-2S*], [*4Fe-4S*], [*3Fe-4S*] (ISCs). Subunits C and D (SDHC and SDHD) contains two transmembrane cytochrome *b hemes* [3, 4]. These two *hemes* subunits along with the interface of SDHB form the ubiquinone (Q) binding site (Q site) [4, 5]. Complex II couples the oxidation of succinate to fumarate with the reduction of Q to ubiquinol (QH_2_). The overall catalytic reaction is given by Eq. 1.

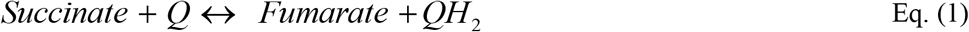

For each molecule of succinate oxidized, FAD is reduced with two electrons to form the fully reduced flavin (FADH_2_). The subsequent transfer of electrons through the iron-sulfur clusters occur one at a time. The first electron is transferred to the [*2Fe-2S*] iron-sulfur center from FADH_2_, producing a flavin radical (FADH^•^). When the [*2Fe-2S*] ISC becomes oxidized, the flavin radical passes the second electron to this ISC and becomes fully oxidized (FAD). Similarly, consecutive one-electron transfers from the ISCs to the Q site and reduces Q to QH_2_. As a part of this process, a stable semiquinone (SQ) is formed [4, 6, 7]. The FADH^•^, FADH_2_ and SQ are putative sources of reactive oxygen species (ROS) in complex II [8–10]. However, the [*3Fe-4S*] ISC has also been speculated to be a source of ROS [11]. In this paper, ROS refers to either superoxide (O_2_^•-^) or hydrogen peroxide (H_2_O_2_) and will be explicitly defined where needed for clarity. Historically considered unavoidable byproducts of aerobic respiration, ROS has been recently recognized as signaling molecules at low concentrations [12, 13]. However, under certain pathological conditions including ischemia/reperfusion (I/R) injury, ROS can increase to levels that contribute to cell death [14–18]. In mammalian mitochondria, 11 sites are known to produce ROS [9]. Complex III is argued as the primary ROS producer under resting condition, and complex I becomes the significant source of ROS under many pathological conditions. Early studies suggested that complex II is not a significant contributor of ROS [18–20]; however, more recent studies have shown that it may become an important source of ROS under a variety of physiological and pathological settings [9, 17, 21–23]. For instance, when complexes I and III are inhibited and the succinate concentration is low (~100 μM), complex II produces ROS at high rates relative to complexes I and III [21, 24]. In addition, skeletal muscle mitochondria respiring on succinate have been shown to produce high rates of ROS [25]. That said, the quantitative contribution to total mitochondrial ROS production from complex II remains to be elucidated.

As stated above, the flavin and quinol binding sites are believed to be the sources of ROS produced by complex II [8–10]. In one study, the flavin site of complex II is reported to produce comparable amount of ROS to the quinone binding site of complex I under resting condition [8]. Recently, Quinlan *et al.* demonstrate that complex II from rat skeletal muscle mitochondria produce ROS at low levels of succinate or when QH_2_ oxidation is inhibited [21, 26]. These studies suggest that most of the ROS from complex II originates from the flavin site [21]. In another study using bovine sub-mitochondrial particles (SMP), Siebels and Drose show that complex II generates ROS when its Q site is inhibited by atpenin [27]. Evidence supporting ROS production at the ISC near the Q site comes from a recent study by Grivennikova *et al*. [11]. However, since these studies are performed under different experimental conditions, the precise mechanism underlying ROS generation by complex II remains unclear. Specifically, the sites producing ROS are an ongoing topic of debate.

Complex II contributes to oxidative stress that occurs in early I/R injury by enabling succinate accumulation during ischemia [28]. During ischemia, complex II reverses due to excess fumarate produced by purine nucleotide breakdown [15]. Succinate accumulation is possible because complex I regenerates QH_2_ needed to sustain the reverse reaction of complex II. Thus, succinate acts as the final electron acceptor instead of oxygen. Succinate accumulates in the surrounding tissue until either the source of fumarate is exhausted, or reperfusion begins. During reperfusion, complex II metabolizes the available succinate to produce QH_2_, leading to hyperpolarization of the inner mitochondrial membrane [29]. The combination of a high QH_2_ levels and hyperpolarized membrane potential drives complex I to enter the so-called reverse electron transport state and produces ROS at extremely high rates [29]. In this state, it is more accurate to describe complex I as entering a near equilibrium state [14, 30]. In this state, the redox centers on complex I are highly reduced and can lead to the generation of significant amounts of free radicals. During transition from late reperfusion to normoxia, fluxes through the ETS and TCA return to normal and ROS levels subside [15, 16]. However, depending on the duration and severity of ischemia, the accumulated ROS levels that occur during ischemia and after reperfusion determine the extent of cellular damage. Therefore, understanding the mechanism of complex II kinetics and ROS production under both physiological [12] and pathological conditions [15] is crucial to develop effective therapies capable of mitigating the detrimental consequences of I/R injury.

Herein, we have developed a computational model of complex II that is biophysically detailed and thermodynamically consistent. The model simulates the redox states of major redox centers: the FAD, ISCs and SQ bound to the Q site. By doing so, the model is capable of simulating both the kinetics of succinate oxidation and ROS production by the enzyme complex. Multiple data sets consisting of succinate oxidation and ROS production rates under different conditions were used to parameterize and validate the model. Our analysis reveals that while the FAD site does produce ROS, the primary source is the [*3Fe-4S*] ISC under normal physiological conditions. However, in the presence of respiratory chain inhibitors, the FAD site is the dominate site and produces significant levels of H_2_O_2_. Moreover, the model shows that the inhibitory effects of atpenin are not simply attributed to competitive binding at the Q site but also include allosteric effects that modulate catalytic turnover. Lastly, this model is an ideal choice for integration into large-scale models of mitochondrial metabolism to study ROS production from the ETS during physiological and pathophysiological conditions.

## Methods and Model Construction

An overview of the thermodynamically constrained, biophysically detailed model of complex II is presented in Fig. 1. The modeling approach is based on our prior work [14, 30]. In brief, structural and thermodynamic data are used to constrain the model, and enzyme state-transitions are governed by the law of mass action.

**Figure 1.**
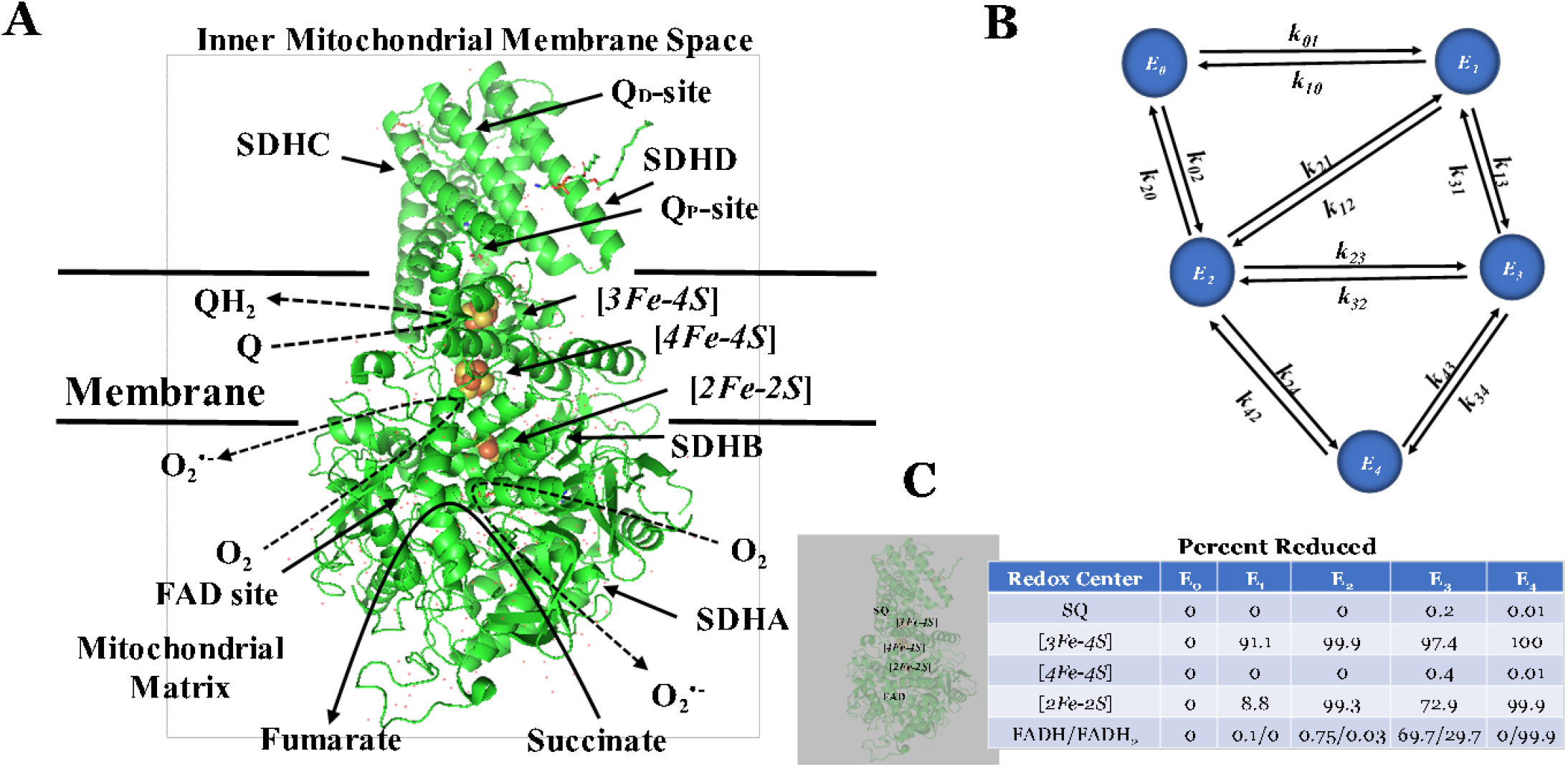
Model diagram of complex II. A) The ribbon diagram along with the major redox centers shows the enzyme consists of four subunits. SDHA constitutes a covalently bound FAD in the dicarboxylate binding site where succinate is oxidized to fumarate to generate the fully reduced flavin in a two-electron step. SDHB is the peripheral subunit that contains three iron-sulfur clusters ([*2Fe2s*], [*4Fe4s*], and [*3Fe4s*]) that transfer electrons one at a time to produce QH_2_ at the Q site located in the interface of SDHB and the *heme* group located between the integral membrane proteins SDHC and SDHD. Q_d_ and Q_p_ are distal and proximal quinone binding sites, respectively. B) State representation of the model is shown where E_*i*_, is the *i*^th^ electronic state corresponds to the number of the electrons residing in the complex’s redox centers. All state transition rates *k_ij_* are shown in the Supplementary Material. Electronic transitions can occur by either one-electron or two-electron steps. C) The percent of each redox center reduced for each electronic state in the model. A fully oxidized Q pool and pH 7 were the conditions used to calculate the percentages. The complex II ribbon diagram was generated from the crystal structure by Sun et al. [3] using PyMOL [31].

The model includes the redox biochemistry reactions that occur at the FAD, ISCs and two Q sites. The FAD site contains the binding site for succinate and other dicarboxylates. While ubiquinone and its analogues can hypothetically bind to both the proximal and distal Q sites, quinone reduction has been shown to occur at the proximal Q site, Q_p_ site. The function of the distal Q site, Q_d_ site, is still unknown [5, 32]. Reactions at these sites are assumed to be independent from each other (i.e., there are no long-distance conformational changes required for enzyme catalysis between the Q_p_ and the FAD sites). However, the Q_d_ site is assumed to exert some control over turnover at the FAD and proximal Q sites when atpenin or other molecules are bound to this site. The enzyme kinetic model consists of five electronic states which constitute the redox state of the entire enzyme. For example, all redox centers are oxidized in the E_0_ state, at least one redox center is one-electron reduced in the E_1_ state, and so on and so forth. A minimum of five electronic states were necessary to fit the experimental data. Including more did not improve fits to the data. Transitions between electronic states are governed by the Gibb’s free energies of the redox reactions involved. Two-electron reactions involve the succinate/fumarate (SUC/FUM), Q/QH_2_ and O_2_/H_2_O_2_ couples. One-electron reactions involve the O_2_/O_2_^•-^ couple. Binding of substrates, products and inhibitors at the FAD and Q sites is assumed to be faster than state transition rates. Forward state-transition rate constants are estimated from experimental data while the reverse rate constants are calculated from the forward rates and equilibrium constants for the respective reaction. The equilibrium constants are computed from the midpoint potentials taken from the literature (Table S1) and adjusted to account for the effects of pH and temperature. The rates corresponding to these reactions are shown in Supplementary Material from Eqs. S78-S91. The transitions between each electronic state is fully reversible and governed by the law of mass action.

Within each electronic state, the enzyme complex can exist in various substates characterized by the combination of redox centers reduced or oxidized. Since electrons on the complex reorganize on very fast time scales (< μs) relative to turnover, we compute these substates using the Boltzmann distribution as shown in Eq. 2.

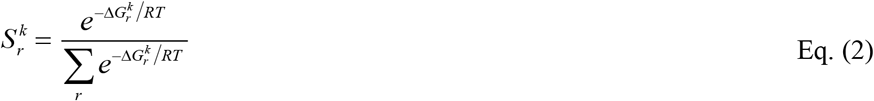

In Eq. 2, *S_r_^k^* is the fraction of redox centers *r* existing in the electronic state *k* that is reduced, and Δ*G_r_^k^* are the free energy change for each redox center *r* calculated from the linear superposition of the midpoint potentials. The redox centers *r* can consist of a single redox center or any other combination of redox centers in the complex. To calculate the free energy change for the combination of redox center, the individual free energies for the redox center reactions are summed (i.e., they are independent from each other). The number of combinations of reduced redox centers for each state is given by the binomial coefficient where *n* is the number of redox centers and *k* is the number of electrons on the complex. For details on calculating substates, see Eqs. S45-S77 in the Supplementary Material.

As mentioned above, substrates and products are assumed to bind/unbind much faster than electronic state transitions. It is assumed that binding events are independent of substates and that binding at the Q and FAD sites are independent of each other. When the enzyme is in the appropriate enzyme-substrate complex configuration, the appropriate redox reaction proceeds. Binding polynomials (BP) are used to give the fraction of the enzyme in a certain enzyme-substrate configuration. The BP for the Q_p_-site (P_Qp_) partitions this binding site into the unbound, Q-, QH_2_- and atpenin-bound fractions. The expressions 1/P_*Qp*_, [Q]/K*_Q_*/P_*Qp*_, [QH_2_]/K_*Q*_/P_*Qp*_ and [*atpenin*]/K_*A*_/P_*Qp*_, indicates the fractions of the total Q sites unbound or bound to ubiquinone, ubiquinol or atpenin, respectively. In a similar manner, the Q_d_-site can be partitioned into free and bound states as well. The BP for the FAD site (P_FAD_) partitions this binding site into the unbound, succinate-, fumarate-, malate-, malonate- and oxaloacetate-bound fractions. Similarly, the expressions 1/P_*FAD*_, [*succinate*]/K_*SUC*_/P_*FAD*_, [f*umarate*]/K_*FUM*_/P_*FAD*_, [*malate*]/K_*MAL*_/P_*FAD*_, [*malonate*]/K_*MALO*_/P_*FAD*_ and [*oxaloacetate*]/K_*OAA*_/P_*FAD*_ give the fractions of the total FAD sites bound to succinate, fumarate, malate, malonate and oxaloacetate, respectively. The BPs for the FAD- and Q-sites are given below.

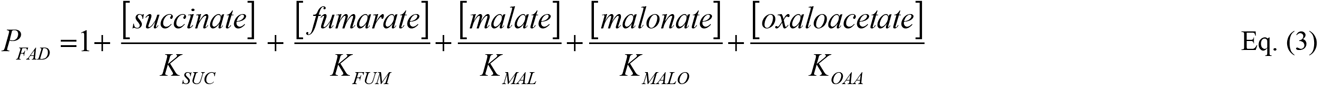

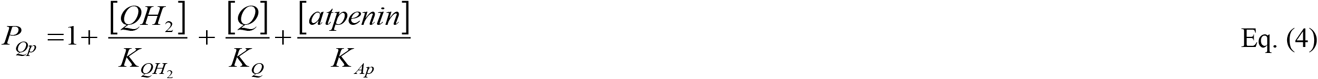

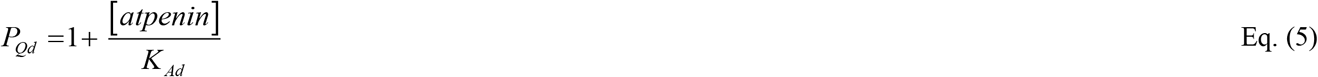

Steady-state turnover of succinate oxidation and ROS production rates are calculated using the solution of the linear equations governing the electronic transitions as shown in Eq. S92 in the Supplementary Material. The first five rows correspond to the permissible electronic state-transitions. The last row is used to set the steady-state solution to fractional occupancies (i.e., ∑_*i*_*E_i_* = 1; see the Supplementary Material for further details). The Moore-Penrose pseudo inverse is then used to calculate the unique solution to the linear system of equations [33]. The edges connecting to the electronic states shown in Fig. 1 represents the partial reactions that govern how state *i* is connected to state *j*. These reactions rates, *k_ij_*, represent molecular processes such as the reduction of FAD to FADH_2_ by succinate, oxidation of FADH^•^ or FADH_2_ by oxygen, and Q reduction at the Q site. The equations for the partial reactions are given in the Eqs. S78-S91 in the Supplementary Material. Before the enzyme can transition between electronic states, it must be in the appropriate enzyme-substrate complex configuration. For example, succinate oxidation can only occur when the FAD binding site is available for succinate to bind, and the FAD is fully oxidized. Binding polynomials (Eqs. 3-5) are used to calculate the fraction of succinate bound to the complex, and the Boltzmann distribution (Eq. 2) is used to calculate the fraction of the protein complex with a fully oxidized FAD within a given electronic state. The net steady-state rate of succinate oxidation is then computed by summing over the electronic state transition rates as shown in Eq. 6. Next, the net steady-state O_2_^•-^ production rate is computed by summing the net O_2_^•-^ production when the FADH^•^ or [*3Fe-4S*] react with oxygen as shown in Eq. 7. The steady-state H_2_O_2_ production rate is computed when the fully reduced flavin reacts with oxygen as given in Eq. 8. Lastly, the steady-state QH_2_ production rate is given by Eq. 9. Because mass and energy conservation are strictly obeyed, phenazine and TMPD reduction rates are also computed using Eq. 9. For simplicity, we assume reduction of these exogenous electron acceptors occurs in rapid, sequential one-electron steps and lump them together as a two-electron reduction reaction.

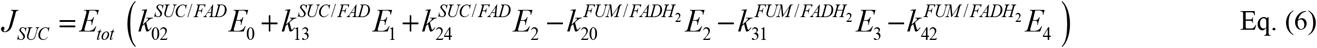

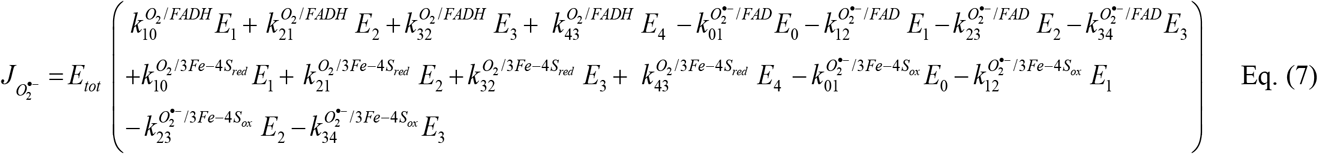

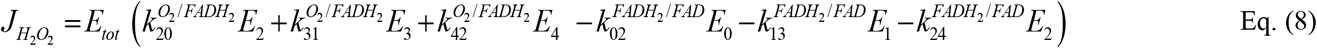

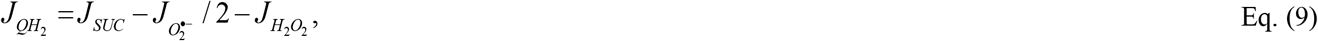

### Experimental data

The model was calibrated using a variety of literature data listed in Table 1. These data include succinate oxidation, Q reduction, O_2_^•-^ and H_2_O_2_ production rates under different experimental conditions. The data set contains information about the kinetics and ROS production by complex II necessary to identify model parameters. Most of the kinetic data on succinate oxidation is from bovine heart mitochondria [34, 35], and some are from pig heart mitochondria [36]. The ROS data set is from bovine heart mitochondria and includes both O_2_^•-^ and H_2_O_2_ production rates [11, 27]. We did not use data from Quinlan et al. as the reported data include an unknown contribution of the ROS scavenging system [21]. Both succinate oxidation and ROS data sets are obtained under a variety of experimental conditions including variations in enzyme concentrations, temperatures, pH, substrates concentrations and inhibitors concentrations. Experimental conditions are explicitly stated to simulate the experiments as faithfully as possible. However, the use of scaling factors is still necessary to account for experimental differences among the data sets such as species-dependent differences in enzyme kinetics. Even so, the values are in the acceptable range of 0.2 to 5 showing only minor adjustments were needed to fit the data. Also, in some experiments, endogenous quinone was used as the electron acceptor without any information on the redox state of Q pool [11, 27, 35]. In order to simulate these data, we relied on monotonic functions of succinate and inhibitors to predict the Q pool redox state. These equations are given in the Supplementary Material (Eqs. S1-S3).

**Table 1.** Experimental Details of Data Used for Model Fitting

| Species and Enzyme Origin | Electron acceptor | Kinetic Data | ROS Data | pH | Temp | Scaling Factors | Reference |
| --- | --- | --- | --- | --- | --- | --- | --- |
| pig heart enzyme | PMS | yes | no | 7.8 | 25 °C | 1.0 | Zeijlemaker et al. [36] |
| bovine heart enzyme | TMPD | yes | no | 7.8 | 22 °C | 0.83 | Vinogradov et al. [34]. |
| bovine heart SMP | DQ, Q<sub>10</sub> | yes | no | 7.4 | 32 °C | 4.0 | Jones and Hirst [35] |
| bovine heart SMP | Q<sub>10</sub>, PMS | yes | yes | 7.5 | 30 °C | 0.38 | Grivennikova et al. [11] |
| bovine heart SMP | Q<sub>10</sub> | yes | yes | 7.2 | 37 °C | 2.1 | Siebels and Drose [27] |

Q<sub>10</sub>, ubiquinone; DQ, decylubiquinone, PMS, phenazine methosulfate; TMPD, N,N,N',N' tetramethyl-P-phenylenediamine

### Model simulations

The model was numerically simulated using MATLAB (R2019a). The parameter optimization was performed on a Dell desktop PC (64-bit operating system and x64-based processor Intel^®^ core ^™^ i7-7700 CPU @3.60GHz and 16 GB RAM) using the Parallel Computing Toolbox. Manual determination of the parameter upper and lower bounds was followed by the gradient-based optimization algorithm fmincon. The analytic solutions for the state-steady electronic states were obtained with the MATLAB symbolic toolbox. When the standard deviation for data were not given, a standard deviation of 10% of the max value in a given data set was used during parameter estimation.

### Results and Discussion

The model fixed parameters are described in Table S1 of the Supplementary Material. These parameters consist of midpoint potentials and pKa values for the redox centers in the model obtained from the literature [11, 37, 38]. Adjustable parameters and their normalized sensitivity coefficients are given in Table 2. They were identified by simultaneously fitting all succinate oxidation and ROS data sets outlined in Table 1. The adjustable parameters include the forward rate constants of product formation and ROS production rates, the dissociation constants for substrates, products, and inhibitors, and other parameters necessary to properly simulate the environmental conditions. The data sets contain information on high and low electron flux regimes during succinate oxidization and Q reduction and low electron flux regimes during ROS production. Nearly all of the adjustable model parameters are identifiable and in a suitable range. Identifiable parameters are highly sensitive and not strongly correlated (e.g., less than 0.8) with other parameters. Normalized sensitivity coefficients are computed using Eq. S98. Correlation coefficients are calculated using Eq. S99 and are presented in a Fig. S1 of the Supplemental Material as a heat map. The normalized sensitivity coefficients give information about the contribution of each parameters to the model output. The correlation coefficients provide information on the degree the model parameters are linearly dependent with each other. The rate constant for QH_2_ production and the quinone/semiquinone midpoint potential are the only parameters strongly correlated with each other (>0.8). The top five most sensitive parameters are associated with semiquinone formation, succinate oxidation, TMPD reduction, and dicarboxylate inhibition. Not surprisingly, the quinone and quinol dissociation constants are among the least sensitive parameters. This is because there are no reliable dynamic data on quinone and quinol concentrations due to the inability to quantitatively and accurately measure these metabolites during kinetic assays.

**Table 2.**
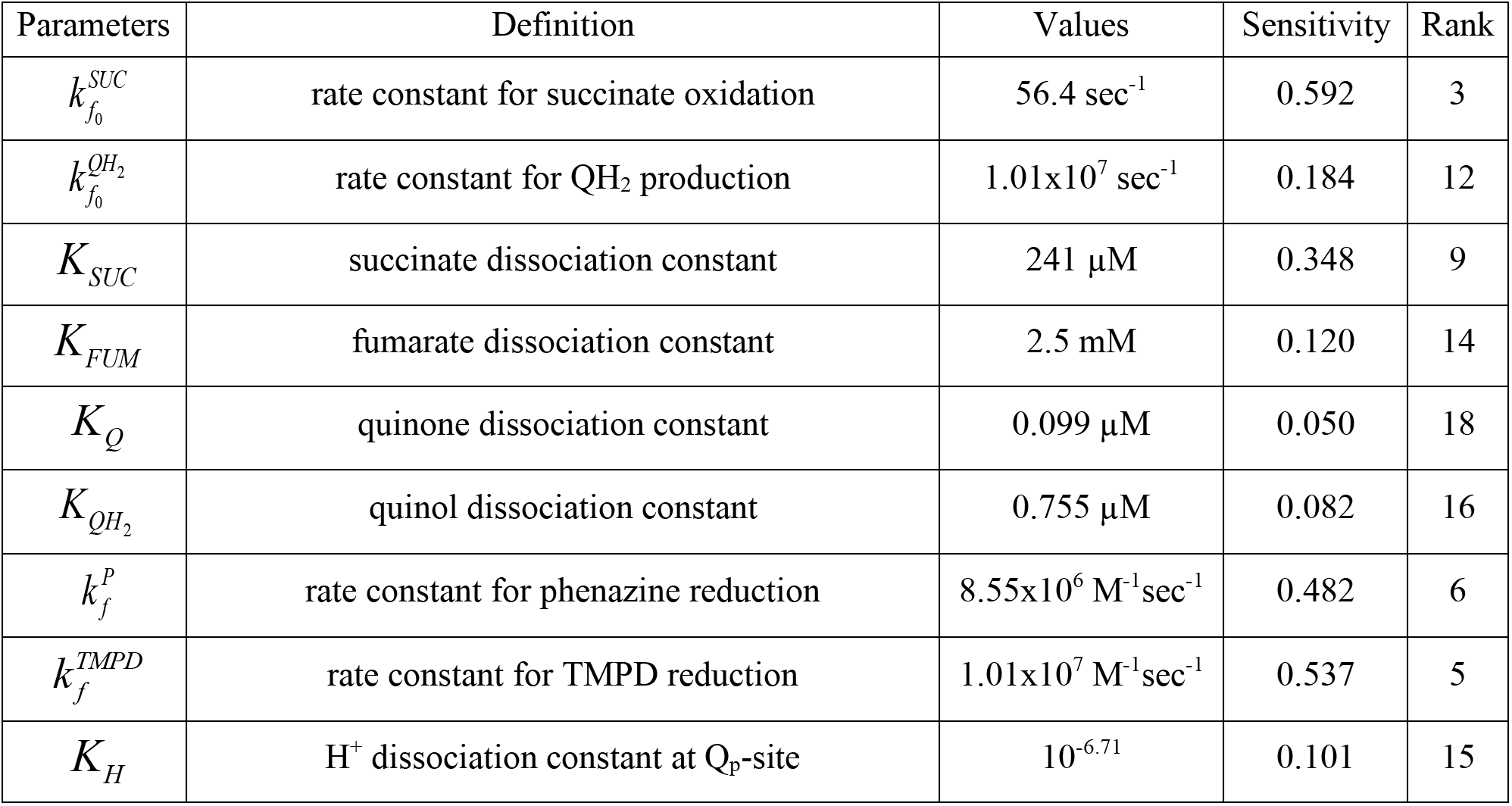
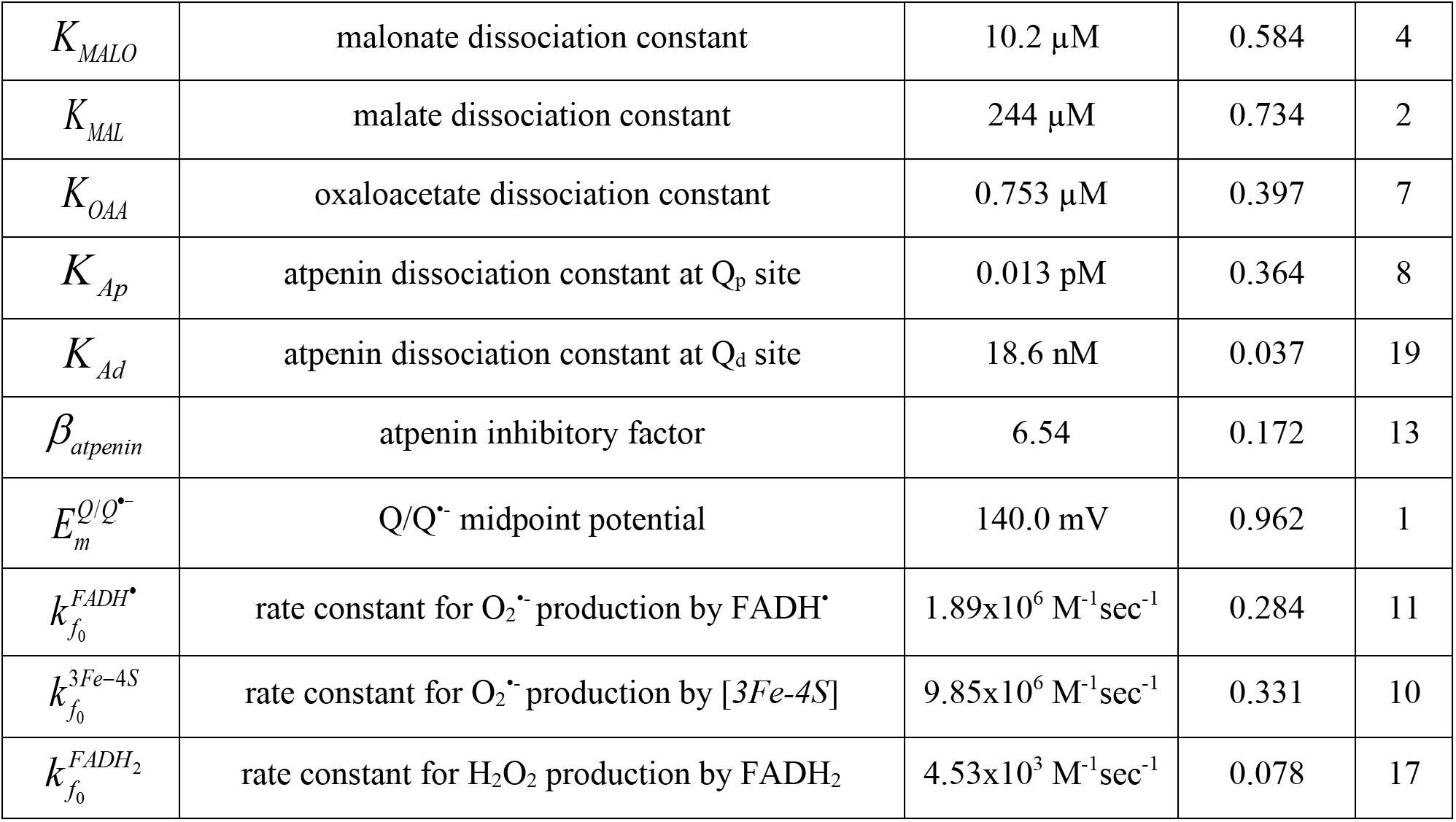
Model Adjustable Parameters

Model simulations and experimental data of succinate oxidation kinetics using PMS and TMPD as electron acceptors are shown in Fig. 2 [11, 34, 36]. Overall, the model is able to reproduce the data very well. The reduction of PMS and TMPD are assumed to obey second-order kinetics (i.e., there is no stable ES complex formed). For details concerning the reduction kinetics, see Eqs. S78-S91 in the Supplemental Material. As seen in Fig 2, the succinate oxidation rates when either PMS or TPMD is the electron acceptor are similar for a given concentration. This is due to the similar fitted second-order rate constants given in Table 2. The model is also able to reproduce pH-dependent succinate oxidation rates. Specially, at pH above 8, the reaction becomes independent of pH and precipitously drops in a pH and electron acceptor concentration dependent manner. When the acceptor concentration is high, the rate doesn’t drop until the pH drops below 7; however, when the acceptor concentration is low, the rate begins to drop below pH 8. This is because higher concentrations of electron acceptors compensate for lower concentrations of the enzyme in the right protonation state. This pH effect is ascribed to an active-site sulfhydryl group with a pK_a_ around 7 near the flavoprotein which is believed to be required for succinate oxidation [39, 40]. In the model, these results are obtained using an explicit pH-dependence for succinate oxidation at the flavin site as shown in Eq. S16.

**Figure 2.**
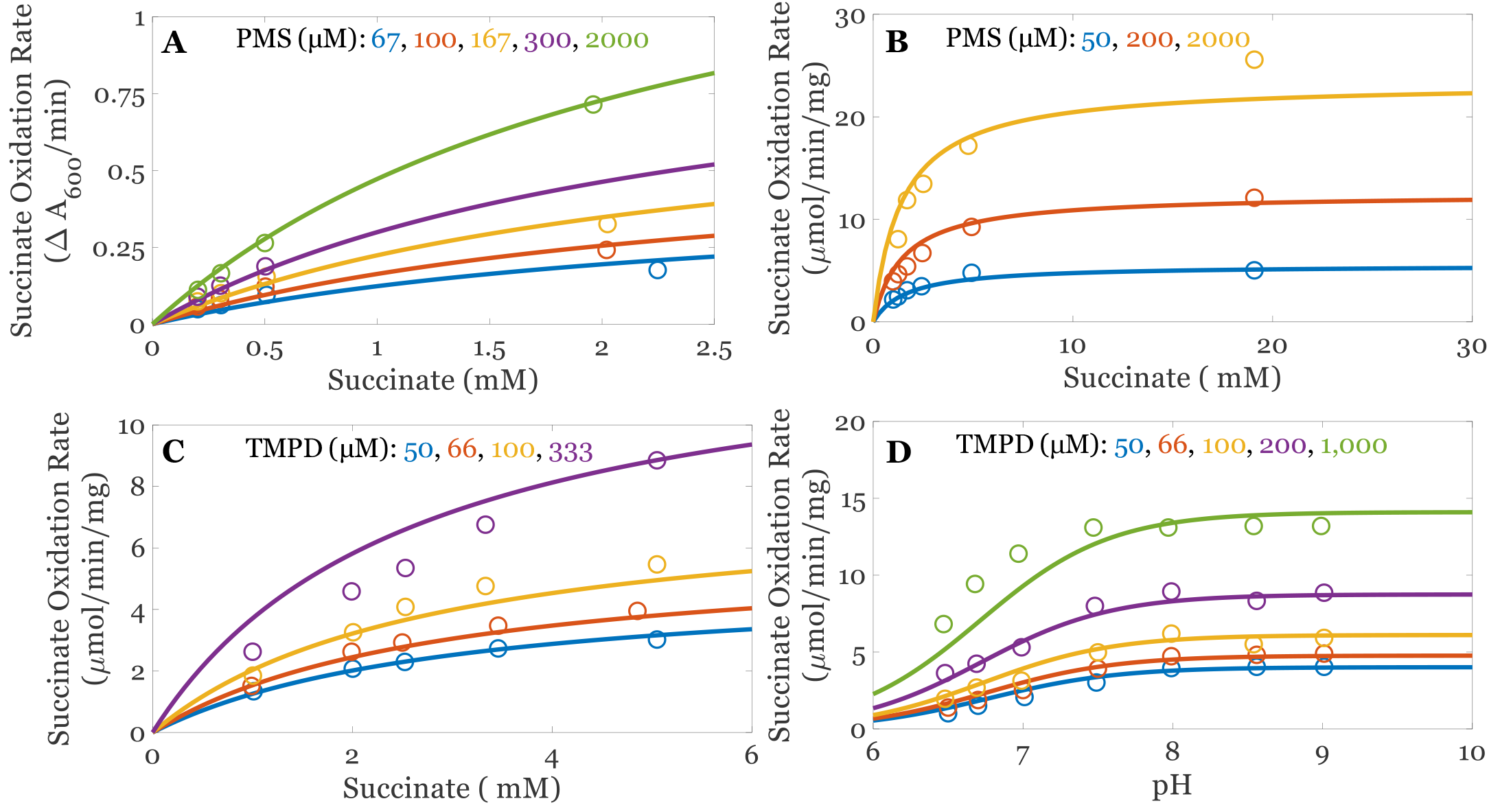
Succinate oxidation rates at varied concentrations of different electron acceptors. Model simulations (lines) are compared to experimental data (open circles) from [11, 36]. A-B) The effect of varying succinate and PMS concentrations on succinate oxidation rates. The PMS concentrations are (in μM) 67 (blue), 100 (red), 167 (yellow), 300 (purple) and 2000 (green) in panel A and 50 (blue), 200 (red) and 2000 (yellow) in panel B. For panel B, the malonate concentration was 50 μM. C) Model simulations of the effect of varying succinate and TMPD concentrations compared to the data from [34]. The TMPD concentrations (in μM) are 50 (blue), 66 (red), 100 (yellow) and 333 (purple). Malonate was present at 100 μM. D) Model simulations of the effect of varying pH and TMPD concentrations on succinate oxidation rates compared to the data from [34]. The concentrations of WB (in μM) are 50 (blue), 66 (red), 100 (yellow), 200 (purple) and 1000 (green). The succinate concentration was fixed at 100 μM.

The effects of succinate, competitive inhibitors, and quinone reductase inhibitors on succinate oxidation rates are given in Fig. 3. Data from Jones and Hirst are shown in panels A-C (top row), and data from Siebels and Drose are shown in panels D-F (bottom row). Both data sets are obtained using bovine heart SMP but differ in the enzyme and substrate concentrations. The data from Jones and Hirst were performed using 0.30 μg/mL complex II, 5 mM succinate and 100 μM decylubiquinone [35]. In Siebels and Drose, SMP was present at 0.12 mg/mL fueled with 100 μM succinate. Atpenin A5 titration leads to a significant drop in succinate oxidation rates in both data set. However, the atpenin-titration results are quantitatively different. This is likely due to the different enzyme concentrations and experimental conditions employed. For example, at an atpenin concentration of 25 nM, succinate oxidation is 93% inhibited in the Jones and Hirst data set (panel B), but at 30 nM it is only inhibited by 65% in the Siebels and Drose dataset (panel D). To fit these disparities, the model fitting results in a compromise where it underestimates the atpenin-dependent inhibition for the Jones and Hirst dataset and overestimates it for the Siebels and Drose dataset. In order to fit these atpenin inhibitory data, two independent atpenin binding sites were necessary. The first binding site is the canonical Q_p_ site which competes with Q and QH_2_ binding. In addition to this binding site, a second site that inhibited both Q reduction and succinate oxidation was required. This site is the Q_d_ site and possess a much lower affinity for atpenin than the Q_p_ site. This model result is supported by crystal structures that show occupancy of the Q_d_ site at high atpenin concentrations [5, 41]. Titrating succinate-binding competitors malonate, malate, and oxaloacetate similarly inhibits succinate oxidation rates. However, the concentration required to inhibit succinate oxidation is different for each inhibitor. Oxaloacetate is the most potent inhibitor, followed by malonate and malate. These results coincide with the fitted dissociation constants shown in Table 2.

**Figure 3.**
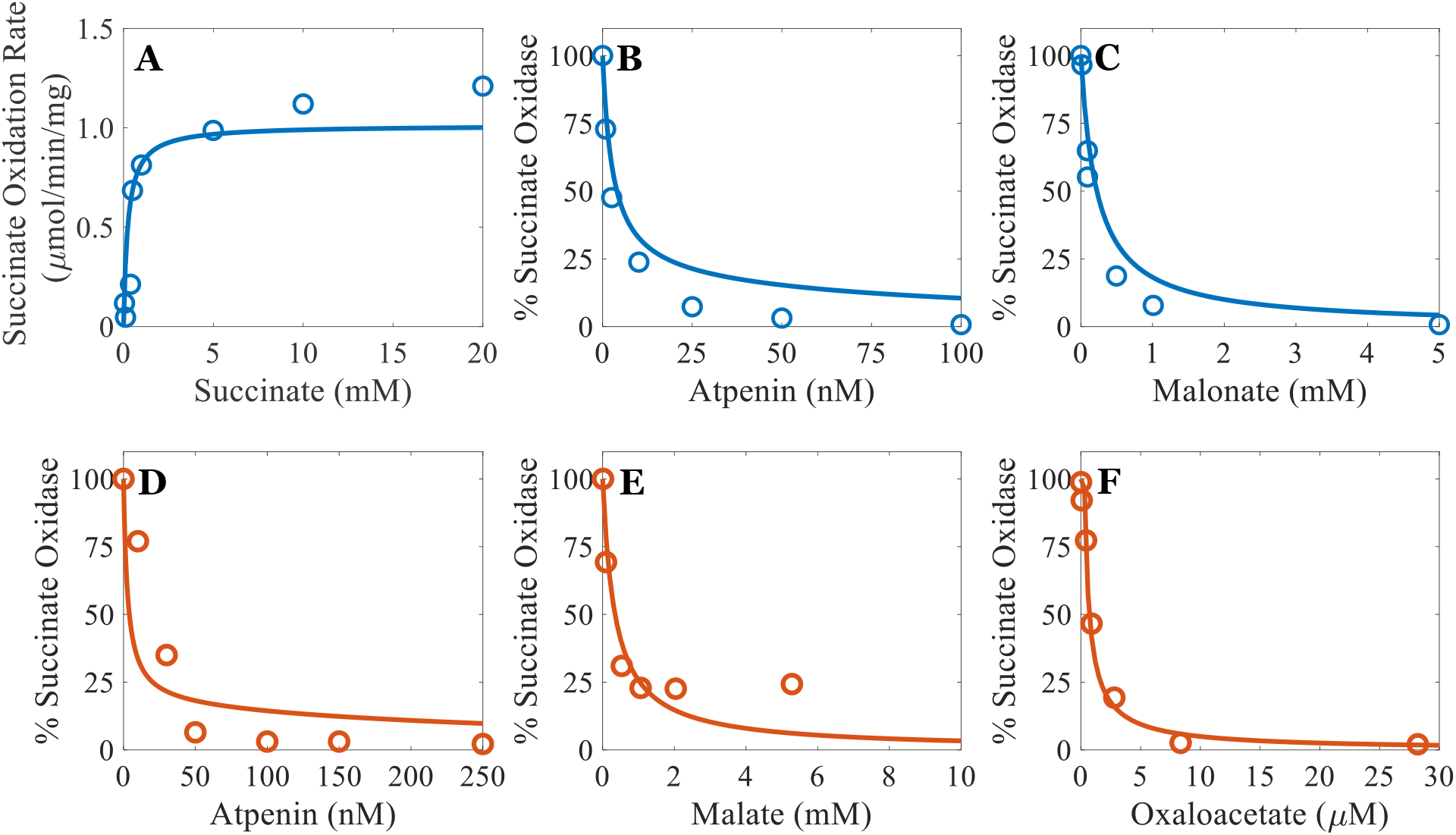
Succinate oxidation kinetics in the presence of SDH inhibitors using bovine heart SMP. Model simulations (lines) are compared to experimental data (open circles). Experimental data are from Jones and Hirst (A-C) and Siebels and Drose (D-F). Data from Jones and Hirst are obtained using 30 μg/mL solubilized membranes, 5 mM succinate and 100 μM decylubiquinone. A) Succinate oxidation rate saturates near 5 mM succinate with an apparent K_M_ of 2 mM. B-C) Relative succinate oxidation rates are significantly inhibited with increasing concentrations of atpenin (0-100 nM) and malonate (0-5 mM). E-F) Data from Siebels and Drose are obtained using 0.12 mg/mL SMP and 100 μM succinate. Relative succinate oxidation rates are significantly inhibited with increasing concentrations of atpenin (0-250 nM), malate (0-10 mM) and oxaloacetate (0-30 μM).

ROS production by complex II displays a biphasic dependency on succinate concentrations as shown in Fig. 4. At low concentrations, ROS production is stimulated. However, ROS production is suppressed at higher concentrations. ROS production from the FAD site of complex II requires two conditions. First, there must be a source of electrons to reduce FAD such as succinate or quinol. Second, the FAD site must be available to interact with oxygen. ROS production from the [*3Fe-4S*] ISC has similar requirements where the ISC binding pocket needs to be reduced and unoccupied. Dicarboxylates such as succinate, malate, and oxaloacetate inhibit ROS production at the FAD site by physically blocking the reaction of between FAD and oxygen. Likewise, molecules that bind to the Q_p_ site can also physically block the ISC from reacting with oxygen. However, ROS production from the ISC is independent of dicarboxylates but does depend on the present of molecules that bind to the Q site. In the experiments by Grivennikova *et al.* [11], peak ROS production occurs at around 50 μM succinate as shown in Fig. 4A. However, in the experiments from Siebels and Drose [27], the peak is shifted to approximately 150 μM. The primary difference between these two studies is the presence of myxothiazol in the Grivennikova *et al.* experiments and atpenin in the Siebels and Drose experiments. With myxothiazol, the Q pool is expected to be fully reduced and capable of reducing complex II via QH_2_ oxidation at the Q_p_ site. This causes a higher proportion of the enzyme to be reduced and capable of producing ROS at lower succinate concentrations. In contrast, the only reductant for complex II when atpenin is present is succinate. Therefore, it takes a higher concentration of succinate to reduce enough of the enzyme to reach peak ROS production rates. However, as stated above, succinate also blocks ROS production from the FAD site, so the ability of succinate to drive ROS production is limited.

**Figure 4.**
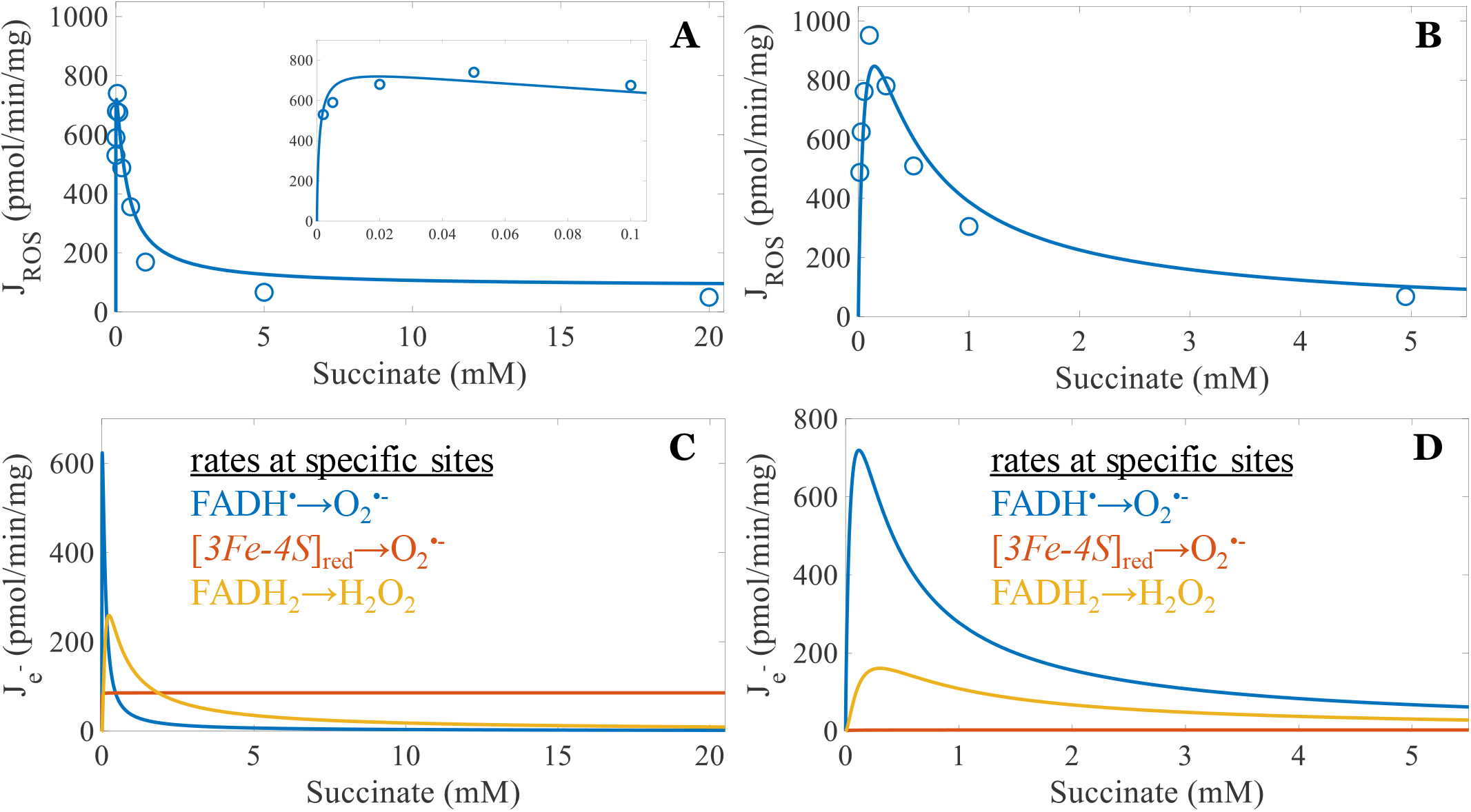
Maximum ROS produced by complex II occurs at sub-millimolar concentrations of succinate. Model simulations (lines) are compared to experimental data (open circles) from bovine heart SMP. A) Experimental J_H2O2_ are obtained in the presence of 10 μM rotenone, 1.6 μM myxothiazol, 2 units/mL horseradish peroxidase (HrP) and 6 units/mL bovine superoxide dismutase (SOD) [11]. B) Experimental J_H2O2_ are obtained in the presence of 50 nM atpenin [27]. C) Model simulations of site-specific ROS production rates for the conditions shown in panel A). C) Model simulations of site-specific ROS production rates for the conditions shown in panel B). In both panels C) and D), rates are given with respect to the same electron equivalents as in A) and B).

The difference in total and site-specific ROS production rates in the presence of stigmatellin or atpenin as the succinate concentration is increased is shown in Fig. 5. Stigmatellin induces the largest rate of ROS production from complex II. With stigmatellin present, the Q pool is fully reduced and quinol oxidation at the Q_p_ site serves as an additional source of electrons for free radical production. So turnover at both electron input and output (reverse) of the enzyme complex results in higher ROS production rates. Superoxide production from the FAD at low succinate concentrations is highest under these conditions. As the succinate concentration increases, hydrogen peroxide production from the FAD exceeds the superoxide rate from the FAD. But as the succinate concentration further increases, the ROS production rate from the FAD decreases due to succinate binding to the FAD, making it unavailable to interact of oxygen. As a result, ROS from the [*3Fe-4S*] ISC becomes the dominant source. However, when atpenin is present, electron transfer from the ISC to quinone or oxygen is blocked [41, 42]. Under these conditions, superoxide production from the FAD is the dominate source of ROS, and the ISC produces no ROS. In addition, the enzyme is more oxidized compared to when stigmatellin is present, so the amount of enzyme with a fully reduced flavin needed to produce hydrogen peroxide is lower. Thus, the rate of hydrogen peroxide production is lower. In the presence of either inhibitor, the FAD is the major source of ROS.

**Figure 5.**
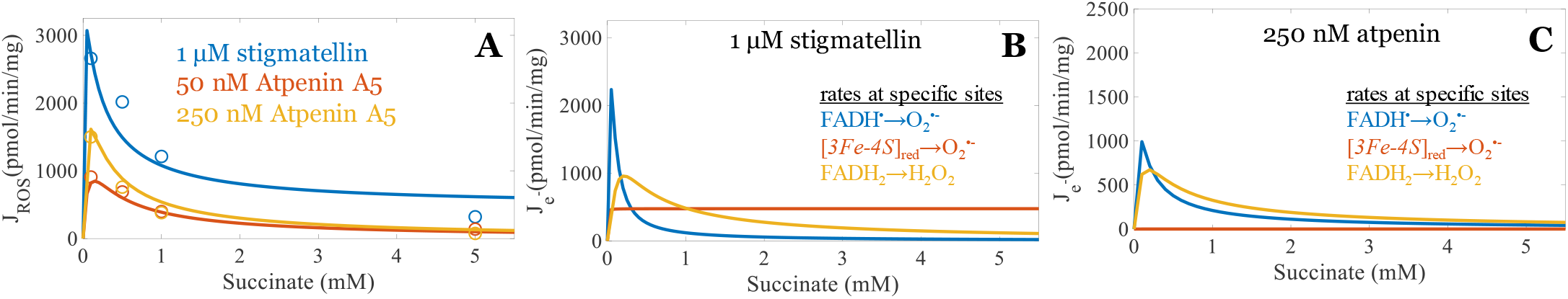
Succinate-dependent ROS production of SMPs in the presence of stigmatellin or atpenin. A) Model simulations (lines) are compared to experimental data (open circles) from Siebels and Drose (21). B) Site-specific ROS production rates when 1 μM stigmatellin is present. C) Site-specific ROS production rates when 250 nM atpenin is present. Site-specific rates are given with respect to the same electron equivalents as in A).

The effects of titrating atpenin and non-complex II dicarboxylate substrates on ROS production rate are presented in Fig. 6. Increasing atpenin concentration leads to increased ROS production rate and approaches maximum at 250 nM. In the presence of 250 nM atpenin, increasing malate or oxaloacetate concentration causes the ROS production rate to significantly decrease. Malate concentrations in the mM range lead to a significant drop in ROS production rates. Oxaloacetate is much more potent and able to completely inhibit ROS production after 20 μM. These data further corroborate that the availability of the FAD site for oxygen binding is essential for ROS formation by complex II. As more FAD sites are occupied by dicarboxylates, less ROS is produced despite the optimal atpenin concentration. These results are supported by prior work demonstrating that ROS are only produced from the FAD site when the dicarboxylate site is unoccupied [8, 9]. As these TCA cycle dicarboxylates regulate complex II turnover, it is fortunate that they also do not lead to excess ROS production like the Q site inhibitor atpenin. This would necessarily lead to oxidative stress when these dicarboxylates accumulate under conditions such as IR injury.

**Figure 6.**
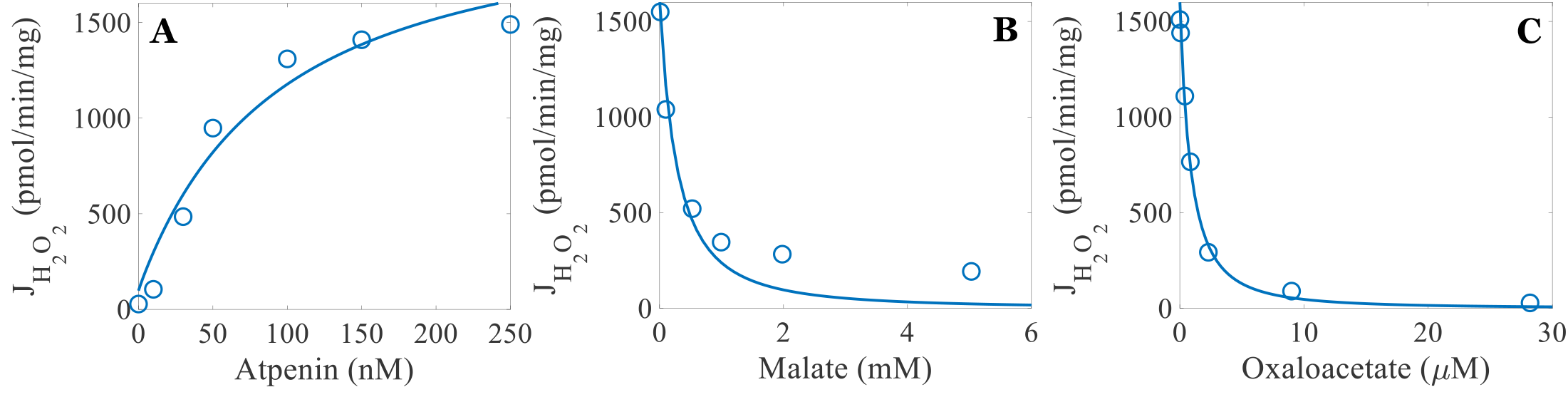
Effects of titrating atpenin and non-complex II dicarboxylate substrates on ROS production by complex II. Model simulations (lines) are compared to experimental data (open circles) for each panel (21). Experimental data are obtained from bovine heart SMP. A) Rate of H_2_O_2_ production is increased at increasing concentrations of atpenin (0-250 nM), a potent and specific inhibitor of quinone reductase activity. B-C) In the presence of 250 nM atpenin, ROS production rates are decreased as the concentrations of malate and oxaloacetate is increased. Malate and oxaloacetate are competitive inhibitors of succinate binding to the complex.

Titrating fumarate in the presence of varying succinate concentrations results in drop in superoxide and total ROS production rates as shown in Fig. 7. When atpenin was absent from the experiments, it was assumed that the quinone/quinol redox couple was in equilibrium with the fumarate/succinate couple. Therefore, the equilibrium relationship between the free energies associated with the fumarate/succinate and quinone/quinol couples were used to calculate the quinone and quinol concentrations. When aptenin was present, the Q pool redox state predictor functions (Eq. S1) was used to calculate the quinone and quinol concentrations. At a given fumarate to succinate ratio, the ROS production rate is higher in the presence of complex III inhibitors alone compared to when atpenin is included. Complex III inhibitors, stigmatellin and myxothiazol, bind to the Q_p_ site of complex III and prevent quinol oxidation. Although they bind to two different sites in the Q_p_-site pocket, the net result is a Q pool that is nearly fully reduced (Eq. S2). And without atpenin present, significant turnover at the Q_p_ site of complex II, in addition to turnover at the FAD site, leads to more reduced FAD fraction and hence more ROS production. While atpenin stimulates ROS production from the FAD site, the rate is one third to one half of that with stigmatellin alone. This difference highlights the importance of the Q_p_ site as a source of electrons via quinol oxidation for the ROS production at the FAD site. While the [*3Fe-4S*] ISC and FAD sites can all produce ROS, both experimental data and model simulations suggest that most of ROS comes from the reduced flavin under these conditions. In addition, the model simulations show that hydrogen peroxide production rates are significantly suppressed in the presence of fumarate. These finding further corroborates the aforementioned conditions that lead to significant ROS production by complex II requires the flavin to be reduced and available to interact with oxygen.

**Figure 7.**
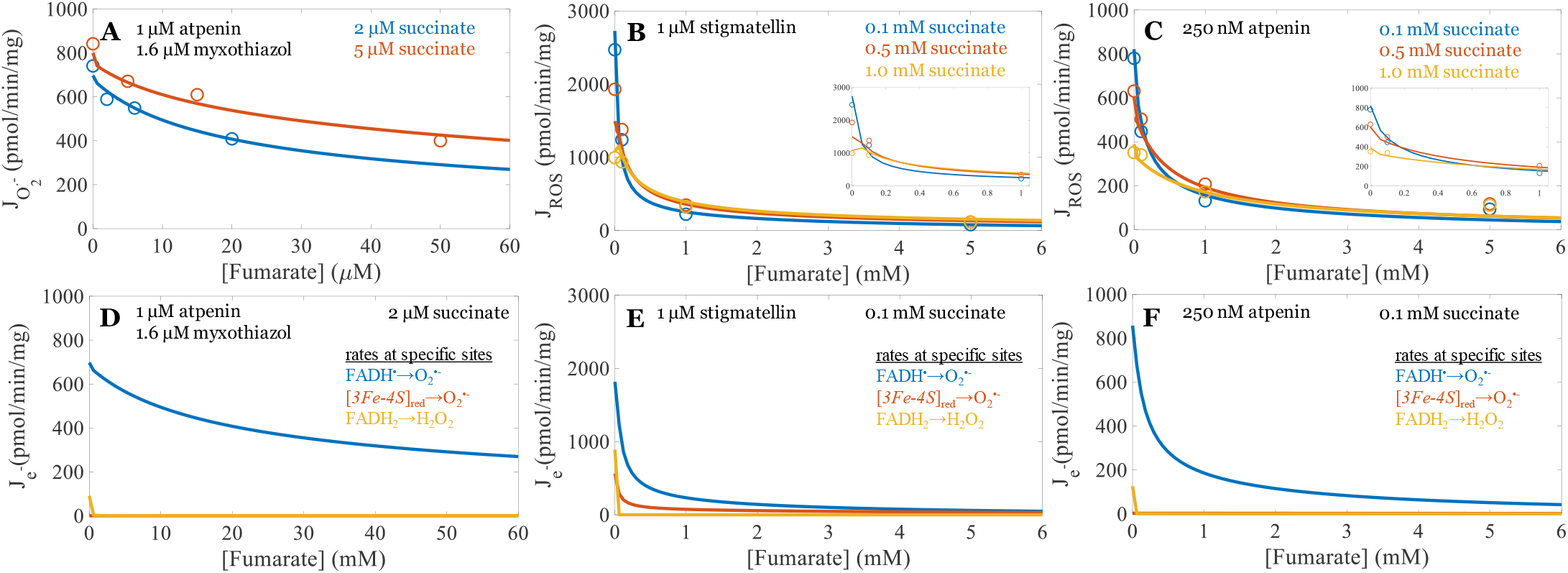
Effects of varying fumarate/succinate ratios on atpenin and complex III inhibitor induced ROS production. Model simulations (lines) are compared with experimental data (open circles). A) Data are obtained from Grivennikova et al. (22). Bovine heart SMP (0.1 mg/mL) are prepared in the presence of 1.6 μM myxothiazol and 10 μM rotenone. Atpenin is added to the final concentration of 1 μM. B-C) Data are from Siebels and Drose (21). Bovine heart SMP (0.12 mg/mL) are prepared without myxothiazol or rotenone. B) Stigmatellin is added to the final concentration of 1 μM C). Atpenin is added to the final concentration of 50 nM. In all conditions, the increasing the fumarate concentration leads to a decrease in ROS production rates. D-F) Site-specific ROS production rates corresponding to their respective panels above. Site-specific rates are given with respect to the same electron equivalents as in above panels.

The model simulations show distinct pH-dependent ROS profiles for stigmatellin and atpenin. In the presence of stigmatellin, both ROS species increase as pH is increased as shown in Fig. 8. This is due to the pH-dependent reaction occurring at the Q_p_ site of the complex. As pH becomes more alkaline, the quinol oxidation becomes more thermodynamically favorable from mass action (see Eq. S25). As a result, the enzyme oxidation status becomes more reduced, and the fraction of reduced FAD is significantly elevated. In addition to this mechanism, ROS production from the FAD site was best fit when the FAD radical or fully reduced FAD cofactor was deprotonated as shown in Eqs. S18-20. In contrast, this rise in ROS production as the conditions become more alkaline are not observed when atpenin is present. With atpenin, quinol oxidation at the Q_p_ site is inhibited, so the elevated fraction of FAD that is reduced is not present. Therefore, ROS production is overall lower under this condition despite the thermodynamic favorability of ROS production from the FAD site in alkaline conditions. Prior studies have argued that the FAD site is able to produce both superoxide and hydrogen peroxide [8, 9, 21, 24, 43]. However, which is the dominant species and under which conditions is a standing question. In the study by Siebels and Drose [27], it was determined that hydrogen peroxide was the primary ROS species produced from the FAD site. We suspect their prep contained low, yet significant amounts, of superoxide dismutase contamination as described by Grivennikova et al. (11). In support of this, the simulations shown in Fig. 8 reveal that under both stigmatellin and atpenin induced ROS production, complex II mainly produces ROS as superoxide form the FAD site. These findings are supported by several studies which used rat skeletal muscle mitochondria to show that most ROS originating from the FAD site is superoxide [21, 26]. In addition, a different study found that the vast majority of total ROS produced by *E. coli* complex II is superoxide [43]. These findings and the model simulation results strongly point to superoxide as the major free radical species originating from the FAD site.

**Figure 8.**
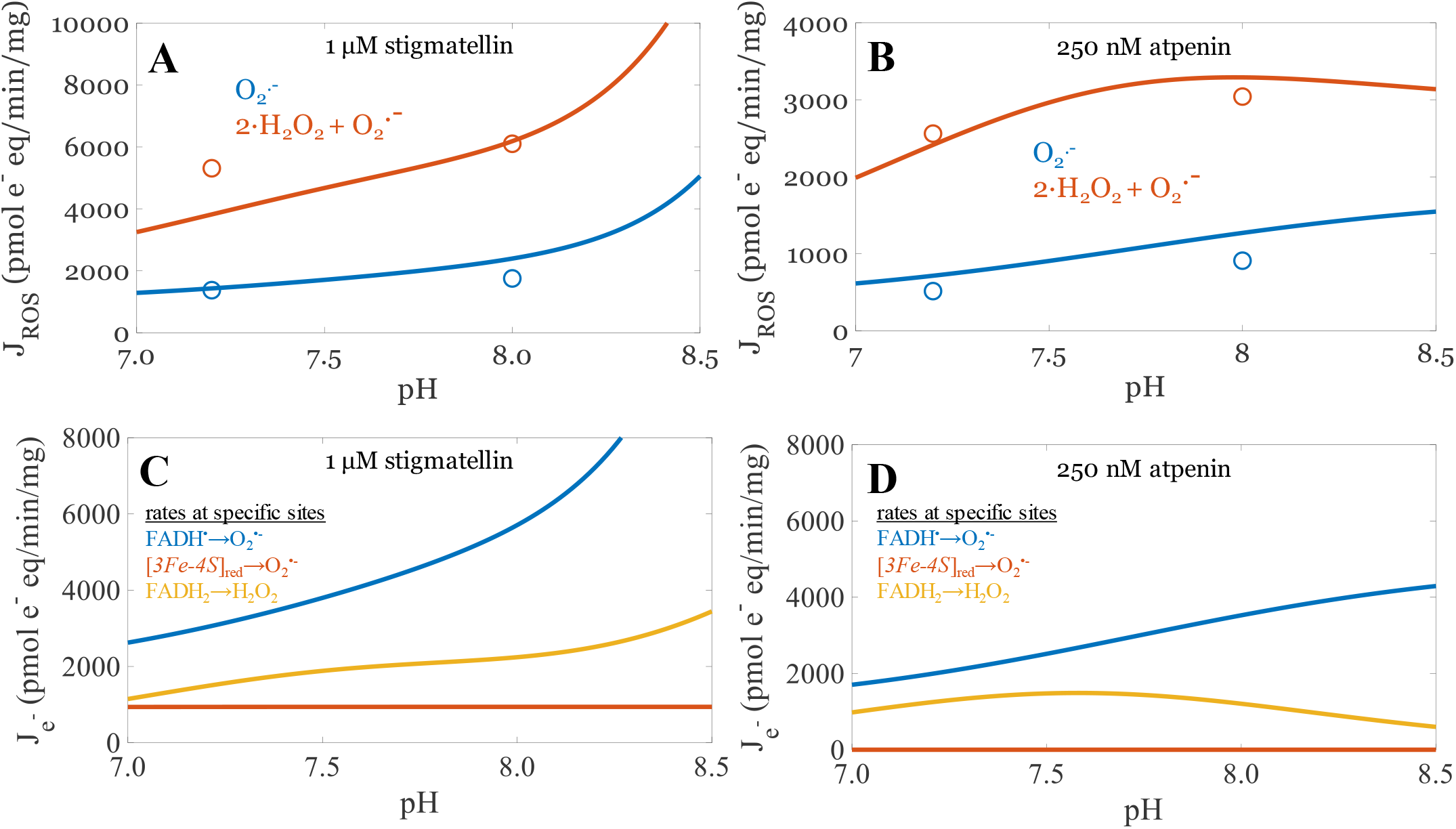
The effects of pH on ROS production rates in the presence of inhibitors. In A), the stigmatellin concentration was 1 μM. In B), the atpenin concentration was 250 nM. Model simulations (lines) are compared to experimental data (open circles) from Siebels and Drose (21). Experimental data show O_2_^•-^ in blue and total ROS (H_2_O_2_ + O_2_^•-^) in orange using bovine heart SMP. Malonate of 1.5 mM is added in both experiments. As discussed in Grivennkova et al. [11], we assumed the presence of a small but significant amount of contaminating superoxide dismutase in these experiments. Based on model analysis, this amounts to an approximate 65% underestimation of the superoxide production rate in the Siebels and Drose experiments. This factor was explicitly included when simulating these data. C-D) Site-specific ROS production rates corresponding to their respective panels above. Site-specific rates are given with respect to the same electron equivalents as in above panels.

The model was corroborated using the experimental data from Grivennikova et al. [44]. In this study, ROS production rates from SMPs were measured with a variety of fuel substrates, inhibitors, and oxygen concentrations. From these data, the complex II specific data were selected and used to test model validity. As shown in Fig. 9, the model is capable of reproducing the experimental data with high fidelity without tuning the parameters. It reproduces the linear relationship between ROS production and oxygen concentration in Fig. 9A. In addition, the model matches the kinetic rate of succinate oxidase under the conditions studied in Fig. 9B. The quinol concentration and complex II electronic state fractions are shown as predictions during the conditions in Fig. 9B and 9C, respectively. Furthermore, the contribution of complex II to the total ROS detected during this experiment is shown in Fig. 9D. Comparing this value to the total value Grivennikova et al. measured during this experiment reveals that complex II contributes approximately 2% to the total ROS produced by SMPs during uninhibited metabolism.

**Figure 9.**
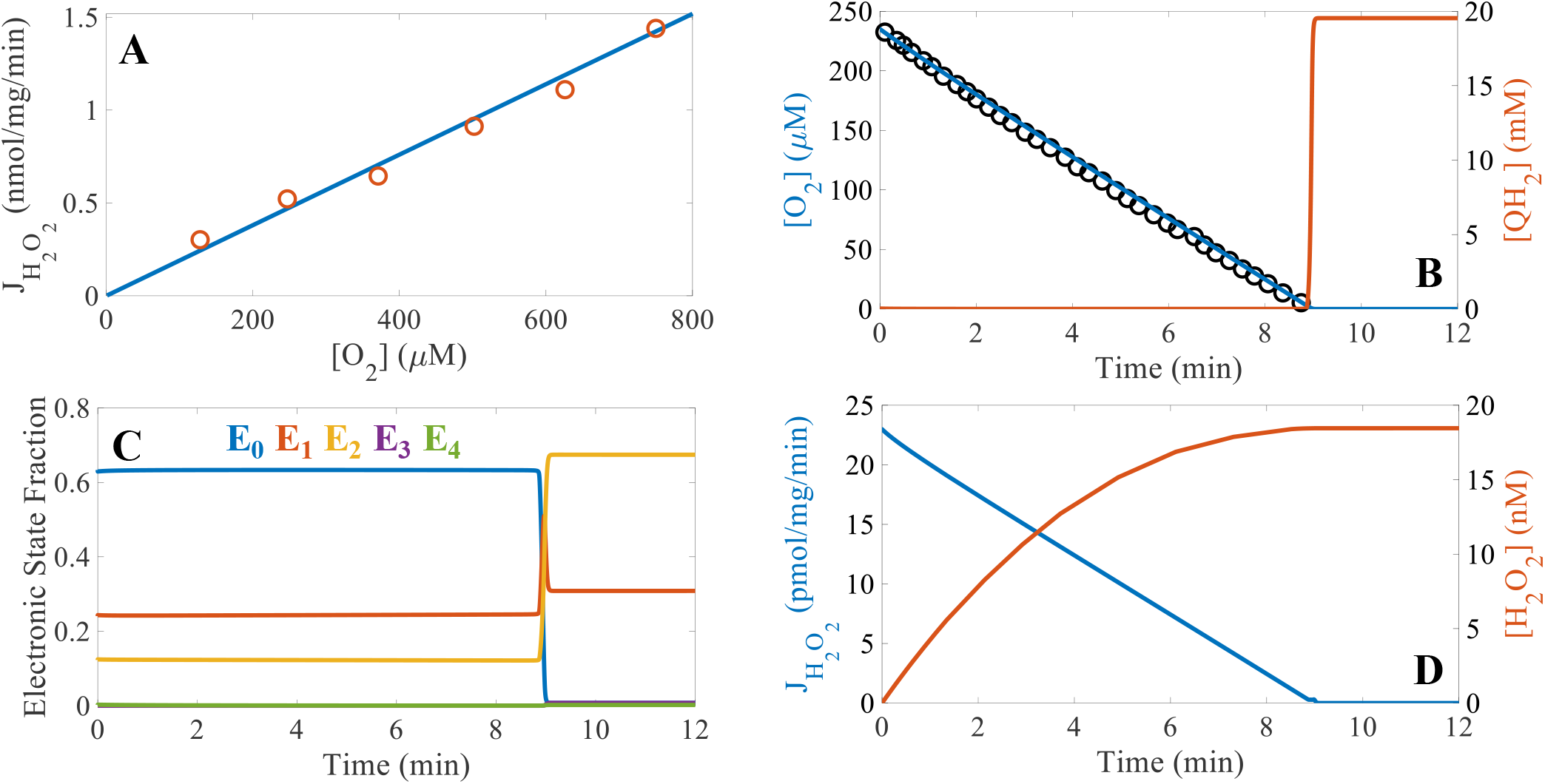
Model corroboration of ROS production and complex II kinetics. The data are from Grivennikova et al. [44]. The enzyme concentration for these simulations was 0.85 nM in A) and 20 nM in B-D). The ODE system used to simulate oxygen concentration dynamics is given in Eqs. S100-106 in the Supplemental Material.

Model prediction of the rate of succinate oxidation and ROS production as a function of the succinate/fumarate and quinol/quinone ratios in the absence of inhibitors is shown in Fig. 10. The succinate oxidation rate is maximum when the succinate/fumarate ratio is high and the quinol/quinone ratio is low. The turnover in the forward direction predicted by the model is in the range previously determined by other groups [45–47]. The model also predicts that enzyme’s turnover in the reverse direction is approximately double that of the forward direction. This has a profound effect on the enzyme behavior during pathological conditions such as ischemia whereby this enzyme is speculated to be responsible for the significant accumulation of succinate during ischemia [15, 16]. In contrast to the inhibitor-based experiments described above, the model predicts that ROS production from the [*3Fe-4S*] ISC is the primary source of free radicals when no inhibitors are present. This is not surprising considering under inhibitor free conditions, the enzyme is mostly in the E_0_, E_1_, and E_2_ electronic states. In these states, electrons on the complex have a higher probability residing on the ISCs instead of the FAD. Therefore, ROS production form the FAD of complex II is not expected to significantly contribute to the total ROS produced by functional, respiring mitochondria. In addition, hydrogen peroxide production under these conditions is negligible. Hydrogen peroxide production from complex II only occurs when the electronic state E_4_ is elevated when fumarate is zero or in the low micromolar range and where the probability of a fully reduced FAD is high (Fig. 1C) which only happens when inhibitors are present.

**Figure 10.**
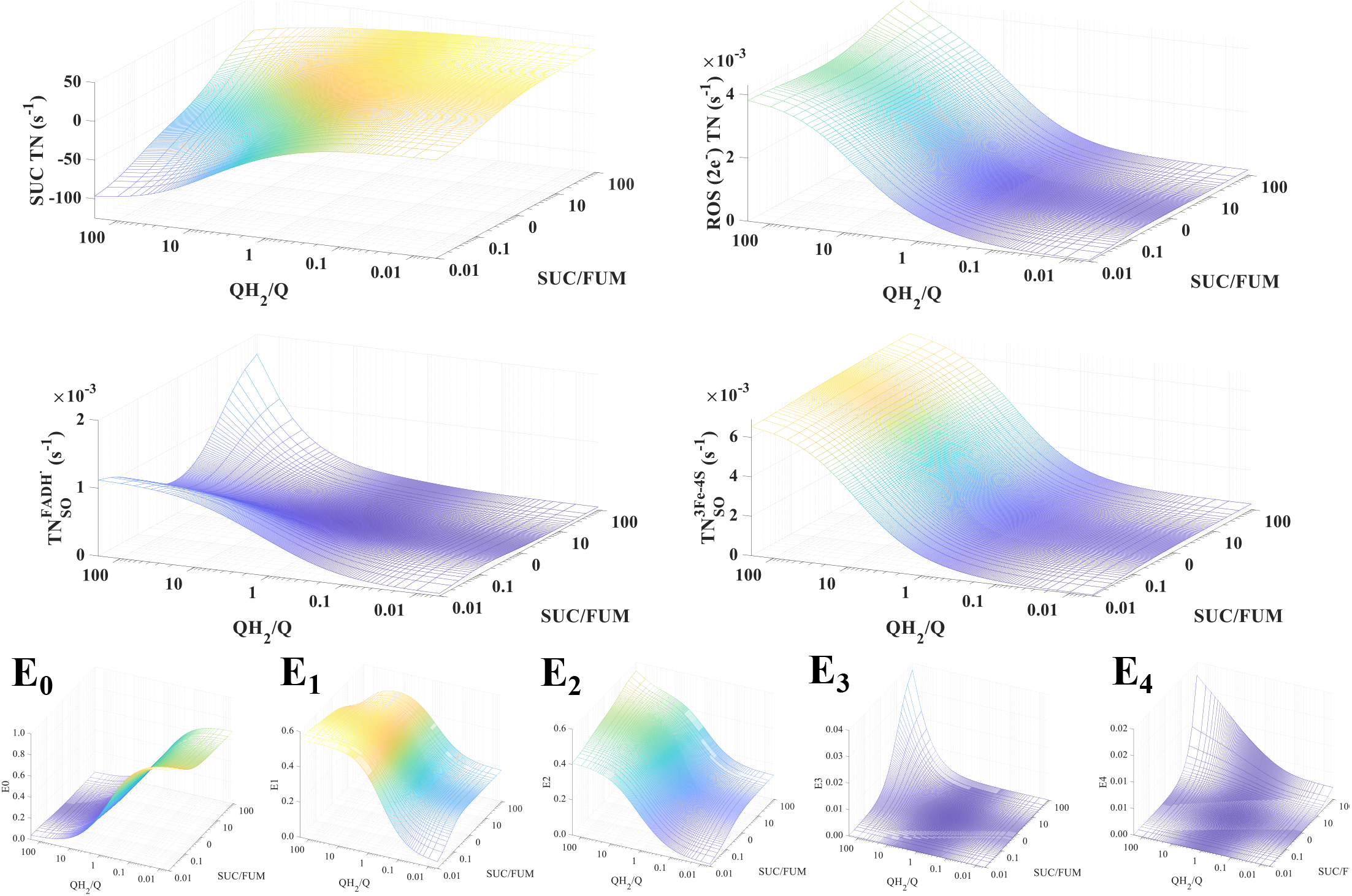
Model simulations of succinate and ROS turnover of complex II as a function of succinate/fumarate and QH_2_/Q ratios. Steady-state electronic states for these conditions is given in bottom row. The simulations were conducted at 37 °C with a total Q pool of 20 mM, with succinate + fumarate = 20 mM, [O_2_] equal to 20 μM, and at pH 7.

In complex II, the two major sites for ROS production are proposed to be the covalently bound FAD and the Q site [8, 9, 21, 24, 43]. However, which site is the dominant source, what ROS species is formed, and what conditions are conducive for ROS production has not been quantitatively addressed until this study. Using *S. cerevisiae* as an experimental model, several groups determined that the Q site is a strong contender for the site of ROS production [10, 48]. Additional studies on the *E. coli* support the Q site origin hypothesis [43, 49]. However, studies using *A. suum* argued that both the FAD and the Q sites can produce ROS [50]. In contrast, other studies reported that only the FAD site produce ROS [9, 43]. In mammalian complex II, studies have shown ROS is produced when the enzyme is fueled by succinate only after the Q site was inhibited [8, 21]. But neither study demonstrably showed whether superoxide [11] or hydrogen peroxide [27] is the dominate ROS species formed. In contrast, it has been determined that complex II from *E. coli* mainly produces superoxide from the FAD site [43]. This and other studies concluded that quinol fumarate oxidoreductases generate superoxide from the fully reduced FAD [4, 43]. However, the mechanism underlying the ROS production was not elucidated. To answer these questions, we developed and analyzed a computational model of complex II that quantiatively describes the necessary conditions for ROS production, the contribution of each site responsible for ROS production, and the factors that control how much ROS is produced by complex II.

In summary, a kinetic model of complex II that is also capable of simulating ROS production rates under a variety of conditions is developed, corroborated, and analyzed. The types of ROS production from complex II, the sites of ROS productions, and the conditions necessary for ROS production was identified using this computational model. The model includes the major redox centers in the four subunits of complex II which include the FAD site, dicarboxylate binding site, ISC centers, and two quinone binding sites. Model analysis reveals that ROS production by complex II can originate either from the FAD site or the [*3Fe-4S*] ISC. Our study resolves a critical issue pertaining to the origin of ROS production from complex II and shows that the primary species formed is superoxide. During physiogical conditions, the [*3Fe-4S*] ISC is the primary source; however, the FAD can become a major source during pathalogical conditions. Hydrogen peroxide is only produced in appreciable quantities by the enzyme when inhibited and in the absence offumarate. As a result, this model is an ideal choice for integration into large-scale models of mitochondrial metabolism to study ROS production from the ETS during physiological and pathophysiological conditions.

## Acknowledgments

This work was supported by NIH grant R00-HL121160.

## References

[1] R. Dhingra, L.A. Kirshenbaum, Succinate dehydrogenase/complex II activity obligatorily links mitochondrial reserve respiratory capacity to cell survival in cardiac myocytes, Cell Death Dis 6 (2015) e1956.

[2] M.S. Hwang, J. Rohlena, L.F. Dong, J. Neuzil, S. Grimm, Powerhouse down: Complex II dissociation in the respiratory chain, Mitochondrion 19 Pt A (2014) 20–8.

[3] F. Sun, X. Huo, Y. Zhai, A. Wang, J. Xu, D. Su, M. Bartlam, Z. Rao, Crystal structure of mitochondrial respiratory membrane protein complex II, Cell 121(7) (2005) 1043–57.

[4] V. Yankovskaya, R. Horsefield, S. Tornroth, C. Luna-Chavez, H. Miyoshi, C. Leger, B. Byrne, G. Cecchini, S. Iwata, Architecture of succinate dehydrogenase and reactive oxygen species generation, Science 299(5607) (2003) 700–4.

[5] E. Maklashina, G. Cecchini, The quinone-binding and catalytic site of complex II, Biochim Biophys Acta 1797(12) (2010) 1877–82.

[6] F.J. Ruzicka, H. Beinert, K.L. Schepler, W.R. Dunham, R.H. Sands, Interaction of ubisemiquinone with a paramagnetic component in heart tissue, Proc Natl Acad Sci U S A 72(8) (1975) 2886–90.

[7] J.C. Salerno, T. Ohnishi, Studies on the stabilized ubisemiquinone species in the succinate-cytochrome c reductase segment of the intact mitochondrial membrane system, Biochem J 192(3) (1980) 769–81.

[8] C.L. Quinlan, I.V. Perevoschikova, R.L. Goncalves, M. Hey-Mogensen, M.D. Brand, The determination and analysis of site-specific rates of mitochondrial reactive oxygen species production, Methods Enzymol 526 (2013) 189–217.

[9] M.D. Brand, Mitochondrial generation of superoxide and hydrogen peroxide as the source of mitochondrial redox signaling, Free Radic Biol Med 100 (2016) 14–31.

[10] S.S. Szeto, S.N. Reinke, B.D. Sykes, B.D. Lemire, Ubiquinone-binding site mutations in the Saccharomyces cerevisiae succinate dehydrogenase generate superoxide and lead to the accumulation of succinate, J Biol Chem 282(37) (2007) 27518–26.

[11] V.G. Grivennikova, V.S. Kozlovsky, A.D. Vinogradov, Respiratory complex II: ROS production and the kinetics of ubiquinone reduction, Biochim Biophys Acta Bioenerg 1858(2) (2017) 109–117.

[12] L.A. Sena, N.S. Chandel, Physiological roles of mitochondrial reactive oxygen species, Mol Cell 48(2) (2012) 158–67.

[13] F. Toren, Oxygen radicals and signaling., Current Opinion in Cell Biology 10 (1998) 248–253.

[14] J.N. Bazil, Analysis of a Functional Dimer Model of Ubiquinol Cytochrome c Oxidoreductase, Biophys J 113(7) (2017) 1599–1612.

[15] E.T. Chouchani, V.R. Pell, E. Gaude, D. Aksentijevic, S.Y. Sundier, E.L. Robb, A. Logan, S.M. Nadtochiy, E.N.J. Ord, A.C. Smith, F. Eyassu, R. Shirley, C.H. Hu, A.J. Dare, A.M. James, S. Rogatti, R.C. Hartley, S. Eaton, A.S.H. Costa, P.S. Brookes, S.M. Davidson, M.R. Duchen, K. Saeb-Parsy, M.J. Shattock, A.J. Robinson, L.M. Work, C. Frezza, T. Krieg, M.P. Murphy, Ischaemic accumulation of succinate controls reperfusion injury through mitochondrial ROS, Nature 515(7527) (2014) 431–435.

[16] E.T. Chouchani, V.R. Pell, A.M. James, L.M. Work, K. Saeb-Parsy, C. Frezza, T. Krieg, M.P. Murphy, A Unifying Mechanism for Mitochondrial Superoxide Production during Ischemia-Reperfusion Injury, Cell Metab 23(2) (2016) 254–63.

[17] D.J. Hausenloy, D.M. Yellon, Myocardial ischemia-reperfusion injury: a neglected therapeutic target, J Clin Invest 123(1) (2013) 92–100.

[18] Reha S. R, Robinson B. H, Mitochondria, Oxygen Free Radicals, and Apoptosis., Am J Med Genet (106) (2001) 62–70.

[19] M.D. Brand, The sites and topology of mitochondrial superoxide production, Exp Gerontol 45(7-8) (2010) 466–72.

[20] Reha S. R, Robinson B. H, Mitochondria, oxygen free radicals, disease and ageing, Treands Biochem Sci 25 (2000) 502–505.

[21] C.L. Quinlan, A.L. Orr, I.V. Perevoshchikova, J.R. Treberg, B.A. Ackrell, M.D. Brand, Mitochondrial complex II can generate reactive oxygen species at high rates in both the forward and reverse reactions, J Biol Chem 287(32) (2012) 27255–64.

[22] M.A. Konstam, M.S. Kiernan, D. Bernstein, B. Bozkurt, M. Jacob, N.K. Kapur, R.D. Kociol, E.F. Lewis, M.R. Mehra, F.D. Pagani, A.N. Raval, C. Ward, C. American Heart Association Council on Clinical, Y. Council on Cardiovascular Disease in the, S. Council on Cardiovascular, Anesthesia, Evaluation and Management of Right-Sided Heart Failure: A Scientific Statement From the American Heart Association, Circulation 137(20) (2018) e578–e622.

[23] V.P. Harjola, A. Mebazaa, J. Celutkiene, D. Bettex, H. Bueno, O. Chioncel, M.G. Crespo-Leiro, V. Falk, G. Filippatos, S. Gibbs, A. Leite-Moreira, J. Lassus, J. Masip, C. Mueller, W. Mullens, R. Naeije, A.V. Nordegraaf, J. Parissis, J.P. Riley, A. Ristic, G. Rosano, A. Rudiger, F. Ruschitzka, P. Seferovic, B. Sztrymf, A. Vieillard-Baron, M.B. Yilmaz, S. Konstantinides, Contemporary management of acute right ventricular failure: a statement from the Heart Failure Association and the Working Group on Pulmonary Circulation and Right Ventricular Function of the European Society of Cardiology, Eur J Heart Fail 18(3) (2016) 226–41.

[24] A.L. Orr, C.L. Quinlan, I.V. Perevoshchikova, M.D. Brand, A refined analysis of superoxide production by mitochondrial sn-glycerol 3-phosphate dehydrogenase, J Biol Chem 287(51) (2012) 42921–35.

[25] F.L. Muller, Y. Liu, M.A. Abdul-Ghani, M.S. Lustgarten, A. Bhattacharya, Y.C. Jang, H. Van Remmen, High rates of superoxide production in skeletal-muscle mitochondria respiring on both complex I- and complex II-linked substrates, Biochem J 409(2) (2008) 491–9.

[26] C.L. Quinlan, I.V. Perevoshchikova, M. Hey-Mogensen, A.L. Orr, M.D. Brand, Sites of reactive oxygen species generation by mitochondria oxidizing different substrates, Redox Biol 1 (2013) 304–12.

[27] I. Siebels, S. Drose, Q-site inhibitor induced ROS production of mitochondrial complex II is attenuated by TCA cycle dicarboxylates, Biochim Biophys Acta 1827(10) (2013) 1156–64.

[28] D.B. Zorov, M. Juhaszova, S.J. Sollott, Mitochondrial reactive oxygen species (ROS) and ROS-induced ROS release, Physiol Rev 94(3) (2014) 909–50.

[29] J.N. Bazil, D.A. Beard, K.C. Vinnakota, Catalytic Coupling of Oxidative Phosphorylation, ATP Demand, and Reactive Oxygen Species Generation, Biophys J 110(4) (2016) 962–71.

[30] J.N. Bazil, V.R. Pannala, R.K. Dash, D.A. Beard, Determining the origins of superoxide and hydrogen peroxide in the mammalian NADH:ubiquinone oxidoreductase, Free Radic Biol Med 77 (2014) 121–9.

[31] Schrödinger, The PyMOL molecular graphics system, Version 1.8., LLC, NewYark, NY. (2017).

[32] G. Cecchini, Function and structure of complex II of the respiratory chain, Annu Rev Biochem 72 (2003) 77–109.

[33] Todd. A.J, The generalized Inverse for Matrix., Mathematical Proc of the Cambridge Philosophical Society 51(3) (1954) 406–413.

[34] A.D. Vinogradov, V.G. Grivennikova, E. Gavrikova, V., Studies on the succinate dehydrogenating system. I. Kinetics of the succinate dehydrogenase interaction with a semiquindiimine radical of N,N,N,N tetramethyle p Phenylenediamine, BBA 545 (1979) 141–154.

[35] A.J. Jones, J. Hirst, A spectrophotometric coupled enzyme assay to measure the activity of succinate dehydrogenase, Anal Biochem 442(1) (2013) 19–23.

[36] W.P. Zeijlemaker, D.V. Derartanian, C. Veeger, E.C. Slater, Studies on succinate dehydrogenase IV: Kinetics of the overall reaction catalysed by preparations of the purified enzyme, BBA 178 (1969) 213–224.

[37] Tomoko Ohnishi, Tsoo E. King, John C. Salernog, Haywood Blum, JohnR. Bowyer, Takamitsu Maida, Thermodynamic and Electron Paramagnetic Resonance Characterization of Flavin in Succinate Dehydrogena, J Biol Chem 256 (1981) 5577–5582.

[38] R.J. Mailloux, Teaching the fundamentals of electron transfer reactions in mitochondria and the production and detection of reactive oxygen species, Redox Biol 4 (2015) 381–98.

[39] A.D. Vinogradov, E.V. Gavrikova, V.V. Zuevsky, Reactivity of the sulfhydryl groups of soluble succinate dehydrogenase, Eur J Biochem 63(2) (1976) 365–71.

[40] W.C. Kenney, The reaction of N-ethylmaleimide at the active site of succinate dehydrogenase, J Biol Chem 250(8) (1975) 3089–94.

[41] R. Horsefield, V. Yankovskaya, G. Sexton, W. Whittingham, K. Shiomi, S. Omura, B. Byrne, G. Cecchini, S. Iwata, Structural and computational analysis of the quinone-binding site of complex II (succinate-ubiquinone oxidoreductase): a mechanism of electron transfer and proton conduction during ubiquinone reduction, J Biol Chem 281(11) (2006) 7309–16.

[42] H. Miyadera, K. Shiomi, H. Ui, Y. Yamaguchi, R. Masuma, H. Tomoda, H. Miyoshi, A. Osanai, K. Kita, S. Omura, Atpenins, potent and specific inhibitors of mitochondrial complex II (succinate-ubiquinone oxidoreductase), Proc Natl Acad Sci U S A 100(2) (2003) 473–7.

[43] K.R. Messner, J.A. Imlay, Mechanism of superoxide and hydrogen peroxide formation by fumarate reductase, succinate dehydrogenase, and aspartate oxidase, J Biol Chem 277(45) (2002) 42563–71.

[44] V.G. Grivennikova, A.V. Kareyeva, A.D. Vinogradov, Oxygen-dependence of mitochondrial ROS production as detected by Amplex Red assay, Redox Biol 17 (2018) 192–199.

[45] A.D. Vinogradov, A.B. Kotlyar, V.I. Burov, Y.O. Belikova, Regulation of succinate dehydrogenase and tautomerization of oxaloacetate, Adv Enzyme Regul 28 (1989) 271–80.

[46] E. Maklashina, P. Hellwig, R.A. Rothery, V. Kotlyar, Y. Sher, J.H. Weiner, G. Cecchini, Differences in protonation of ubiquinone and menaquinone in fumarate reductase from Escherichia coli, J Biol Chem 281(36) (2006) 26655–64.

[47] W.G. Hanstein, K.A. Davis, M.A. Ghalambor, Y. Hatefi, Succinate dehydrogenase. II. Enzymatic properties, Biochemistry 10(13) (1971) 2517–24.

[48] J. Guo, B.D. Lemire, The ubiquinone-binding site of the Saccharomyces cerevisiae succinate-ubiquinone oxidoreductase is a source of superoxide, J Biol Chem 278(48) (2003) 47629–35.

[49] J.A. Imlay, I. Fridovich, Assay of metabolic superoxide production in Escherichia coli, J Biol Chem 266(11) (1991) 6957–65.

[50] M.P. Paranagama, K. Sakamoto, H. Amino, M. Awano, H. Miyoshi, K. Kita, Contribution of the FAD and quinone binding sites to the production of reactive oxygen species from Ascaris suum mitochondrial complex II, Mitochondrion 10(2) (2010) 158–65.

