## Supplementary Information for "Analysis of Mammalian Succinate Dehydrogenase Kinetics and Reactive Oxygen Species Production"

**Summary:** The supplementary material has four sections. The first section contains the heat map showing correlation between the adjustable parameters given in Table 1 of the main manuscript, the parameter table for fixed parameters, Table S1, and the environmental parameters, Table S2. The second section contains the list of mathematical equations that govern the model behavior. The third section lists the equations for the ODE model used to simulate the results in Figure 9 of the main manuscript. The fourth section details the Matlab code used to simulate the model and generate the figures.

#### Section 1

**Substrates:** succinate (SUC), quinone (Q), oxygen ($O_2$)
**Products:** fumarate (FUM), quinol ($QH_2$), superoxide, ($O_2^{\bullet -}$), and hydrogen peroxide ($H_2O_2$)
**Inhibitors:** atpenin (A5), malonate (MALO), oxaloacetate (OAA), malate (MAL)
**Other notations:** $[2Fe-2S]=ISC_1$, $[4Fe-4S]=ISC_2$, $[4Fe-3S]=ISC_3$

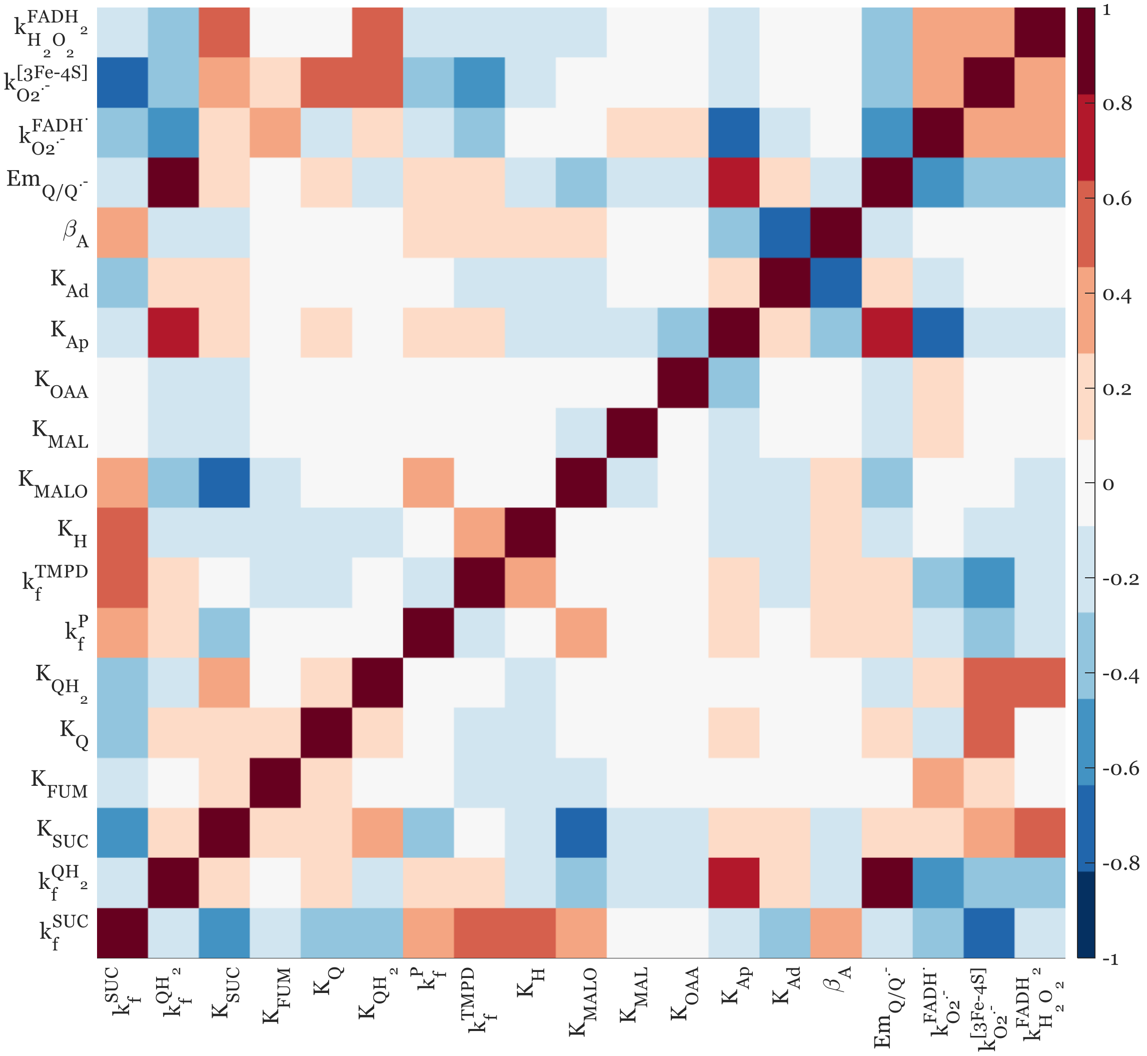


**Figure S1. Correlation heat map for adjustable parameters given in Table 2.** Matrix of correlation coefficient between the adjustable parameters. The normalized parameter sensitivity matrix is computed using Eq. S98. The sensitivity coefficients were computed from the parameter sensitivity matrix given in Eq. S199. Correlation coefficients range between −1 (negative correlation) to +1 (positive correlation). A coefficient value of 0 means the two parameters are uncorrelated.

**Table S1. Fixed Model Parameters**

| Parameters | Definition | Values | Units | References |
|---|---|---|---|---|
| $R$ | Ideal gas constant | 8.314 | J/mol/K | - |
| $F$ | Faraday's constant | 96.5 | J/mV/mol | - |
| $K_{FADH}$ | pKa for flavin free radical | 8 | - | (1) |

| | | | | |
|---|---|---|---|---|
| $K_{FADH_2}$ | pKa for fully reduce flavin | 7.7 | - | (1) |
| $E_m^{0\ FAD/FADH}$ | FAD/FADH midpoint potential | 385 | mV | (1) |
| $E_m^{0\ FADH/FADH_2}$ | FADH/$FADH_2$ midpoint potential | 284 | mV | (1) |
| $E_m^{0\ FAD/FADH_2}$ | FADH/$FADH_2$ midpoint potential | 333.8 | mV | (1) |
| $E_m^{ISC_1}$ | Midpoint potential of [*2Fe-2S*]$_{ox,,red}$ | 0 | mV | (2) |
| $E_m^{ISC_2}$ | Midpoint potential of [*4Fe-4S*]$_{ox,,red}$ | -260 | mV | (2) |
| $E_m^{ISC_3}$ | Midpoint potential of [*3Fe-4S*]$_{ox,,red}$ | 60 | mV | (2) |
| $E_m^{O_2/O_2^{\bullet -}}$ | $O_2/O_2^{\bullet -}$ midpoint potential | -160 | mV | (3) |
| $E_m^{O_2/H_2O_2}$ | $O_2/H_2O_2$ midpoint potential | 940 | mV | (3) |
| $E_m^{0\ FUM/SUC}$ | FUM/SUC midpoint potential | 445 | mV | (4) |
| $E_m^{0\ Q/QH_2}$ | Q/$QH_2$ midpoint potential | 464 | mV | (5) |
| $E_m^{0\ P}$ | Phenazine midpoint potential | 358 | mV | (6) |
| $E_m^{0\ TMPD}$ | TMPD midpoint potential | 270 | mV | (7) |
| $[Q]_{tot}$ | Total mitochondrial quinone concentration | 20 | mM | (8) |

Midpoint potentials are given at 273 K and pH 0 except for the ISCs, oxygen/superoxide, and oxygen/hydrogen peroxide couples. Those are given with respect to 273 K and pH 7. All potentials are given as reduction potentials.

**Table S2. Environmental Parameters**

| Parameters | Description | Value | Sensitivity | Rank |
|---|---|---|---|---|
| Q | succinate/$QH_2$ constant | 1 M | 0.102 | 3 |
| $Q_{bc1}$ | succinate/$QH_2$ constant in the presence of complex III inhibitors | 16.6 μM | 0.046 | 5 |
| $Q_{A5}$ | atpenin inhibitory constant | 1 M | $4.39x10^{-8}$ | 7 |
| $Q_{MAL}$ | malate inhibitory constant | 60.8 mM | $2.31x10^{-6}$ | 6 |
| $Q_{OAA}$ | oxaloacetate inhibitory constant | 0.43 nM | 0.345 | 1 |
| $Q_{FUM}$ | fumarate inhibitory constant | 8.31 μM | 0.050 | 4 |
| $Q_{MALO}$ | malonate inhibitory constant | 2.1 mM | 0.162 | 2 |

#### Section 2

**Quinol and quinone concentration equations**

$$[QH_2] = Q_{tot} / \left(1 + Q / [SUC]\right) \cdot \left(1 + [A5] / Q_{A5}\right) \cdot \left(1 + [MALO] / Q_{MALO}\right)$$
$$\cdot \left(1 + [MAL] / Q_{MAL}\right) \cdot \left(1 + [FUM] / Q_{FUM}\right) \cdot \left(1 + [OAA] / Q_{OAA}\right) \qquad \text{Eq. S1}$$

If complex III inhibitors are present, the following equation is used:

$$[QH_2] = Q_{tot} / \left(1 + Q_{bc_1} / [SUC]\right) \cdot \left(1 + [A5] / Q_{A5}\right) \cdot \left(1 + [MALO] / Q_{MALO}\right) \qquad \text{Eq. S2}$$
$$\cdot \left(1 + [MAL] / Q_{MAL}\right) \cdot \left(1 + [FUM] / Q_{FUM}\right) \cdot \left(1 + [OAA] / Q_{OAA}\right)$$

$$[Q] = [Q]_{tot} - [QH_2] \qquad \text{Eq. S3}$$

**Oxidation-reduction reactions**

One-electron reactions

$$FAD + H^+ + e^- \rightarrow FADH^\bullet \qquad \text{Eq. S4}$$

$$FADH^\bullet + H^+ + e^- \rightarrow FADH_2 \qquad \text{Eq. S5}$$

$$O_2 + e^- \rightarrow O_2^{\bullet -} \qquad \text{Eq. S6}$$

$$Q + e^- \rightarrow Q^{\bullet -} \qquad \text{Eq. S7}$$

$$Q^{\bullet -} + 2H^+ + e^- \rightarrow QH_2 \qquad \text{Eq. S8}$$

Two-electron reactions

$$FAD + 2H^+ + 2e^- \rightarrow FADH_2 \qquad \text{Eq. S9}$$

$$FUM + 2H^+ + 2e^- \rightarrow SUC \qquad \text{Eq. S10}$$

$$O_2 + 2H^+ + 2e^- \rightarrow H_2O_2 \qquad \text{Eq. S11}$$

$$Q + 2H^+ + 2e^- \rightarrow QH_2 \qquad \text{Eq. S12}$$

**Binding polynomials for enzyme, substrates, products and regulators**

$$P_{Qp} = 1 + \frac{[QH_2]}{K_{QH_2}} + \frac{[Q]}{K_Q} + \frac{[A5]}{K_{Ap}} \qquad \text{Eq. S13}$$

$$P_{Qd} = 1 + \frac{[A5]}{K_{Ad}} \qquad \text{Eq. S14}$$

$$P_{FAD} = 1 + \frac{[SUC]}{K_{SUC}} + \frac{[FUM]}{K_{FUM}} + \frac{[MAL]}{K_{MAL}} + \frac{[MALO]}{K_{MALO}} + \frac{[OAA]}{K_{OAA}} \qquad \text{Eq. S15}$$

**Forward rate constants for succinate oxidation and quinol reduction**

$$k_f SUC = k_{f_0} SUC \cdot (1 + [A5] / K_{A_2} \big/ (1 + \beta_A [A5] / K_{A_2}) / (1 + [H^+] \big/ K_H) \qquad \text{Eq. S16}$$

$$k_f QH_2 = k_{f_0} QH_2 \cdot (1 + [A5] / K_{A_2} \big/ (1 + \beta_A [A5] / K_{A_2}) \qquad \text{Eq. S17}$$

**Flavin and [*3Fe-4S*] free radical production rates**

$$k_f^{FADH^{g}} = k_{f_0}^{FADH^{g}} \big/ P_{FAD} \big/ (1 + [H^+] \big/ K_{FADH}) \qquad \text{Eq. S18}$$

$$k_f^{3Fe-4S} = k_{f_0}^{3Fe-4S} \big/ P_Q \qquad \text{Eq. S19}$$

$$k_f^{FADH_2} = k_{f_0}^{FADH_2} \Big/ P_{FAD} \Big/ (1+[H^+] \big/ K_{FADH_2}) \qquad \text{Eq. S20}$$

#### Midpoint potential pH corrections

$$E_m^{FAD/FADH} = E_m^{0\,FAD/FADH} + \left(RT/F\right).\log([H^+]\cdot(1+K_{FADH}/[H^+])) \qquad \text{Eq. S21}$$

$$E_m^{FADH/FADH_2} = E_m^{0\,FADH/FADH_2} + \left(RT/F\right)\log\left([H^+]\cdot(1+[H^+]/K_{FADH_2})/(1+[H^+]/K_{FADH})\right) \qquad \text{Eq. S22}$$

$$E_m^{FAD/FADH_2} = E_m^{0\,FAD/FADH_2} + \left(RT/2F\right)\log\begin{pmatrix}[H^+]^2\cdot(1+[H^+]/K_{FADH_2}) \\ \cdot(1+K_{FADH}/[H^+])/(1+[H^+]/K_{FADH})\end{pmatrix} \qquad \text{Eq. S23}$$

$$E_m^{SQ/QH_2} = 2E_m^{0\,SQ/QH_2} + 2\left(RT/F\right)\log([H^+]) - E_m^{Q/SQ} \qquad \text{Eq. S24}$$

$$E_m^{Q/QH_2} = E_m^{0\,Q/QH_2} + \left(RT/F\right)\log([H^+]) \qquad \text{Eq. S25}$$

$$E_m^{FUM/SUC} = E_m^{0\,FUM/SUC} + \left(RT/F\right)\log([H^+]) \qquad \text{Eq. S26}$$

$$E_m^{P} = E_m^{0\,P} + \left(RT/2F\right)\log([H^+]) \qquad \text{Eq. S27}$$

$$E_m^{TMPD} = E_m^{0\,TMPD} + \left(RT/2F\right)\log([H^+]) \qquad \text{Eq. S28}$$

#### Midpoint potential bound state corrections

$$E_{mb}^{Q/SQ} = E_m^{Q/SQ} + \left(RT/F\right)\log(K_Q) \qquad \text{Eq. S29}$$

$$E_{mb}^{SQ/QH_2} = E_m^{SQ/QH_2} - \left(RT/F\right)\log(K_{QH_2}) \qquad \text{Eq. S30}$$

$$E_{mb}^{FUM/SUC} = E_m^{FUM/SUCC} - \left(RT/2F\right)\log(K_{SUC}) + \left(RT/2F\right)\log(K_{FUM}) \qquad \text{Eq. S31}$$

#### Equilibrium constants

$$K_{eq}^{FUM/FADH_2} = e^{2F/RT\left(E_{mb}^{FUM/SUC} - E_m^{FAD/FADH_2}\right)} \qquad \text{Eq. S32}$$

$$K_{eq}^{QH_2/ISC_3} = e^{F/RT\left(E_m^{SQ/QH_2} - E_m^{ISC_3}\right)} \qquad \text{Eq. S33}$$

$$K_{eq}^{GEA/ISC_1ISC_3} = e^{2F/RT\left(E_m^{GEA} - E_m^{ISC3} - E_m^{ISC1}\right)} \qquad \text{Eq. S34}$$

$$K_{eq}^{O_2^{\bullet -}/ISC_3} = e^{F/RT\left(E_m^{O_2/O_2^{\bullet -}} - E_m^{ISC_3}\right)} \qquad \text{Eq. S35}$$

$$K_{eq}^{O_2^{\bullet -}/FADH} = e^{F/RT\left(E_m^{O_2/O_2^{\bullet -}} - E_m^{FAD/FADH}\right)} \qquad \text{Eq. S36}$$

$$K_{eq}^{H_2O_2/FADH_2} = e^{2F/RT\left(E_m^{O_2/H_2O_2} - E_m^{FAD/FADH_2}\right)} \qquad \text{Eq. S37}$$

Here, GEA stands for "general electron acceptor" and is used when either phenazine or TMPD is the acceptor. The reaction is considered a concerted two-electron reduction of the acceptor. We use $ISC_3$ and $ISC_1$ as the electron source since these are the redox centers most highly reduced. Using $ISC_3$ and $ISC_2$ only causes the forward rate constant for phenazine and TMPD reduction to increase which doesn't change the simulation results.

#### Boltzmann redox poise potentials

$$E_h^{FAD/FADH} = E_m^{FAD/FADH} \qquad \text{Eq. S38}$$

$$E_h^{FADH/FADH_2} = E_m^{FADH/FADH_2} \qquad \text{Eq. S39}$$

$$E_h^{ISC_1} = E_m^{ISC_1} \qquad \text{Eq. S40}$$

$$E_h^{ISC_2} = E_m^{ISC_2} \qquad \text{Eq. S41}$$

$$E_h^{ISC_3} = E_m^{ISC_3} \qquad \text{Eq. S42}$$

$$E_h^{Q/SQ} = E_{mb}^{Q/SQ} + \left(RT/F\right)\log(Q/K_Q/P_Q) \qquad \text{Eq. S43}$$

**Formation energies for each redox center**

Before calculating the substate fractions for each electron state, the midpoint potentials are converted into the free energies using the following equation.

$$\Delta G = -nF\Delta E \qquad \text{Eq. S44}$$

Here, n is the number of electrons transferred in the reaction.

**Substate fraction calculations.** To determining the transition rates, each combination of the redox centers (substates) reduced in each electronic state are calculated by Boltzmann distribution. Here, $S_r^k$ is the fraction of redox centers *r* existing in the electronic state *k* that is reduced, and $\Delta G_r^k$ are the free energy change for each redox center *r* calculated from the linear superposition of the midpoint potentials.

$$S_r^k = \frac{e^{-\Delta G_r^k/RT}}{\sum_r e^{-\Delta G_r^k/RT}} \qquad \text{Eq. S45}$$

**Denominators for each electronic state**

$$D_1 = e^{-\Delta G_{FADH}/RT} + e^{-\Delta G_{ISC_1}/RT} + e^{-\Delta G_{ISC_2}/RT} + e^{-\Delta G_{ISC_3}/RT} + e^{-\Delta G_{SQ}/RT} \qquad \text{Eq. S46}$$

$$\begin{aligned} D_2 &= \left(e^{-\Delta G_{FADH}/RT} e^{-\Delta G_{FADH_2}/RT}\right) + \left(e^{-\Delta G_{FADH}/RT} e^{-\Delta G_{ISC_1}/RT}\right) + \left(e^{-\Delta G_{FADH}/RT} e^{-\Delta G_{ISC_2}/RT}\right) \\ &+ \left(e^{-\Delta G_{FADH}/RT} e^{-\Delta G_{ISC_3}/RT}\right) + \left(e^{-\Delta G_{FADH}/RT} e^{-\Delta G_{SQ}/RT}\right) + \left(e^{-\Delta G_{ISC_1}/RT} e^{-\Delta G_{ISC_2}/RT}\right) + \left(e^{-\Delta G_{ISC_1}/RT} e^{-\Delta G_{ISC_3}/RT}\right) \\ &+ \left(e^{-\Delta G_{ISC_1}/RT} e^{-\Delta G_{SQ}/RT}\right) + \left(e^{-\Delta G_{ISC_2}/RT} e^{-\Delta G_{ISC_3}/RT}\right) + \left(e^{-\Delta G_{ISC_2}/RT} e^{-\Delta G_{SQ}/RT}\right) + \left(e^{-\Delta G_{ISC_3}/RT} e^{-\Delta G_{SQ}/RT}\right) \end{aligned} \qquad \text{Eq. S47}$$

$$\begin{aligned} D_3 &= \left(e^{-\Delta G_{FADH}/RT} e^{-\Delta G_{FADH_2}/RT} e^{-\Delta G_{ISC_1}/RT}\right) + \left(e^{-\Delta G_{FADH}/RT} e^{-\Delta G_{FADH_2}/RT} e^{-\Delta G_{ISC_2}/RT}\right) \\ &+ \left(e^{-\Delta G_{FADH}/RT} e^{-\Delta G_{FADH_2}/RT} e^{-\Delta G_{ISC_3}/RT}\right) + \left(e^{-\Delta G_{FADH}/RT} e^{-\Delta G_{FADH_2}/RT} e^{-\Delta G_{SQ}/RT}\right) \\ &+ \left(e^{-\Delta G_{FADH}/RT} e^{-\Delta G_{ISC_1}/RT} e^{-\Delta G_{ISC_2}/RT}\right) + \left(e^{-\Delta G_{FADH}/RT} e^{-\Delta G_{ISC_1}/RT} e^{-\Delta G_{ISC_3}/RT}\right) \\ &+ \left(e^{-\Delta G_{FADH}/RT} e^{-\Delta G_{ISC_1}/RT} e^{-\Delta G_{SQ}/RT}\right) + \left(e^{-\Delta G_{FADH}/RT} e^{-\Delta G_{ISC_2}/RT} e^{-\Delta G_{ISC_3}/RT}\right) \\ &+ \left(e^{-\Delta G_{FADH}/RT} e^{-\Delta G_{ISC_2}/RT} e^{-\Delta G_{SQ}/RT}\right) + \left(e^{-\Delta G_{FADH}/RT} e^{-\Delta G_{ISC_3}/RT} e^{-\Delta G_{SQ}/RT}\right) \\ &+ \left(e^{-\Delta G_{ISC_1}/RT} e^{-\Delta G_{ISC_2}/RT} e^{-\Delta G_{ISC_3}/RT}\right) + \left(e^{-\Delta G_{ISC_1}/RT} e^{-\Delta G_{ISC_2}/RT} e^{-\Delta G_{SQ}/RT}\right) \\ &+ \left(e^{-\Delta G_{ISC_1}/RT} e^{-\Delta G_{ISC_3}/RT} e^{-\Delta G_{SQ}/RT}\right) + \left(e^{-\Delta G_{ISC_2}/RT} e^{-\Delta G_{ISC_3}/RT} e^{-\Delta G_{SQ}/RT}\right) \end{aligned} \qquad \text{Eq. S48}$$

$$\begin{aligned} D_4 &= \left(e^{-\Delta G_{FADH}/RT} e^{-\Delta G_{FADH_2}/RT} e^{-\Delta G_{ISC_1}/RT} e^{-\Delta G_{ISC_2}/RT}\right) + \left(e^{-\Delta G_{FADH}/RT} e^{-\Delta G_{FADH_2}/RT} e^{-\Delta G_{ISC_1}/RT} e^{-\Delta G_{ISC_3}/RT}\right) \\ &+ \left(e^{-\Delta G_{FADH}/RT} e^{-\Delta G_{FADH_2}/RT} e^{-\Delta G_{ISC_1}/RT} e^{-\Delta G_{SQ}/RT}\right) + \left(e^{-\Delta G_{FADH}/RT} e^{-\Delta G_{FADH_2}/RT} e^{-\Delta G_{ISC_2}/RT} e^{-\Delta G_{ISC_3}/RT}\right) + \\ &+ \left(e^{-\Delta G_{FADH}/RT} e^{-\Delta G_{FADH_2}/RT} e^{-\Delta G_{ISC_2}/RT} e^{-\Delta G_{SQ}/RT}\right) + \left(e^{-\Delta G_{FADH}/RT} e^{-\Delta G_{FADH_2}/RT} e^{-\Delta G_{ISC_3}/RT} e^{-\Delta G_{SQ}/RT}\right) + \\ &+ \left(e^{-\Delta G_{FADH}/RT} e^{-\Delta G_{ISC_1}/RT} e^{-\Delta G_{ISC_2}/RT} e^{-\Delta G_{ISC_3}/RT}\right) + \left(e^{-\Delta G_{FADH}/RT} e^{-\Delta G_{ISC_1}/RT} e^{-\Delta G_{ISC_2}/RT} e^{-\Delta G_{SQ}/RT}\right) + \\ &+ \left(e^{-\Delta G_{FADH}/RT} e^{-\Delta G_{ISC_1}/RT} e^{-\Delta G_{ISC_3}/RT} e^{-\Delta G_{SQ}/RT}\right) + \left(e^{-\Delta G_{FADH}/RT} e^{-\Delta G_{ISC_2}/RT} e^{-\Delta G_{ISC_3}/RT} e^{-\Delta G_{SQ}/RT}\right) + \\ &+ \left(e^{-\Delta G_{ISC_1}/RT} e^{-\Delta G_{ISC_2}/RT} e^{-\Delta G_{ISC_3}/RT} e^{-\Delta G_{SQ}/RT}\right) \end{aligned} \qquad \text{Eq. S49}$$

***E₀* substates used in state transitions**

$$S_X^0 = 1 \qquad \text{Eq. S50}$$

"$X$" represents any redox center as all are oxidized in the $E_0$ electronic state.

***E₁* substates used in state transitions**

$$S^1_{FADH} = \frac{\left(e^{-\Delta G_{FADH}/RT}\right)}{D_1} \qquad \text{Eq. S51}$$

$$S^1_{FAD} = 1 - S^1_{FADH} \qquad \text{Eq. S52}$$

$$S^1_{ISC_{3,red}} = \frac{\left(e^{-\Delta G_{ISC3}/RT}\right)}{D_1} \qquad \text{Eq. S53}$$

$$S^1_{ISC_{3,ox}} = 1 - S^1_{ISC_3} \qquad \text{Eq. S54}$$

$$S^1_{\notin SQ, ISC_{3,ox}} = 1 - S^1_{ISC_3} - S^1_{SQ} \qquad \text{Eq. S55}$$

$$S^1_{ISC_{1,ox}, ISC_{3,ox}} = \frac{\left(e^{-\Delta G_{FADH}/RT} + e^{-\Delta G_{ISC_2}/RT} + e^{-\Delta G_{SQ}/RT}\right)}{D_1} \qquad \text{Eq. S56}$$

***E₂* substates used in state transitions**

$$S^2_{FADH} = \frac{\left(e^{-\Delta G_{FADH}/RT}\right)\left(e^{-\Delta G_{ISC_1}/RT} + e^{-\Delta G_{ISC_2}/RT} + e^{-\Delta G_{ISC_3}/RT} + e^{-\Delta G_{SQ}/RT}\right)}{D_2} \qquad \text{Eq. S57}$$

$$S^2_{FADH_2} = \frac{\left(e^{-\Delta G_{FADH}/RT} e^{-\Delta G_{FADH_2}/RT}\right)}{D_2} \qquad \text{Eq. S58}$$

$$S^2_{FAD} = 1 - S^2_{FADH} - S^2_{FADH_2} \qquad \text{Eq. S59}$$

$$S^2_{ISC_{3,red}} = \frac{\left(e^{-\Delta G_{ISC_3}/RT}\right)\left(e^{-\Delta G_{FADH}/RT} + e^{-\Delta G_{ISC_1}/RT} + e^{-\Delta G_{ISC_2}/RT} + e^{-\Delta G_{SQ}/RT}\right)}{D_2} \qquad \text{Eq. S60}$$

$$S^2_{ISC_{3,ox}} = 1 - S^2_{ISC_{3,red}} \qquad \text{Eq. S61}$$

$$S^2_{ISC_{1,red}, ISC_{3,red}} = \frac{\left(e^{-\Delta G_{ISC_3}/RT} e^{-\Delta G_{ISC_1}/RT}\right)}{D_2} \qquad \text{Eq. S62}$$

$$S^2_{ISC_{1,ox}, ISC_{3,ox}} = 1 - S^2_{ISC_{1,red}, ISC_{3,red}} \qquad \text{Eq. S63}$$

$$S^2_{SQ\sim ISC_{3,red}} = \frac{\left(e^{-\Delta G_{ISC_3}/RT} e^{-\Delta G_{SQ}/RT}\right)}{D_2} \qquad \text{Eq. S64}$$

$$S^2_{\notin SQ, ISC_{3,ox}} = 1 - S^2_{SQ\sim ISC_{3,red}} \qquad \text{Eq. S65}$$

***E₃* substates used in state transitions**

$$S^3_{FADH} = \frac{\left(e^{-\Delta G_{FADH}/RT}\right)\begin{pmatrix} e^{-\Delta G_{ISC_1}/RT} e^{-\Delta G_{ISC_2}/RT} + e^{-\Delta G_{ISC_1}/RT} e^{-\Delta G_{ISC_3}/RT} + e^{-\Delta G_{ISC_1}/RT} e^{-\Delta G_{SQ}/RT} \\ + e^{-\Delta G_{ISC_2}/RT} e^{-\Delta G_{ISC_3}/RT} + e^{-\Delta G_{ISC_2}/RT} e^{-\Delta G_{SQ}/RT} + e^{-\Delta G_{ISC_3}/RT} e^{-\Delta G_{SQ}/RT} \end{pmatrix}}{D_3} \qquad \text{Eq. S66}$$

$$S^3_{FADH_2} = \frac{\left(e^{-\Delta G_{FADH}/RT} e^{-\Delta G_{FADH_2}/RT}\right)\left(e^{-\Delta G_{ISC_1}/RT} + e^{-\Delta G_{ISC_2}/RT} + e^{-\Delta G_{ISC_3}/RT} + e^{-\Delta G_{SQ}/RT}\right)}{D_3} \qquad \text{Eq. S67}$$

$$S^3_{FAD} = 1 - S^3_{FADH} - S^3_{FADH_2} \qquad \text{Eq. S68}$$

$$S^3_{ISC_{3,red}} = \frac{\left(e^{-\Delta G_{ISC_3}/RT}\right)\begin{pmatrix} e^{-\Delta G_{FADH}/RT}e^{-\Delta G_{FADH_2}/RT} + e^{-\Delta G_{FADH}/RT}e^{-\Delta G_{ISC_1}/RT} + e^{-\Delta G_{FADH}/RT}e^{-\Delta G_{ISC_2}/RT} \\ + e^{-\Delta G_{FADH}/RT}e^{-\Delta G_{SQ}/RT} + e^{-\Delta G_{ISC_1}/RT}e^{-\Delta G_{ISC_2}/RT} + e^{-\Delta G_{ISC_1}/RT}e^{-\Delta G_{SQ}/RT} \\ + e^{-\Delta G_{ISC_2}/RT}e^{-\Delta G_{SQ}/RT} \end{pmatrix}}{D_3} \quad \text{Eq. S69}$$

$$S^3_{ISC_{3,ox}} = 1 - S^3_{ISC_{3,red}} \quad \text{Eq. S70}$$

$$S^3_{ISC_{1,red},ISC_{3,red}} = \frac{\left(e^{-\Delta G_{ISC_1}/RT}e^{-\Delta G_{ISC_3}/RT}\right)\left(e^{-\Delta G_{FADH}/RT} + e^{-\Delta G_{ISC_2}/RT} + e^{-\Delta G_{SQ}/RT}\right)}{D_3} \quad \text{Eq. S71}$$

$$S^3_{SQ\sim ISC_{3,red}} = \frac{\left(e^{-\Delta G_{ISC_3}/RT}e^{-\Delta G_{SQ}/RT}\right)\left(e^{-\Delta G_{FADH}/RT} + e^{-\Delta G_{ISC_1}/RT} + e^{-\Delta G_{ISC_2}/RT}\right)}{D_3} \quad \text{Eq. S72}$$

**$E_4$ substates used in state transitions**

$$S^4_{FADH} = \frac{\left(e^{-\Delta G_{FADH}/RT}\right)\begin{pmatrix} e^{-\Delta G_{ISC_1}/RT}e^{-\Delta G_{ISC_2}/RT}e^{-\Delta G_{ISC_3}/RT} + e^{-\Delta G_{ISC_1}/RT}e^{-\Delta G_{ISC_2}/RT}e^{-\Delta G_{SQ}/RT} \\ + e^{-\Delta G_{ISC_1}/RT}e^{-\Delta G_{ISC_3}/RT}e^{-\Delta G_{SQ}/RT} + e^{-\Delta G_{ISC_1}/RT}e^{-\Delta G_{ISC_3}/RT}e^{-\Delta G_{SQ}/RT} \end{pmatrix}}{D_4} \quad \text{Eq. S73}$$

$$S^4_{FADH_2} = \frac{\left(e^{-\Delta G_{FADH}/RT}e^{-\Delta G_{FADH_2}/RT}\right)\begin{pmatrix} e^{-\Delta G_{ISC_1}/RT}e^{-\Delta G_{ISC_2}/RT} + e^{-\Delta G_{ISC_1}/RT}e^{-\Delta G_{ISC_3}/RT} + e^{-\Delta G_{ISC_2}/RT}e^{-\Delta G_{ISC_3}/RT} \\ + e^{-\Delta G_{ISC_1}/RT}e^{-\Delta G_{SQ}/RT} + e^{-\Delta G_{ISC_2}/RT}e^{-\Delta G_{SQ}/RT} + e^{-\Delta G_{ISC_3}/RT}e^{-\Delta G_{SQ}/RT} \end{pmatrix}}{D_4} \quad \text{Eq. S74}$$

$$S^4_{ISC_{3,red}} = \frac{\left(e^{-\Delta G_{ISC_3}/RT}\right)\begin{pmatrix} e^{-\Delta G_{FADH}/RT}e^{-\Delta G_{FADH_2}/RT}e^{-\Delta G_{ISC_1}/RT} + e^{-\Delta G_{FADH}/RT}e^{-\Delta G_{FADH_2}/RT}e^{-\Delta G_{ISC_2}/RT} \\ + e^{-\Delta G_{FADH}/RT}e^{-\Delta G_{FADH_2}/RT}e^{-\Delta G_{SQ}/RT} + e^{-\Delta G_{FADH}/RT}e^{-\Delta G_{ISC_1}/RT}e^{-\Delta G_{ISC_2}/RT} \\ + e^{-\Delta G_{FADH}/RT}e^{-\Delta G_{ISC_1}/RT}e^{-\Delta G_{SQ}/RT} + e^{-\Delta G_{FADH}/RT}e^{-\Delta G_{ISC_2}/RT}e^{-\Delta G_{SQ}/RT} \\ + e^{-\Delta G_{ISC_1}/RT}e^{-\Delta G_{ISC_2}/RT}e^{-\Delta G_{SQ}/RT} \end{pmatrix}}{D_3} \quad \text{Eq. S75}$$

$$S^4_{ISC_{1,red},ISC_{3,red}} = \frac{\left(e^{-\Delta G_{ISC_1}/RT}e^{-\Delta G_{ISC_3}/RT}\right)\left(e^{-\Delta G_{FADH}/RT}e^{-\Delta G_{FADH_2}/RT} + e^{-\Delta G_{FADH}/RT}e^{-\Delta G_{ISC_2}/RT} + e^{-\Delta G_{FADH}/RT}e^{-\Delta G_{SQ}/RT}\right.}{D_3} \quad \text{Eq. S76}$$

$$S^4_{SQ\sim ISC_{3,red}} = \frac{\left(e^{-\Delta G_{SQ}/RT}e^{-\Delta G_{ISC_3}/RT}\right)\left(e^{-\Delta G_{FADH}/RT}e^{-\Delta G_{FADH_2}/RT} + e^{-\Delta G_{FADH}/RT}e^{-\Delta G_{ISC_1}/RT} + e^{-\Delta G_{FADH}/RT}e^{-\Delta G_{ISC_2}/RT}\right)}{D_4} \quad \text{Eq. S77}$$

**State transition details and rates**

$$E_0 \rightarrow E_1$$

$$O_2^{\bullet -} + FAD \rightarrow O_2 + FAD^{\bullet -}$$

$$O_2^{\bullet -} + ISC_{3,ox} \rightarrow O_2 + ISC_{3,red}$$

1) $$k_{01}^{O_2^{\bullet -}/FAD} = \frac{k_f^{FADH^{g}}}{K_{eq}^{SO/FADH}}[O_2^{\bullet -}]S^0$$ Eq. S78

$$k_{01}^{O_2^{\bullet -}/3Fe-4S} = \frac{k_f^{3Fe-4S}}{K_{eq}^{O_2^{\bullet -}/ISC_3}}[O_2^{\bullet -}]S^0$$

$$k_{01} = k_{01}^{O_2^{\bullet -}/FAD} + k_{01}^{O_2^{\bullet -}/3Fe-4S}$$

$$E_1 \rightarrow E_0$$

$$FAD^{\bullet -} + O_2 \rightarrow FAD + O_2^{\bullet -}$$

$$ISC_{3,red} + O_2 \rightarrow ISC_{3,ox} + O_2^{\bullet -}$$

2) $$k_{10}^{O_2/FADH^{g}} = k_f^{FADH^{g}}[O_2]S_{FADH}^1$$ Eq. S79

$$k_{10}^{O_2/ISC_3} = k_f^{3Fe-4S}[O_2]S_{ISC_{3,red}}^1$$

$$k_{10} = k_{10}^{O_2/FADH^{g}} + k_{10}^{O_2/ISC_3}$$

$$E_0 \rightarrow E_2$$

$$H_2O_2 + FAD \rightarrow O_2 + FADH_2$$

$$SUC + FAD \rightarrow FUM + FADH_2$$

$$QH_2 + [4Fe-3S]_{ox} \rightarrow Q^{\bullet -} : \; [4Fe-3S]_{red} + 2H^+$$

$$GEA_{red} + [4Fe-3S]_{ox}, [2Fe-2S]_{ox} \rightarrow GEA_{ox} + [4Fe-3S]_{red}, [2Fe-2S]_{red}$$

$$k_{02}^{H_2O_2/FAD} = \frac{k_f^{FADH_2}}{k_{eq}^{H_2O_2/FADH_2}}[H_2O_2]S^0$$

3) $$k_{02}^{SUC/FAD} = k_f^{SUC}\frac{[SUC]/K_{SUC}}{P_{FAD}}S^0$$ Eq. S80

$$k_{02}^{QH_2/\notin SQ, ISC_{3,ox}} = \frac{k_f^{QH_2}}{K_{eq}^{QH_2/ISC_3}}\frac{[QH_2]/K_{QH_2}}{P_Q}S^0$$

$$k_{02}^{GEA/ISC_{1,ox}, ISC_{3,ox}} = \frac{k_f^{GEA}}{K_{eq}^{GEA/ISC_1 ISC_3}}[GEA]S^0$$

$$k_{02} = k_{02}^{H_2O_2/FAD} + k_{02}^{SUC/FAD} + k_{02}^{QH_2/\notin SQ, ISC_{3,ox}} + k_{02}^{GEA/ISC_{1,ox}, ISC_{3,ox}}$$

$$E_2 \rightarrow E_0$$

$$O_2 + FADH_2 \rightarrow H_2O_2 + FAD$$

$$FADH_2 + FUM \rightarrow FAD + SUC$$

$$Q^{\bullet-} : \ [4Fe-3S]_{red} + 2H^+ \rightarrow QH_2 + [4Fe-3S]_{ox}$$

$$GEA_{ox} + [4Fe-3S]_{red}, [2Fe-2S]_{red} \rightarrow GEA_{red} + [4Fe-3S]_{ox}, [2Fe-2S]_{ox}$$

4) $$k_{20}^{O_2/FADH_2} = k_f^{H_2O_2} [O_2] S_{FADH_2}^2$$ Eq. S81

$$k_{20}^{FUM/FADH_2} = \frac{k_f^{SUC}}{k_{eq}^{FUM/FADH_2}} \frac{[FUM]/K_{FUM}}{P_{FAD}} S_{FADH_2}^2$$

$$k_{20}^{SQ\sim ISC_3/QH_2} = k_f^{QH_2} S_{SQ\sim ISC_{3,red}}^2$$

$$k_{20}^{ISC_1 ISC_3/GAE} = k_f^{GAE} S_{ISC_{1,red}, ISC_{3,red}}^2$$

$$k_{20} = k_{20}^{O_2/H_2O_2} + k_{20}^{FUM/FADH_2} + k_{20}^{SQ\sim ISC_3/QH_2} + k_{20}^{ISC_1 ISC_3/GAE}$$

$$E_1 \rightarrow E_2$$

$$O_2^{\bullet-} + FAD \rightarrow O_2 + FAD^{\bullet-}$$

$$O_2^{\bullet-} + \left[3Fe-4S\right]_{Ox} \rightarrow O_2 + \left[3Fe-4S\right]_{red}$$

5) $$k_{12}^{O_2^{\bullet-}/FAD} = \frac{k_f^{FADH}}{K_{eq}^{SO/FADH}} [O_2^{\bullet-}] S_{FAD}^1$$ Eq. S82

$$k_{12}^{O_2^{\bullet-}/3Fe-4S} = \frac{k_f^{3Fe-4S}}{K_{eq}^{O_2^{\bullet-}/ISC_3}} [O_2^{\bullet-}] S_{ISC_{3,ox}}^1$$

$$k_{12} = k_{12}^{O_2^{\bullet-}/FAD} + k_{12}^{O_2^{\bullet-}/3Fe-4S}$$

$$E_2 \rightarrow E_1$$

$$FAD^{\bullet-} + O_2 \rightarrow FAD + O_2^{\bullet-}$$

$$[3Fe-4S]_{red} + O_2 \rightarrow [3Fe-4S]_{ox} + O_2^{\bullet-}$$

6) $$k_{21}^{O_2/FADH^{g}} = k_f^{FADH^{g}} [O_2] S_{FADH}^2$$ Eq. S83

$$k_{21}^{O_2/3Fe-4S} = k_f^{3Fe-4S} [O_2] S_{ISC_{3,red}}^2$$

$$k_{21} = k_{21}^{O_2/FADH^{g}} + k_{21}^{O_2/3Fe-4S}$$

$$E_1 \rightarrow E_3$$
$$H_2O_2 + FAD \rightarrow O_2 + FADH_2$$
$$SUC + FAD \rightarrow FUM + FADH_2$$
$$QH_2 + [4Fe-3S]_{ox} \rightarrow Q^{\bullet-} : [4Fe-3S]_{red} + 2H^+$$
$$GEA_{red} + [4Fe-3S]_{ox}, [2Fe-2S]_{ox} \rightarrow GEA_{ox} + [4Fe-3S]_{red}, [2Fe-2S]_{red}$$

7)

$$k_{13}^{H_2O_2/FAD} = \frac{k_f^{FADH_2}}{k_{eq}^{H_2O_2/FADH_2}}[H_2O_2]S_{FAD}^1$$
$$k_{13}^{SUC/FAD} = k_f^{SUC}\frac{[SUC]/K_{SUC}}{P_{FAD}}S_{FAD}^1 \qquad \text{Eq. S84}$$
$$k_{13}^{QH_2/\notin SQ, ISC_{3,ox}} = \frac{k_f^{QH_2}}{K_{eq}^{QH_2/ISC_3}}\frac{[QH_2]/K_{QH_2}}{P_Q}S_{\notin SQ, ISC_{3,ox}}^1$$
$$k_{13}^{GEA/ISC_{1,ox}, ISC_{3,ox}} = \frac{k_f^{GEA}}{K_{eq}^{GEA/ISC_1ISC_3}}[GEA]S_{ISC_{1,ox}, ISC_{3,ox}}^1$$
$$k_{13} = k_{13}^{H_2O_2/FAD} + k_{13}^{SUC/FAD} + k_{13}^{QH_2/\notin SQ, ISC_{3,ox}} + k_{13}^{GEA/ISC_{1,ox}, ISC_{3,ox}}$$

$$E_3 \rightarrow E_1$$
$$O_2 + FADH_2 \rightarrow H_2O_2 + FAD$$
$$FADH_2 + FUM \rightarrow FAD + SUC$$
$$Q^{\bullet-} : [4Fe-3S]_{red} + 2H^+ \rightarrow QH_2 + [4Fe-3S]_{ox}$$
$$GEA_{ox} + [4Fe-3S]_{red}, [2Fe-2S]_{red} \rightarrow GEA_{red} + [4Fe-3S]_{ox}, [2Fe-2S]_{ox}$$

8)

$$k_{31}^{O_2/FADH_2} = k_f^{H_2O_2}[O_2]S_{FADH_2}^3 \qquad \text{Eq. S85}$$
$$k_{31}^{FUM/FADH_2} = \frac{k_f^{SUC}}{k_{eq}^{FUM/FADH_2}}\frac{[FUM]/K_{FUM}}{P_{FAD}}S_{FADH_2}^3$$
$$k_{31}^{SQ\sim ISC_3/QH_2} = k_f^{QH_2}S_{SQ\sim[4Fe-3S]_{red}}^3$$
$$k_{31}^{ISC_1ISC_3/GAE} = k_f^{GAE}S_{ISC_{1,red}, ISC_{3,red}}^3$$
$$k_{31} = k_{31}^{O_2/H_2O_2} + k_{31}^{FUM/FADH_2} + k_{31}^{SQ\sim ISC_3/QH_2} + k_{31}^{ISC_1ISC_3/GAE}$$

$$E_2 \rightarrow E_3$$
$$O_2^{\bullet-} + FAD \rightarrow O_2 + FAD^{\bullet-}$$
$$O_2^{\bullet-} + \left[3Fe-4S\right]_{Ox} \rightarrow O_2 + \left[3Fe-4S\right]_{red}$$

9)

$$k_{23}^{O_2^{\bullet-}/FAD} = \frac{k_f^{FADH}}{K_{eq}^{SO/FADH}}[O_2^{\bullet-}]S_{FAD}^2 \qquad \text{Eq. S86}$$
$$k_{23}^{O_2^{\bullet-}/3Fe-4S} = \frac{k_f^{3Fe-4S}}{K_{eq}^{O_2^{\bullet-}/ISC_3}}[O_2^{\bullet-}]S_{ISC_{3,ox}}^2$$
$$k_{23} = k_{23}^{O_2^{\bullet-}/FAD} + k_{23}^{O_2^{\bullet-}/3Fe-4S}$$

$$E_3 \rightarrow E_2$$

$$FAD^{\bullet -} + O_2 \rightarrow FAD + O_2^{\bullet -}$$

$$[3Fe-4S]_{red} + O_2 \rightarrow [3Fe-4S]_{ox} + O_2^{\bullet -}$$

10) $$k_{32}^{O_2/FADH^{g}} = k_f^{FADH^{g}} [O_2] S_{FADH}^3$$ Eq. S87

$$k_{32}^{O_2/3Fe-4S} = k_f^{3Fe-4S} [O_2] S_{ISC_{3,red}}^3$$

$$k_{32} = k_{32}^{O_2/FADH^{g}} + k_{32}^{O_2/3Fe-4S}$$

$$E_2 \rightarrow E_4$$

$$H_2O_2 + FAD \rightarrow O_2 + FADH_2$$

$$SUC + FAD \rightarrow FUM + FADH_2$$

$$QH_2 + [4Fe-3S]_{ox} \rightarrow Q^{\bullet -} : \ [4Fe-3S]_{red} + 2H^+$$

$$GEA_{red} + [4Fe-3S]_{ox}, [2Fe-2S]_{ox} \rightarrow GEA_{ox} + [4Fe-3S]_{red}, [2Fe-2S]_{red}$$

$$k_{24}^{H_2O_2/FAD} = \frac{k_f^{FADH_2}}{k_{eq}^{H_2O_2/FADH_2}} [H_2O_2] S_{FAD}^2$$

11) Eq. S88

$$k_{24}^{SUC/FAD} = k_f^{SUC} \frac{[SUC]/K_{SUC}}{P_{FAD}} S_{FAD}^2$$

$$k_{24}^{QH_2/\notin SQ, ISC_{3,ox}} = \frac{k_f^{QH_2}}{K_{eq}^{QH_2/ISC_3}} \frac{[QH_2]/K_{QH_2}}{P_Q} S_{\notin SQ, ISC_{3,ox}}^2$$

$$k_{24}^{GEA/ISC_{1,ox}, ISC_{3,ox}} = \frac{k_f^{GEA}}{K_{eq}^{GEA/ISC_1 ISC_3}} [GEA] S_{ISC_{1,ox}, ISC_{3,ox}}^2$$

$$k_{24} = k_{24}^{H_2O_2/FAD} + k_{24}^{SUC/FAD} + k_{24}^{QH_2/\notin SQ, ISC_{3,ox}} + k_{24}^{GEA/ISC_{1,ox}, ISC_{3,ox}}$$

$$E_4 \rightarrow E_2$$

$$O_2 + FADH_2 \rightarrow H_2O_2 + FAD$$

$$FADH_2 + FUM \rightarrow FAD + SUC$$

$$Q^{\bullet -} : \ [4Fe-3S]_{red} + 2H^+ \rightarrow QH_2 + [4Fe-3S]_{ox}$$

$$GEA_{ox} + [4Fe-3S]_{red}, [2Fe-2S]_{red} \rightarrow GEA_{red} + [4Fe-3S]_{ox}, [2Fe-2S]_{ox}$$

$$k_{42}^{O_2/FADH_2} = k_f^{H_2O_2} [O_2] S_{FADH_2}^4$$

12) Eq. S89

$$k_{42}^{FUM/FADH_2} = \frac{k_f^{SUC}}{k_{eq}^{FUM/FADH_2}} \frac{[FUM]/K_{FUM}}{P_{FAD}} S_{FADH_2}^4$$

$$k_{42}^{SQ\sim ISC_3/QH_2} = k_f^{QH_2} S_{SQ\sim[4Fe-3S]_{red}}^4$$

$$k_{42}^{ISC_1 ISC_3/GAE} = k_f^{GAE} S_{ISC_{1,red}, ISC_{3,red}}^4$$

$$k_{42} = k_{42}^{O_2/H_2O_2} + k_{42}^{FUM/FADH_2} + k_{42}^{SQ\sim ISC_3/QH_2} + k_{42}^{ISC_1 ISC_3/GAE}$$

$$E_3 \rightarrow E_4$$
$$O_2^{\bullet-} + FAD \rightarrow O_2 + FAD^{\bullet-}$$
$$O_2^{\bullet-} + [3Fe-4S]_{Ox} \rightarrow O_2 + [3Fe-4S]_{red}$$

13) $$k_{34}^{O_2^{\bullet-}/FAD} = \frac{k_f^{FADH}}{K_{eq}^{SO/FADH}}[O_2^{\bullet-}]S_{FAD}^3$$ Eq. S90

$$k_{34}^{O_2^{\bullet-}/3Fe-4S} = \frac{k_f^{3Fe-4S}}{K_{eq}^{O_2^{\bullet-}/ISC_3}}[O_2^{\bullet-}]S_{ISC_{3,ox}}^3$$

$$k_{34} = k_{34}^{O_2^{\bullet-}/FAD} + k_{34}^{O_2^{\bullet-}/3Fe-4S}$$

$$E_4 \rightarrow E_3$$
$$FAD^{\bullet-} + O_2 \rightarrow FAD + O_2^{\bullet-}$$
$$[3Fe-4S]_{red} + O_2 \rightarrow [3Fe-4S]_{ox} + O_2^{\bullet-}$$

14) $$k_{43}^{O_2/FADH^{g}} = k_f^{FADH^{g}}[O_2]S_{FADH}^4$$ Eq. S91

$$k_{43}^{O_2/3Fe-4S} = k_f^{3Fe-4S}[O_2]S_{ISC_{3,red}}^4$$

$$k_{43} = k_{43}^{O_2/FADH^{g}} + k_{43}^{O_2/3Fe-4S}$$

**Steady-state equations**

$$\begin{bmatrix} -(k_{01}+k_{02}) & k_{01} & k_{02} & 0 & 0 \\ k_{01} & -(k_{10}+k_{12}+k_{13}) & k_{21} & k_{31} & 0 \\ k_{02} & k_{12} & -(k_{20}+k_{21}+k_{23}+k_{24}) & k_{32} & k_{42} \\ 0 & k_{13} & k_{23} & -(k_{31}+k_{32}+k_{43}) & k_{34} \\ 0 & 0 & k_{24} & k_{34} & -(k_{42}+k_{43}) \\ 1 & 1 & 1 & 1 & 1 \end{bmatrix} \begin{bmatrix} E_0 \\ E_1 \\ E_2 \\ E_3 \\ E_4 \end{bmatrix} = \begin{bmatrix} 0 \\ 0 \\ 0 \\ 0 \\ 0 \\ 1 \end{bmatrix}$$ Eq. S92

Five electron states were the minimal number required to fit all the data. Increasing the number to the maximum allowable of seven states does not significantly improve the model fits to the data.

**Succinate oxidation steady-state rate**

$$J_{SUC} = E_{tot}\left(k_{02}^{SUC/FAD}E_0 + k_{13}^{SUC/FAD}E_1 + k_{24}^{SUC/FAD}E_2 - k_{20}^{FUM/FADH_2}E_2 - k_{31}^{FUM/FADH_2}E_3 - k_{42}^{FUM/FADH_2}E_4\right)$$ Eq. S93

**Superoxide formation steady-state rate**

$$J_{O_2^{\bullet-}} = E_{tot}\begin{pmatrix} k_{10}^{O_2/FADH}E_1 + k_{21}^{O_2/FADH}E_2 + k_{32}^{O_2/FADH}E_3 + k_{43}^{O_2/FADH}E_4 - k_{01}^{O_2^{\bullet-}/FAD}E_0 - k_{12}^{O_2^{\bullet-}/FAD}E_1 - k_{23}^{O_2^{\bullet-}/FAD}E_2 - k_{34}^{O_2^{\bullet-}/FAD}E_3 \\ +k_{10}^{O_2/3Fe-4S_{red}}E_1 + k_{21}^{O_2/3Fe-4S_{red}}E_2 + k_{32}^{O_2/3Fe-4S_{red}}E_3 + k_{43}^{O_2/3Fe-4S_{red}}E_4 - k_{01}^{O_2^{\bullet-}/3Fe-4S_{ox}}E_0 - k_{12}^{O_2^{\bullet-}/3Fe-4S_{ox}}E_1 \\ -k_{23}^{O_2^{\bullet-}/3Fe-4S_{ox}}E_2 - k_{34}^{O_2^{\bullet-}/3Fe-4S_{ox}}E_3 \end{pmatrix}$$ Eq. S94

**Hydrogen peroxide formation steady-state rate**

$$J_{H_2O_2} = E_{tot}\left(k_{20}^{O_2/FADH_2}E_2 + k_{31}^{O_2/FADH_2}E_3 + k_{42}^{O_2/FADH_2}E_4 - k_{02}^{FADH_2/FAD}E_0 - k_{13}^{FADH_2/FAD}E_1 - k_{24}^{FADH_2/FAD}E_2\right)$$ Eq. S95

**Quinol reduction steady-state rate**

$$J_{QH_2} = J_{SUC} - J_{O_2^{\bullet-}} / 2 - J_{H_2O_2} \qquad \text{Eq. S96}$$

**Phenazine and TMPD Reduction steady-state rate**

$$J_{GEA} = J_{SUC} - J_{O_2^{\bullet-}} / 2 - J_{H_2O_2} \qquad \text{Eq. S97}$$

**Parameter sensitivity matrix and correlation coefficients.** The normalized parameter sensitivity matrix is computed using Eq. S98. Each model output, $f_i$, is congruent with the experimental data. The parameter sensitivities were computed using the complex variable approach as described in Squire and Trap (9).

$$S_{i,j} = \frac{df_i}{dp_j} \frac{p_j}{f_i} \qquad \text{Eq. S98}$$

The sensitivity coefficients presented in Table 2 were computed by averaging all the non-zero sensitivity coefficient for a given parameter. This was done by using Eq. S99.

$$\bar{S}_j = \frac{1}{N_i} \sum_{\forall i : \left|\frac{df_i}{dp_j}\right| > 0} \left| S_{i,j} \right| \qquad \text{Eq. S99}$$

Where $N_i$ is the number of non-zero elements in $i^{th}$ row of $S$. The parameter correlation coefficients are pairwise linear correlation coefficients computed for each pair of columns in the normalized parameter sensitivity matrix.

#### Section 3

$$\frac{d[SUC]}{dt} = -J_{SUC}$$ Eq. S100

$$\frac{d[FUM]}{dt} = J_{SUC}$$ Eq. S101

$$J_{O_2} = 5\times10^{-7}\left[[QH_2]/\left([QH_2]+10^{-6}\right)\right]\cdot\left[[O_2]/\left([O_2]+5\times10^{-7}\right)\right]$$ Eq. S102

$$\frac{d[QH_2]}{dt} = \left(2\cdot J_{O_2} - J_{QH_2}\right)/V_{lp}$$ Eq. S103

$$\frac{d[QH_2]}{dt} = -\left(2\cdot J_{O_2} - J_{QH_2}\right)/V_{lp}$$ Eq. S104

$$\frac{d[O_2]}{dt} = -\left(J_{O_2^{\bullet -}} + J_{H_2O_2} + J_{O_2}\right)$$ Eq. S105

$$\frac{d[H_2O_2]}{dt} = J_{O_2^{\bullet -}}/2 + J_{H_2O_2}$$ Eq. S106

This system was integrated using ode15s with default tolerances and the backwards differentiation formulas. For Eq. S102, the oxygen consumption rate was taken from the data (10) (30 µM/sec) and converted to M/min. The enzyme concentration, $[E_{tot}]$ was 20 nM computed from 0.2 mg/ml SMP and 100 pmol SDH/mg taken from (11). The kinetic constants for $QH_2$ and $O_2$ were estimated from prior work (8,12). The quinone dynamics were computed with respect to a lipid volume fraction, $V_{lp}$, of approximately 50 µl lipid volume per liter assay buffer based on 250 nl/mg lipid volume (8) and 0.2 mg/ml SMP concentration.

#### Section 4

Model Code Description

SDH_model_code.zip - zip file containing the following files:
data.mat - mat file containing data structure
SDH_parameters.mat - mat file containing model parameters
main_complexII.m - function containing SDH model code
Generate_Figures.m - function containing code to simulate the model and plot the figures

### References


1. Tomoko Ohnishi, Tsoo E . King, John C . Salernog, Haywood Blum, JohnR . Bowyer, and Takamitsu Maida. (1981) Thermodynamic and Electron Paramagnetic Resonance Characterization of Flavin in Succinate Dehydrogena. *J Biol Chem* **256**, 5577-5582
2. Grivennikova, V. G., Kozlovsky, V. S., and Vinogradov, A. D. (2017) Respiratory complex II: ROS production and the kinetics of ubiquinone reduction. *Biochim Biophys Acta Bioenerg* **1858**, 109-117
3. Mailloux, R. J. (2015) Teaching the fundamentals of electron transfer reactions in mitochondria and the production and detection of reactive oxygen species. *Redox Biol* **4**, 381-398
4. Tinoco, I., Sauer, K., and Wang, J. C. (1995) *Physical chemistry : principles and applications in biological sciences*, 3rd ed., Prentice Hall, Englewood Cliffs, N.J.
5. Alberty, R. A. (2003) *Thermodynamics of biochemical reactions*, Wiley-Interscience, Hoboken, N.J.
6. Ksenzhek, O. S., Petrova, S. A., and Kolodyazhny, M. V. (1977) Electrochemical properties of some redox indicators. *Bioelectrochemistry and Bioenergetics* **4**, 346-357
7. Szentrimay, R., Yeh, P., and Kuwana, T. (1977) Evaluation of Mediator-Titrants for the Indirect Coulometric Titration of Biocomponents. **38**, 143-169
8. Bazil, J. N. (2017) Analysis of a Functional Dimer Model of Ubiquinol Cytochrome c Oxidoreductase. *Biophys J* **113**, 1599-1612
9. Squire, W., and G. Trapp. (1998) Using complex variables to estimate derivatives of real functions. *Siam Rev* **40**, 110-112
10. Grivennikova, V. G., Kareyeva, A. V., and Vinogradov, A. D. (2018) Oxygen-dependence of mitochondrial ROS production as detected by Amplex Red assay. *Redox Biol* **17**, 192-199
11. Quinlan, C. L., Orr, A. L., Perevoshchikova, I. V., Treberg, J. R., Ackrell, B. A., and Brand, M. D. (2012) Mitochondrial complex II can generate reactive oxygen species at high rates in both the forward and reverse reactions. *J Biol Chem* **287**, 27255-27264
12. Malyala, S., Zhang, Y., Strubbe, J. O., and Bazil, J. N. (2019) Calcium phosphate precipitation inhibits mitochondrial energy metabolism. *PLoS Comput Biol* **15**, e1006719
