## Supplementary material for "Analysis of Mammalian Succinate Dehydrogenase Kinetics and Reactive Oxygen Species Production": Source code: Generate_Figures.m

```
%% Plot Model Outputs
% inputs: 
% data - data structure
% p - parameter structure
% load SDH_data and SDH_params for data and p structures, respectively
% output
% figures used in main text and supplemental material, note that some
% legends and axes labels were done in power point
%
% Copyright:
% Jason Bazil November 2019
% Bazil Laboratory
% Department of Physiology
% Michigan State University

function Generate_Figures(data,p)

% Set model
fun = @main_complexII; 

fittingfunctions = {@S1968_F1A1,@V1979_F21,@V1979_F71,@H2013_F3A,@H2013_F3B,@H2013_F3C,...
                    @S2013_F1A1,@S2013_F1B,@S2013_F1C,@S2013_F1D,@S2013_F2,...
                    @S2013_F3A,@S2013_F3B,@S2013_F4B,@S2013_F4C,@G2017_F2,@G2017_F5,@G2017_F6B};

model = getoutput(fun,p,fittingfunctions);

colors=[0 0.447 0.741; 0.85 0.325 0.098; 0.929 0.694 0.125; 0.494 0.184 0.556; 0.466 0.674 0.188];

%% FIGURE 2

% Kinetics of Succinate Oxidation: Slatter Fig 1A, Grivennikova Fig 6B, Vinogradov Figs 2 & 7
% Panel A: SLATTER, 1968, Fig 1A

im=[find(strcmp(model.label,'Slater_Fig1Aset1a.txt')) ...
    find(strcmp(model.label,'Slater_Fig1Aset1b.txt')) ...
    find(strcmp(model.label,'Slater_Fig1Aset1c.txt')) ...
    find(strcmp(model.label,'Slater_Fig1Aset1d.txt')) ...
    find(strcmp(model.label,'Slater_Fig1Aset1e.txt'))];

id=[find(strcmp(data.label,'Slater_Fig1Aset1a.txt')) ...
    find(strcmp(data.label,'Slater_Fig1Aset1b.txt')) ...
    find(strcmp(data.label,'Slater_Fig1Aset1c.txt')) ...
    find(strcmp(data.label,'Slater_Fig1Aset1d.txt')) ...
    find(strcmp(data.label,'Slater_Fig1Aset1e.txt'))];

figure
hold on
box on
for i=1:numel(id)
    plot(data.Y{id(i)}(:,1),data.Y{id(i)}(:,2),'o','color',colors(i,:),'markersize',30,'linewidth',4)
    plot(model.Y{im(i)}(:,1),model.Y{im(i)}(:,2),'color',colors(i,:),'linewidth',8)
end
ylabel({'Succinate Oxidation Rate';'(\Delta A_{600}/min)'});
xlabel({'Succinate (mM)'})
xlim([0 2.5])
ylim([0 1])
set(gca,'ytick',[0 0.25 0.50 0.75 1.00])
set(gcf,'color','none'),set(gca,'color','none'),set(gca,'FontName','Georgia','FontSize',50);
set(gcf, 'Position', [1 1 1920 1080]); % Maximize figure.

% Panel B: GRIVENNIKOVA, 2017, Fig 6B
% Succinate dehydrogenase activity of SQR using PMS assay

im=[find(strcmp(model.label,'Grivennikova17_fig6Bset1a.txt')) ...
    find(strcmp(model.label,'Grivennikova17_fig6Bset1b.txt')) ...
    find(strcmp(model.label,'Grivennikova17_fig6Bset1c.txt'))];

id=[find(strcmp(data.label,'Grivennikova17_fig6Bset1a.txt')) ...
    find(strcmp(data.label,'Grivennikova17_fig6Bset1b.txt')) ...
    find(strcmp(data.label,'Grivennikova17_fig6Bset1c.txt'))];

figure
hold on
box on
for i=1:numel(id)
plot(data.Y{id(i)}(:,1),data.Y{id(i)}(:,2),'o','color',colors(i,:),'markersize',30,'linewidth',4)
plot(model.Y{im(i)}(:,1),model.Y{im(i)}(:,2),'color',colors(i,:),'linewidth',8)
end
ylabel({'Succinate Oxidation Rate';'(\mumol/min/mg)'});
xlabel({'Succinate ( mM)'})
ylim([0 30])
set(gcf,'color','none'),set(gca,'color','none'),set(gca,'FontName','Georgia','FontSize',50);
set(gcf, 'Position', [1 1 1920 1080]); % Maximize figure.

% Panel C: VINOGRADOV, 1979, Fig 2  
% Effect of varying succinate (x axis) and WB (different curves)
% concentrations on succinate oxidation rate by the soluble succinate
% dehydrogenase. The WB concentrations are 0.05 mM (set 1a), 0.066 mM (set 1b),
% 0.1 mM (set 1c) and 0.33 mM (set 1d).

im=[find(strcmp(model.label,'Vinog_Fig2Aset1a.txt')) ...
    find(strcmp(model.label,'Vinog_Fig2Aset1b.txt')) ...
    find(strcmp(model.label,'Vinog_Fig2Aset1c.txt')) ...
    find(strcmp(model.label,'Vinog_Fig2Aset1d.txt'))];

id=[find(strcmp(data.label,'Vinog_Fig2Aset1a.txt')) ...
    find(strcmp(data.label,'Vinog_Fig2Aset1b.txt')) ...
    find(strcmp(data.label,'Vinog_Fig2Aset1c.txt')) ...
    find(strcmp(data.label,'Vinog_Fig2Aset1d.txt'))];

figure
hold on
box on
for i=1:numel(id)
    plot(data.Y{id(i)}(:,1),data.Y{id(i)}(:,2),'o','color',colors(i,:),'markersize',30,'linewidth',4)
    plot(model.Y{im(i)}(:,1),model.Y{im(i)}(:,2),'color',colors(i,:),'linewidth',8)
end

set(gcf, 'Position', get(0,'Screensize')); % Maximize figure.
ylabel({'Succinate Oxidation Rate';'(\mumol/min/mg)'});
xlabel({'Succinate ( mM)'})
xlim([0 6])
ylim([0 10])
set(gcf,'color','none'),set(gca,'color','none'),set(gca,'FontName','Georgia','FontSize',50);
set(gcf, 'Position', [1 1 1920 1080]); % Maximize figure.

% Panel D: VINOGRADOV, 1979, Fig 7
% Effect of varying pH (x axis) and WB concentrations (different curves) on succinate oxidation rate by the soluble
% succinate dehydrogenase. The WB concentrations are 0.05 mM (set 1a),
% 0.066 mM (set 1b), 0.1 mM (set 1c), 0.2 mM (set 1d) and infinite concentrations (set 1e)

im=fliplr([find(strcmp(model.label,'Vinog_Fig7Aset1a.txt')) ...
           find(strcmp(model.label,'Vinog_Fig7Aset1b.txt')) ...
           find(strcmp(model.label,'Vinog_Fig7Aset1c.txt')) ...
           find(strcmp(model.label,'Vinog_Fig7Aset1d.txt')) ...
           find(strcmp(model.label,'Vinog_Fig7Aset1e.txt'))]);
       
id=fliplr([find(strcmp(data.label,'Vinog_Fig7Aset1a.txt')) ...
           find(strcmp(data.label,'Vinog_Fig7Aset1b.txt')) ...
           find(strcmp(data.label,'Vinog_Fig7Aset1c.txt')) ...
           find(strcmp(data.label,'Vinog_Fig7Aset1d.txt')) ...
           find(strcmp(data.label,'Vinog_Fig7Aset1e.txt'))]);

figure
hold on
box on
for i=1:numel(id)
    plot(data.Y{id(i)}(:,1),data.Y{id(i)}(:,2),'o','color',colors(i,:),'markersize',30,'linewidth',4)
    plot(model.Y{im(i)}(:,1),model.Y{im(i)}(:,2),'color',colors(i,:),'linewidth',8)
end

set(gcf, 'Position', get(0,'Screensize')); % Maximize figure.
ylabel({'Succinate Oxidation Rate';'(\mumol/min/mg)'});
xlabel({'pH'})
xlim([6 10])
ylim([0 20])
set(gcf,'color','none'),set(gca,'color','none'),set(gca,'FontName','Georgia','FontSize',50);
set(gcf, 'Position', [1 1 1920 1080]); % Maximize figure.

%% FIGURE 3 includes Hirst Figs 3A, 3B and 3C and Siebels Figs 1C, 4B and 4C (succinate oxidase activity only)
% Succinate oxidation kinetics in the absence and presence of inhibitors
% atpenin and malonate using bovine SMP

figure
% Panel A: Hirst, Fig 3A
% Succinate concentrations varied
ax1=subplot(2,3,1);
im=[find(strcmp(model.label,'Hirst_Fig3Aset1a.txt')) ...
    find(strcmp(model.label,'Hirst_Fig3Bset1a.txt')) ...
    find(strcmp(model.label,'Hirst_Fig3Cset1a.txt'))];

id=[find(strcmp(data.label,'Hirst_Fig3Aset1a.txt')) ...
    find(strcmp(data.label,'Hirst_Fig3Bset1a.txt')) ...
    find(strcmp(data.label,'Hirst_Fig3Cset1a.txt'))];

hold on
box on
i=1;
plot(data.Y{id(i)}(:,1),data.Y{id(i)}(:,2),'o','color',colors(i,:),'markersize',15,'linewidth',3)
plot(model.Y{im(i)}(:,1),model.Y{im(i)}(:,2),'color',colors(i,:),'linewidth',4)
set(gcf, 'Position', get(0,'Screensize')); % Maximize figure.
ylabel({' Succinate Oxidation Rate'; '(\mumol/min/mg)'});
xlabel({'Succinate (mM)'})
yticks(ax1,[0 .5 1 1.5])
yticklabels(ax1,{'0','0.5','1.0','1.5'})
xticks(ax1,[0 5 10 15 20])
xlim([0 20])
ylim([0 1.5])
set(gcf,'color','none'),set(gca,'color','none'),set(gca,'FontSize',26,'FontName','Times New Roman')

% Panel B: Hirst, Fig 3B
% Atpenin concentrations varied 
ax2=subplot(2,3,2);
hold on
box on
i=2;
plot(data.Y{id(i)}(:,1),data.Y{id(i)}(:,2),'o','color',colors(1,:),'markersize',15,'linewidth',3)
plot(model.Y{im(i)}(:,1),model.Y{im(i)}(:,2),'color',colors(1,:),'linewidth',4)
set(gcf, 'Position', get(0,'Screensize')); % Maximize figure.
xlim([0 100])
ylim([0 110])
xticks(ax2,[0 25 50 75 100])
yticks(ax2,[0 25 50 75 100])
ylabel({'% Succinate Oxidase'});
xlabel({'Atpenin (nM)'})
set(gcf,'color','none'),set(gca,'color','none'),set(gca,'FontSize',26,'FontName','Times New Roman')

% Panel C: Hirst, Fig 3C
% Malonate concentrations varied 
ax3=subplot(2,3,3);
hold on
box on
i=3;
plot(data.Y{id(i)}(:,1),data.Y{id(i)}(:,2),'o','color',colors(1,:),'markersize',15,'linewidth',3)
plot(model.Y{im(i)}(:,1),model.Y{im(i)}(:,2),'color',colors(1,:),'linewidth',4)
set(gcf, 'Position', get(0,'Screensize')); % Maximize figure.
xlim([0 5])
ylim([0 110])
xticks(ax3,[0 1 2 3 4 5])
yticks(ax3,[0 25 50 75 100])
ylabel({'% Succinate Oxidase'});
xlabel({'Malonate (mM)'})
set(gcf,'color','none'),set(gca,'color','none'),set(gca,'FontSize',26,'FontName','Times New Roman')

% Panel D: Siebels, Fig 1C
% Atpenin A5 concentrations varied 
im=[find(strcmp(model.label,'Siebels13_fig1Cset1a.txt')) ...
    find(strcmp(model.label,'Siebels13_fig1Cset2a.txt'))];

id=[find(strcmp(data.label,'Siebels13_fig1Cset1a.txt')) ...
    find(strcmp(data.label,'Siebels13_fig1Cset2a.txt'))];

ax4=subplot(2,3,4);
hold on
box on
i=2;
plot(data.Y{id(i)}(:,1),data.Y{id(i)}(:,2),'o','color',colors(i,:),'markersize',15,'linewidth',4)
plot(model.Y{im(i)}(:,1),model.Y{im(i)}(:,2),'color',colors(i,:),'linewidth',4)
xlim([0 250])
ylim([0 110])
xticks(ax4,[0 50 100 150 200 250])
yticks(ax4,[0 25 50 75 100])
set(gca,'FontSize',26,'FontName','Times New Roman')
ylabel({'% Succinate Oxidase'});
xlabel({'Atpenin (nM)'})
set(gcf,'color','none'),set(gca,'color','none')

% Panel E: Siebels, Fig 4B
% Malate concentrations varied 
ax5=subplot(2,3,5);
im=[find(strcmp(model.label,'Siebels13_fig4Bset1a.txt')) ...
    find(strcmp(model.label,'Siebels13_fig4Bset2a.txt'))];

id=[find(strcmp(data.label,'Siebels13_fig4Bset1a.txt')) ...
    find(strcmp(data.label,'Siebels13_fig4Bset2a.txt'))];

hold on
box on
i=2;
set(gcf, 'Position', get(0,'Screensize')); % Maximize figure.
plot(data.Y{id(i)}(:,1),data.Y{id(i)}(:,2),'o','color',colors(i,:),'markersize',15,'linewidth',4)
plot(model.Y{im(i)}(:,1),model.Y{im(i)}(:,2),'color',colors(i,:),'linewidth',4)
ylabel({'% Succinate Oxidase'});
xlabel({'Malate (mM)'})
xlim([0 10])
ylim([0 110])
xticks(ax5,[0 2 4 6 8 10])
yticks(ax5,[0 25 50 75 100])
set(gca,'FontSize',26,'FontName','Times New Roman')
set(gcf,'color','none'),set(gca,'color','none')

% Panel F: Siebels, Fig 4C
% Oxaloacetate concentrations varied 
ax6=subplot(2,3,6);
im=[find(strcmp(model.label,'Siebels13_fig4Cset1a.txt')) ...
    find(strcmp(model.label,'Siebels13_fig4Cset2a.txt'))];

id=[find(strcmp(data.label,'Siebels13_fig4Cset1a.txt')) ...
    find(strcmp(data.label,'Siebels13_fig4Cset2a.txt'))];
hold on
box on
i=2;
set(gcf, 'Position', get(0,'Screensize')); % Maximize figure.
plot(data.Y{id(i)}(:,1),data.Y{id(i)}(:,2),'o','color',colors(i,:),'markersize',15,'linewidth',4)
plot(model.Y{im(i)}(:,1),model.Y{im(i)}(:,2),'color',colors(i,:),'linewidth',4)
ylabel({'% Succinate Oxidase'});
xlabel({'Oxaloacetate (\muM)'})
xlim([0 30])
ylim([0 110])
xticks(ax6,[0 5 10 15 20 25 30])
yticks(ax6,[0 25 50 75 100])
set(gca,'FontSize',26,'FontName','Times New Roman')
set(gcf,'color','none'),set(gca,'color','none')

%% FIGURE 4: Inhibitor-induced ROS production at varied succinate concentrations 
% These data show that ROS production peaks at sub-saturating [suc], when
% the flavin is reduced and available to interact with oxygen. 

% Panel A: Grivennikova, Fig 2
% JH2O2 in the presence of 10 uM rotenone, 1.6 uM myxothiazol 
figure
im=find(strcmp(model.label,'Grivennikova17_fig2Bset1a.txt'));
id=find(strcmp(data.label,'Grivennikova17_fig2Bset1a.txt'));
hold on
box on
set(gcf, 'Position', get(0,'Screensize')); % Maximize figure.
for i=1:numel(id)
    plot(model.Y{im(i)}(:,1)/1e3,model.Y{im(i)}(:,2)*1e3,'color',colors(i,:),'linewidth',6)
    plot(data.Y{id(i)}(:,1)/1e3,data.Y{id(i)}(:,2)*1e3,'o','color',colors(i,:),'markersize',30,'linewidth',4)
end
ylabel({'J_{ROS} (pmol/min/mg)'});
xlabel({'Succinate (mM)'})
xlim([0 20.5])
ylim([0 1050])
yticks(gca,[0 200 400 600 800 1000])
set(gcf,'color','none'),set(gca,'color','none'),set(gca,'FontSize',54,'FontName','Times New Roman')

% Grivennikova Fig 2A inset plot
figure
im=find(strcmp(model.label,'Grivennikova17_fig2Bset1a.txt'));
id=find(strcmp(data.label,'Grivennikova17_fig2Bset1a.txt'));
hold on
box on
set(gcf, 'Position', get(0,'Screensize')); % Maximize figure.
for i=1:numel(id)
    plot(model.Y{im(i)}(:,1)/1e3,model.Y{im(i)}(:,2)*1e3,'color',colors(i,:),'linewidth',8)
    plot(data.Y{id(i)}(:,1)/1e3,data.Y{id(i)}(:,2)*1e3,'o','color',colors(i,:),'markersize',30,'linewidth',8)
end

xlim([0 0.105])
ylim([0 900])
yticks([0 200 400 600 800])
yticklabels({'0','200','400','600','800'})
set(gcf,'color','none'),set(gca,'color','none'),set(gca,'FontSize',54,'FontName','Times New Roman')

% Panel C: Grivennikova H2O2 and SO specific rates
figure
im=find(strcmp(model.label,'Grivennikova17_fig2Bset1a.txt'));
hold on
box on
i=1;
set(gcf, 'Position', get(0,'Screensize')); % Maximize figure.
plot(model.Y{im(i)}(:,1)/1e3,model.g{im(i)}.JFADH_SO*1e12.*60/0.1/1e3/2,'color',colors(1,:),'linewidth',6)
plot(model.Y{im(i)}(:,1)/1e3,model.g{im(i)}.JISC3_SO*1e12.*60/0.1/1e3/2,'color',colors(2,:),'linewidth',6)
plot(model.Y{im(i)}(:,1)/1e3,model.g{im(i)}.JFADH2_H2O2*1e12.*60/0.1/1e3,'color',colors(3,:),'linewidth',6)
ylabel({'J_{e^-} (pmol/min/mg)'});
xlabel({'Succinate (mM)'})
xlim([0 20.5])
ylim([0 700])
yticks([0 200 400 600 800])
set(gcf,'color','none'),set(gca,'color','none'),set(gca,'FontSize',54,'FontName','Times New Roman')

% Panel B: Siebels, Fig 1B
% JH2O2 in the presence of 50 nM Atpenin A5

im=find(strcmp(model.label,'Siebels13_fig1Bset1a.txt'));
id=find(strcmp(data.label,'Siebels13_fig1Bset1a.txt'));
figure
hold on
box on
set(gcf, 'Position', get(0,'Screensize')); % Maximize figure.
for i=1:numel(id)
    plot(model.Y{im(i)}(:,1),model.Y{im(i)}(:,2),'color',colors(i,:),'linewidth',6)
    plot(data.Y{id(i)}(:,1),data.Y{id(i)}(:,2),'o','color',colors(i,:),'markersize',30,'linewidth',4)
end
ylabel({'J_{ROS} (pmol/min/mg)'});
xlabel({'Succinate (mM)'})
xlim([0 5.5])
ylim([0 1050])
yticks(gca,[0 200 400 600 800 1000])
set(gcf,'color','none'),set(gca,'color','none'),set(gca,'FontSize',54,'FontName','Times New Roman')

% Panel D: Siebels H2O2 and SO specific rates

im=find(strcmp(model.label,'Siebels13_fig1Bset1a.txt'));
figure
hold on
box on
i=1;
set(gcf, 'Position', get(0,'Screensize')); % Maximize figure.
plot(model.Y{im(i)}(:,1),model.g{im(i)}.JFADH_SO*1e12.*60/.12/1e3/2,'color',colors(1,:),'linewidth',6)
plot(model.Y{im(i)}(:,1),model.g{im(i)}.JISC3_SO*1e12.*60/.12/1e3/2,'color',colors(2,:),'linewidth',6)
plot(model.Y{im(i)}(:,1),model.g{im(i)}.JFADH2_H2O2*1e12.*60/.12/1e3,'color',colors(3,:),'linewidth',6)
ylabel({'J_{e^-} (pmol/min/mg)'});
xlabel({'Succinate (mM)'})
xlim([0 5.5])
ylim([0 800])
yticks([0 200 400 600 800])
set(gcf,'color','none'),set(gca,'color','none'),set(gca,'FontSize',54,'FontName','Times New Roman')

%% FIGURE 5
% The effects of different SQR inhibitors (stigmatellin and atpenin A5) on JH2O2
% at sub-saturating succinate concentrations

% Panel A: Siebels, Fig 1A, 1 uM stigmatellin and 50 nM Atpenin A5
figure
im=[find(strcmp(model.label,'Siebels13_fig1Aset1a.txt')) ...
    find(strcmp(model.label,'Siebels13_fig1Aset1c.txt')) ...
    find(strcmp(model.label,'Siebels13_fig1Dset1a.txt'))];

id=[find(strcmp(data.label,'Siebels13_fig1Aset1a.txt')) ...
    find(strcmp(data.label,'Siebels13_fig1Aset1c.txt')) ...
    find(strcmp(data.label,'Siebels13_fig1Dset1a.txt'))];

hold on
box on
for i=1:numel(id)
    plot(data.Y{id(i)}(:,1),data.Y{id(i)}(:,2),'o','MarkerEdgeColor',colors(i,:),'markersize',30,'linewidth',4)
    plot(model.Y{im(i)}(:,1),model.Y{im(i)}(:,2),'color',colors(i,:),'linewidth',8)
end
set(gcf, 'Position', get(0,'Screensize')); % Maximize figure.
ylabel({'J_{ROS}(pmol/min/mg)'});
xlabel({'Succinate (mM)'})
xlim([0 5.5])
ylim([0 3250])
set(gcf,'color','none'),set(gca,'color','none'),set(gca,'FontSize',54,'FontName','Times New Roman')

figure
hold on
box on
i=1;
plot(model.Y{im(i)}(:,1),model.g{im(i)}.JFADH_SO/2*1e12*60/0.12/1e3,'color',colors(1,:),'linewidth',8)
plot(model.Y{im(i)}(:,1),model.g{im(i)}.JISC3_SO/2*1e12*60/0.12/1e3,'color',colors(2,:),'linewidth',8)
plot(model.Y{im(i)}(:,1),model.g{im(i)}.JFADH2_H2O2*1e12*60/0.12/1e3,'color',colors(3,:),'linewidth',8)
set(gcf, 'Position', get(0,'Screensize')); % Maximize figure.
ylabel({'J_{e^-}(pmol/min/mg)'});
xlabel({'Succinate (mM)'})
xlim([0 5.5])
ylim([0 3250])
set(gcf,'color','none'),set(gca,'color','none'),set(gca,'FontSize',54,'FontName','Times New Roman')

figure
hold on
box on
i=3;
plot(model.Y{im(i)}(:,1),model.g{im(i)}.JFADH_SO/2*1e12*60/0.12/1e3,'color',colors(1,:),'linewidth',8)
plot(model.Y{im(i)}(:,1),model.g{im(i)}.JISC3_SO/2*1e12*60/0.12/1e3,'color',colors(2,:),'linewidth',8)
plot(model.Y{im(i)}(:,1),model.g{im(i)}.JFADH2_H2O2*1e12*60/0.12/1e3,'color',colors(3,:),'linewidth',8)
set(gcf, 'Position', get(0,'Screensize')); % Maximize figure.
ylabel({'J_{e^-}(pmol/min/mg)'});
xlabel({'Succinate (mM)'})
xlim([0 5.5])
ylim([0 2500])
set(gcf,'color','none'),set(gca,'color','none'),set(gca,'FontSize',54,'FontName','Times New Roman')

%% FIGURE 6
% Effects of varying concentrations of SQR and SDH inhibitors on JH2O2
% Succinate is present at 100 uM (peak ROS production rate in Siebels)

% Panel A: Siebels, Fig 1C
% JH2O2 production as a function of [Atpenin A5] (nM)
figure
im=find(strcmp(model.label,'Siebels13_fig1Cset1a.txt'));
id=find(strcmp(data.label,'Siebels13_fig1Cset1a.txt'));
hold on
box on
for i=1:numel(id)
    plot(model.Y{im(i)}(:,1),model.Y{im(i)}(:,2),'color',colors(i,:),'linewidth',5)
    plot(data.Y{id(i)}(:,1),data.Y{id(i)}(:,2),'o','color',colors(i,:),'markersize',30,'linewidth',4)
end
set(gcf, 'Position', [1 1 1360 1080]); % Maximize figure.
ylabel({'J_{H_2O_2} (pmol/min/mg)'});
xlabel({'Atpenin (nM)'})
xlim([0 250])
ylim([0 1600])
xticks(gca,[0 50 100 150 200 250])
set(gcf,'color','none'),set(gca,'color','none'),set(gca,'FontSize',54,'FontName','Times New Roman')

% Panel B: Siebels, Fig 4B
% JH2O2 production as a function of [malate] (mM)
figure
im=find(strcmp(model.label,'Siebels13_fig4Bset1a.txt'));
id=find(strcmp(data.label,'Siebels13_fig4Bset1a.txt'));
hold on
box on
for i=1:numel(id)
    plot(model.Y{im(i)}(:,1),model.Y{im(i)}(:,2),'color',colors(i,:),'linewidth',5)
    plot(data.Y{id(i)}(:,1),data.Y{id(i)}(:,2),'o','color',colors(i,:),'markersize',30,'linewidth',4)
end
set(gcf, 'Position', [1 1 1360 1080]); % Maximize figure.
ylabel({'J_{H_2O_2} (pmol/min/mg)'});
xlabel({'Malate (mM)'})
xlim([0 6])
ylim([0 1600])
set(gcf,'color','none'),set(gca,'color','none'),set(gca,'FontSize',54,'FontName','Times New Roman')

% Panel C: Siebels, Fig 4C
% JH2O2 production as a function of [oxaloacetate] (uM)
figure
im=find(strcmp(model.label,'Siebels13_fig4Cset1a.txt'));
id=find(strcmp(data.label,'Siebels13_fig4Cset1a.txt'));
hold on
box on
for i=1:numel(id)
    plot(model.Y{im(i)}(:,1),model.Y{im(i)}(:,2),'color',colors(i,:),'linewidth',5)
    plot(data.Y{id(i)}(:,1),data.Y{id(i)}(:,2),'o','color',colors(i,:),'markersize',30,'linewidth',4)
end
set(gcf, 'Position', [1 1 1360 1080]); % Maximize figure.
ylabel({'J_{H_2O_2} (pmol/min/mg)'});
xlabel({'Oxaloacetate (\muM)'})
xlim([0 30])
ylim([0 1600])
set(gcf,'color','none'),set(gca,'color','none'),set(gca,'FontSize',54,'FontName','Times New Roman')

%% FIGURE 7
% Effects of varying FUM/SUC ratios on atpenin-induced or stigmatellin-induced ROS production

% Panel A: Grivennikova, Fig 5
% Bovine heart SMP (0.1 mg/mL) prepared in the presence of 10 uM rotenone and 1.6 uM myxothiazol
% Atpenin A5 present at 1 uM 
% Succinate present at 2 or 5 uM 

im=[find(strcmp(model.label,'Grivennikova17_fig5set1a.txt')) ...
    find(strcmp(model.label,'Grivennikova17_fig5set1b.txt'))];

id=[find(strcmp(data.label,'Grivennikova17_fig5set1a.txt')) ...
    find(strcmp(data.label,'Grivennikova17_fig5set1b.txt'))];

figure
hold on
box on
for i=1:numel(id)
    plot(data.Y{id(i)}(:,1),data.Y{id(i)}(:,2)*1e3,'o','color',colors(i,:),'markersize',30,'linewidth',4)
    plot(model.Y{im(i)}(:,1),model.Y{im(i)}(:,2)*1e3,'color',colors(i,:),'linewidth',8)
end
set(gcf, 'Position', get(0,'Screensize')); % Maximize figure.
xlim([0 60])
ylim([0 1000])
ylabel({'J_{O_2^{.-}} (pmol/min/mg)'});
xlabel({'[Fumarate] (\muM)'})
set(gcf,'color','none'),set(gca,'color','none'),set(gca,'FontSize',54,'FontName','Times New Roman')

% site-specific ROS production rates
figure
hold on
box on
i=1; % 2 uM succinate condition
plot(model.Y{im(i)}(:,1),model.g{im(i)}.JFADH_SO*1e12*60/0.1/1e3,'color',colors(1,:),'linewidth',8)
plot(model.Y{im(i)}(:,1),model.g{im(i)}.JISC3_SO*1e12*60/0.1/1e3,'color',colors(2,:),'linewidth',8)
plot(model.Y{im(i)}(:,1),model.g{im(i)}.JFADH2_H2O2*1e12*60/0.1/1e3*2,'color',colors(3,:),'linewidth',8)
set(gcf, 'Position', get(0,'Screensize')); % Maximize figure.
xlim([0 60])
ylim([0 1000])
ylabel({'J_{e^-}(pmol/min/mg)'});
xlabel({'[Fumarate] (mM)'})
set(gcf,'color','none'),set(gca,'color','none'),set(gca,'FontSize',54,'FontName','Times New Roman')

% Panel B: Siebels, Fig 3B
% Bovine heart SMP (0.12 mg/mL) prepared in the presence of 10 uM rotenone and 1.6 uM myxothiazol
% Stigmatellin present at 1 uM 
% Succinate (mM): 0.1, 0.5 or 1.0 
% Fumarate (mM): 0, 0.1, 1.0 or 5.0

im=[find(strcmp(model.label,'Siebels13_fig3Bset2a.txt')) ...
    find(strcmp(model.label,'Siebels13_fig3Bset2b.txt')) ...
    find(strcmp(model.label,'Siebels13_fig3Bset2c.txt'))];

id=[find(strcmp(data.label,'Siebels13_fig3Bset2a.txt')) ...
    find(strcmp(data.label,'Siebels13_fig3Bset2b.txt')) ...
    find(strcmp(data.label,'Siebels13_fig3Bset2c.txt'))];

figure
hold on
box on
for i=1:numel(id)
    plot(data.Y{id(i)}(:,1),data.Y{id(i)}(:,2),'o','color',colors(i,:),'markersize',30,'linewidth',4)
    plot(model.Y{im(i)}(:,1),model.Y{im(i)}(:,2),'color',colors(i,:),'linewidth',8)
end
set(gcf, 'Position', get(0,'Screensize')); % Maximize figure.
xlim([0 6])
ylim([0 3000])
ylabel({'J_{ROS} (pmol/min/mg)'});
xlabel({'[Fumarate] (mM)'})
set(gcf,'color','none'),set(gca,'color','none'),set(gca,'FontSize',54,'FontName','Times New Roman')

% Inset
figure 
hold on
box on
for i=1:numel(id)
    plot(data.Y{id(i)}(:,1),data.Y{id(i)}(:,2),'o','color',colors(i,:),'markersize',30,'linewidth',4)
    plot(model.Y{im(i)}(:,1),model.Y{im(i)}(:,2),'color',colors(i,:),'linewidth',8)
end
set(gcf, 'Position', get(0,'Screensize')); % Maximize figure.
xlim([0 1.05])
ylim([0 3000])
set(gcf,'color','none'),set(gca,'color','none'),set(gca,'FontSize',54,'FontName','Times New Roman')

% site-specific ROS production sites
figure
hold on
box on
i=1; % 0.1 mM succinate condition
plot(model.Y{im(i)}(:,1),model.g{im(i)}.JFADH_SO*1e12*60/0.1/1e3/2,'color',colors(1,:),'linewidth',8)
plot(model.Y{im(i)}(:,1),model.g{im(i)}.JISC3_SO*1e12*60/0.1/1e3/2,'color',colors(2,:),'linewidth',8)
plot(model.Y{im(i)}(:,1),model.g{im(i)}.JFADH2_H2O2*1e12*60/0.1/1e3,'color',colors(3,:),'linewidth',8)
set(gcf, 'Position', get(0,'Screensize')); % Maximize figure.
xlim([0 6])
ylim([0 3000])
ylabel({'J_{e^-}(pmol/min/mg)'});
xlabel({'[Fumarate] (mM)'})
set(gcf,'color','none'),set(gca,'color','none'),set(gca,'FontSize',54,'FontName','Times New Roman')

% Panel C: Siebels, Fig 3A
% Bovine heart SMP (0.12 mg/mL) prepared in the presence of 10 uM rotenone and 1.6 uM myxothiazol
% Atpenin A5 present at 50 nM 
% Succinate (mM): 0.1, 0.5 or 1.0 
% Fumarate (mM): 0, 0.1, 1.0 or 5.0

im=[find(strcmp(model.label,'Siebels13_fig3Aset1a.txt')) ...
    find(strcmp(model.label,'Siebels13_fig3Aset1b.txt')) ...
    find(strcmp(model.label,'Siebels13_fig3Aset1c.txt'))];

id=[find(strcmp(data.label,'Siebels13_fig3Aset1a.txt')) ...
    find(strcmp(data.label,'Siebels13_fig3Aset1b.txt')) ...
    find(strcmp(data.label,'Siebels13_fig3Aset1c.txt'))];

figure
hold on
box on
for i=1:numel(id)
    plot(data.Y{id(i)}(:,1),data.Y{id(i)}(:,2),'o','color',colors(i,:),'markersize',30,'linewidth',4)
    plot(model.Y{im(i)}(:,1),model.Y{im(i)}(:,2),'color',colors(i,:),'linewidth',8)
end
set(gcf, 'Position', get(0,'Screensize')); % Maximize figure.
xlim([0 6])
ylim([0 1000])
ylabel({'J_{ROS} (pmol/min/mg)'});
xlabel({'[Fumarate] (mM)'})
set(gcf,'color','none'),set(gca,'color','none'),set(gca,'FontSize',54,'FontName','Times New Roman')

% Inset 
figure
hold on
box on
for i=1:numel(id)
    plot(data.Y{id(i)}(:,1),data.Y{id(i)}(:,2),'o','color',colors(i,:),'markersize',30,'linewidth',4)
    plot(model.Y{im(i)}(:,1),model.Y{im(i)}(:,2),'color',colors(i,:),'linewidth',8)
end
set(gcf, 'Position', get(0,'Screensize')); % Maximize figure.
xlim([0 1.05])
ylim([0 1000])
set(gcf,'color','none'),set(gca,'color','none'),set(gca,'FontSize',54,'FontName','Times New Roman')

% site-specific ROS production sites
figure
hold on
box on
i=1; % 0.1 mM succinate condition
plot(model.Y{im(i)}(:,1),model.g{im(i)}.JFADH_SO*1e12*60/0.1/1e3/2,'color',colors(1,:),'linewidth',8)
plot(model.Y{im(i)}(:,1),model.g{im(i)}.JISC3_SO*1e12*60/0.1/1e3/2,'color',colors(2,:),'linewidth',8)
plot(model.Y{im(i)}(:,1),model.g{im(i)}.JFADH2_H2O2*1e12*60/0.1/1e3,'color',colors(3,:),'linewidth',8)
set(gcf, 'Position', get(0,'Screensize')); % Maximize figure.
xlim([0 6])
ylim([0 1000])
ylabel({'J_{e^-}(pmol/min/mg)'});
xlabel({'[Fumarate] (mM)'})
set(gcf,'color','none'),set(gca,'color','none'),set(gca,'FontSize',54,'FontName','Times New Roman')

%% FIGURE 8
% Panel A: Siebels, Fig 2, Stigmatellin
im=[find(strcmp(model.label,'Siebels13_fig2set1a.txt')) find(strcmp(model.label,'Siebels13_fig2set2a.txt'))];
id=[find(strcmp(data.label,'Siebels13_fig2set1a.txt')) find(strcmp(data.label,'Siebels13_fig2set2a.txt'))];

figure
hold on
box on
for i=1:numel(id)
    plot(data.Y{id(i)}(:,1),data.Y{id(i)}(:,2),'o','MarkerEdgeColor',colors(i,:),'markersize',30,'linewidth',4)
    plot(model.Y{im(i)}(:,1),model.Y{im(i)}(:,2),'color',colors(i,:),'linewidth',8)
end
xlim([7 8.5])
ylim([0 10000])
ylabel({'J_{ROS} (pmol e^- eq/min/mg)'});
xlabel({'pH'})
xtickformat('%.1f')
set(gcf,'color','none'),set(gca,'color','none'),set(gca,'FontName','Georgia','FontSize',50);
set(gcf, 'Position', [1 1 1920 1080]); % Maximize figure.

% site-specific ROS production sites
figure
hold on
box on
i=1; % 0.1 mM succinate condition
plot(model.Y{im(i)}(:,1),model.g{im(i)}.JFADH_SO*1e12*60/0.12/1e3,'color',colors(1,:),'linewidth',8)
plot(model.Y{im(i)}(:,1),model.g{im(i)}.JISC3_SO*1e12*60/0.12/1e3,'color',colors(2,:),'linewidth',8)
plot(model.Y{im(i)}(:,1),model.g{im(i)}.JFADH2_H2O2*1e12*60/0.12/1e3*2,'color',colors(3,:),'linewidth',8)
set(gcf, 'Position', get(0,'Screensize')); % Maximize figure.
xlim([7 8.5])
ylim([0 8000])
ylabel({'J_{e^-} (pmol e^- eq/min/mg)'});
xlabel({'pH'})
xtickformat('%.1f')
set(gcf,'color','none'),set(gca,'color','none'),set(gca,'FontSize',54,'FontName','Times New Roman')

% Panel B: Siebels, Fig 2, Atpenin A5
im=[find(strcmp(model.label,'Siebels13_fig2set1c.txt')) find(strcmp(model.label,'Siebels13_fig2set2c.txt'))];
id=[find(strcmp(data.label,'Siebels13_fig2set1c.txt')) find(strcmp(data.label,'Siebels13_fig2set2c.txt'))];

figure
hold on
box on
for i=1:numel(id)
    plot(data.Y{id(i)}(:,1),data.Y{id(i)}(:,2),'o','MarkerEdgeColor',colors(i,:),'markersize',30,'linewidth',4)
    plot(model.Y{im(i)}(:,1),model.Y{im(i)}(:,2),'color',colors(i,:),'linewidth',8)
end
set(gcf, 'Position', get(0,'Screensize')); % Maximize figure.
xlim([7 8.5])
ylim([0 4000])
ylabel({'J_{ROS} (pmol e^- eq/min/mg)'});
xlabel({'pH'})
% xtickformat('%.1f')
set(gcf,'color','none'),set(gca,'color','none'),set(gca,'FontName','Georgia','FontSize',50);
set(gcf, 'Position', [1 1 1920 1080]); % Maximize figure.

figure
hold on
box on
i=1; % 0.1 mM succinate condition
plot(model.Y{im(i)}(:,1),model.g{im(i)}.JFADH_SO*1e12*60/0.12/1e3,'color',colors(1,:),'linewidth',8)
plot(model.Y{im(i)}(:,1),model.g{im(i)}.JISC3_SO*1e12*60/0.12/1e3,'color',colors(2,:),'linewidth',8)
plot(model.Y{im(i)}(:,1),model.g{im(i)}.JFADH2_H2O2*1e12*60/0.12/1e3*2,'color',colors(3,:),'linewidth',8)
set(gcf, 'Position', get(0,'Screensize')); % Maximize figure.
xlim([7 8.5])
ylim([0 8000])
ylabel({'J_{e^-} (pmol e^- eq/min/mg)'});
xlabel({'pH'})
xtickformat('%.1f')
set(gcf,'color','none'),set(gca,'color','none'),set(gca,'FontSize',54,'FontName','Times New Roman')


%% ++++++++++++++++++++++++++++++++++++++++++++++++++++++++++++++++++
%                                 SDH mesh
%  ++++++++++++++++++++++++++++++++++++++++++++++++++++++++++++++++++

% experimental details
p.T                 = 273.15 + 37;                   % 37 C
p.Etot              = 1;                             % TN
p.atpenin           = 0;                             % M

Succinate           = linspace(0,20,101)*1e-3;       % M
Fumarate            = 20e-3-Succinate;

Q                   = linspace(0,20,201)*1e-3;       % M
QH2                 = 20e-3 - Q;                     % M

x = zeros(numel(fieldnames(p.in)),numel(Succinate)); 
x(p.in.O2,:)        = 20e-6;                         % M
x(p.in.Succinate,:) = Succinate;                     % M
x(p.in.Fumarate,:)  = Fumarate;                      % M
x(p.in.H,:)         = 10^(-7);

for i=1:numel(Q)
    x(p.in.QH2,:) = QH2(i);
    x(p.in.Q,:) =  Q(i);                             %M
    [J(i,:),JSO(i,:),JH2O2(i,:),JQH2(i,:),~,details{i}] = main_complexII(x,p);
    
    E0(i,:) = details{i}.E0;
    E1(i,:) = details{i}.E1;
    E2(i,:) = details{i}.E2;
    E3(i,:) = details{i}.E3;
    E4(i,:) = details{i}.E4;
    
    JISC3_SO(i,:) = details{i}.JISC3_SO;
    JFADH_SO(i,:) = details{i}.JFADH_SO;
    
end

[X,Y]=meshgrid(Succinate./Fumarate,QH2./Q);

%%
figure
mesh(X,Y,J)
set(gca,'XScale','log')
set(gca,'YScale','log')
xlabel('SUC/FUM')
ylabel('QH_2/Q')
zlabel('SUC TN (s^{-1})')
set(gca,'FontSize',40,'FontName','Times New Roman','fontweight','bold')
set(gcf,'color','none'),set(gca,'color','none')
zlim([-125 50])
set(gcf, 'Position', [1 1 1920 1080]); % Maximize figure.
set(gca,'XMinorTick','on','XScale','log','XTick',[0.01 0.1 1 10 100],...
    'XTickLabel',{'0.01','0.1','0','10','100'},'YMinorTick','on','YScale','log',...
    'YTick',[0.01 0.1 1 10 100],'YTickLabel',{'0.01','0.1','1','10','100'});
view([-65 35])

figure
mesh(X,Y,(JSO/2 + JH2O2))
set(gca,'XScale','log')
set(gca,'YScale','log')
xlabel('SUC/FUM')
ylabel('QH_2/Q')
zlabel('ROS (2e^-) TN (s^{-1})')
set(gca,'FontSize',40,'FontName','Times New Roman','fontweight','bold')
set(gcf,'color','none'),set(gca,'color','none')
set(gcf, 'Position', [1 1 1920 1080]); % Maximize figure.
set(gca,'XMinorTick','on','XScale','log','XTick',[0.01 0.1 1 10 100],...
    'XTickLabel',{'0.01','0.1','0','10','100'},'YMinorTick','on','YScale','log',...
    'YTick',[0.01 0.1 1 10 100],'YTickLabel',{'0.01','0.1','1','10','100'});
view([-65 35])

% Superoxide from flavin 
figure
mesh(X,Y,JFADH_SO)
set(gcf, 'Position', get(0,'Screensize')); % Maximize figure.
set(gca,'XScale','log')
set(gca,'YScale','log')
xlabel('SUC/FUM')
ylabel('QH_2/Q')
zlabel('TN^{FADH^.}_{SO} (s^{-1})')
set(gca,'FontSize',40,'FontName','Times New Roman','fontweight','bold')
set(gcf,'color','none'),set(gca,'color','none')
set(gcf, 'Position', [1 1 1920 1080]); % Maximize figure.
set(gca,'XMinorTick','on','XScale','log','XTick',[0.01 0.1 1 10 100],...
    'XTickLabel',{'0.01','0.1','0','10','100'},'YMinorTick','on','YScale','log',...
    'YTick',[0.01 0.1 1 10 100],'YTickLabel',{'0.01','0.1','1','10','100'});
view([-65 35])

% Superoxide from ISC3
figure
mesh(X,Y,JISC3_SO)
set(gcf, 'Position', get(0,'Screensize')); % Maximize figure.
set(gca,'XScale','log')
set(gca,'YScale','log')
xlabel('SUC/FUM')
ylabel('QH_2/Q')
zlabel('TN^{3Fe-4S}_{SO} (s^{-1})')
set(gca,'FontSize',40,'FontName','Times New Roman','fontweight','bold')
set(gcf,'color','none'),set(gca,'color','none')
set(gcf, 'Position', [1 1 1920 1080]); % Maximize figure.
set(gca,'XMinorTick','on','XScale','log','XTick',[0.01 0.1 1 10 100],...
    'XTickLabel',{'0.01','0.1','0','10','100'},'YMinorTick','on','YScale','log',...
    'YTick',[0.01 0.1 1 10 100],'YTickLabel',{'0.01','0.1','1','10','100'});
view([-65 35])

figure
mesh(X,Y,E0)
set(gca,'XScale','log')
set(gca,'YScale','log')
xlabel('SUC/FUM')
ylabel('QH_2/Q')
zlabel('E0')
ztickformat('%.1f')
set(gca,'FontSize',30,'FontName','Times New Roman')
set(gcf,'color','none'),set(gca,'color','none')
set(gcf, 'Position', [1 1 1080 1080]); % Maximize figure.
set(gca,'XMinorTick','on','XScale','log','XTick',[0.01 0.1 1 10 100],...
    'XTickLabel',{'0.01','0.1','0','10','100'},'YMinorTick','on','YScale','log',...
    'YTick',[0.01 0.1 1 10 100],'YTickLabel',{'0.01','0.1','1','10','100'});
view([-65 35])

figure
mesh(X,Y,E1)
set(gca,'XScale','log')
set(gca,'YScale','log')
xlabel('SUC/FUM')
ylabel('QH_2/Q')
zlabel('E1')
ztickformat('%.1f')
set(gca,'FontSize',30,'FontName','Times New Roman')
set(gcf,'color','none'),set(gca,'color','none')
set(gcf, 'Position', [1 1 1080 1080]); % Maximize figure.
set(gca,'XMinorTick','on','XScale','log','XTick',[0.01 0.1 1 10 100],...
    'XTickLabel',{'0.01','0.1','0','10','100'},'YMinorTick','on','YScale','log',...
    'YTick',[0.01 0.1 1 10 100],'YTickLabel',{'0.01','0.1','1','10','100'});
view([-65 35])

figure
mesh(X,Y,E2)
set(gca,'XScale','log')
set(gca,'YScale','log')
xlabel('SUC/FUM')
ylabel('QH_2/Q')
zlabel('E2')
ztickformat('%.1f')
set(gca,'FontSize',30,'FontName','Times New Roman')
set(gcf,'color','none'),set(gca,'color','none')
set(gcf, 'Position', [1 1 1080 1080]); % Maximize figure.
set(gca,'XMinorTick','on','XScale','log','XTick',[0.01 0.1 1 10 100],...
    'XTickLabel',{'0.01','0.1','0','10','100'},'YMinorTick','on','YScale','log',...
    'YTick',[0.01 0.1 1 10 100],'YTickLabel',{'0.01','0.1','1','10','100'});
view([-65 35])

figure
mesh(X,Y,E3)
set(gca,'XScale','log')
set(gca,'YScale','log')
xlabel('SUC/FUM')
ylabel('QH_2/Q')
zlabel('E3')
ztickformat('%.2f')
set(gca,'FontSize',30,'FontName','Times New Roman')
set(gcf,'color','none'),set(gca,'color','none')
set(gcf, 'Position', [1 1 1080 1080]); % Maximize figure.
set(gca,'XMinorTick','on','XScale','log','XTick',[0.01 0.1 1 10 100],...
    'XTickLabel',{'0.01','0.1','0','10','100'},'YMinorTick','on','YScale','log',...
    'YTick',[0.01 0.1 1 10 100],'YTickLabel',{'0.01','0.1','1','10','100'});
view([-65 35])

figure
mesh(X,Y,E4)
set(gca,'XScale','log')
set(gca,'YScale','log')
xlabel('SUC/FUM')
ylabel('QH_2/Q')
zlabel('E4')
ztickformat('%.2f')
set(gca,'FontSize',30,'FontName','Times New Roman')
set(gcf,'color','none'),set(gca,'color','none')
set(gcf, 'Position', [1 1 1080 1080]); % Maximize figure.
set(gca,'XMinorTick','on','XScale','log','XTick',[0.01 0.1 1 10 100],...
    'XTickLabel',{'0.01','0.1','0','10','100'},'YMinorTick','on','YScale','log',...
    'YTick',[0.01 0.1 1 10 100],'YTickLabel',{'0.01','0.1','1','10','100'});
view([-65 35])


%% ++++++++++++++++++++++++++++++++++++++++++++++++++++++++++++++++++
%                                 SDH Dynamics Simulation
%  ++++++++++++++++++++++++++++++++++++++++++++++++++++++++++++++++++

G_SMP_O2 = p.G_SMP_O2;

fun = @main_complexII; 

fittingfunctions = {@G2018_F2C};

model = getoutput(fun,p,fittingfunctions);

figure
axes('FontName','Times New Roman','FontSize',55)
hold on
box on
plot(model.Y{1}(:,1),model.Y{1}(:,2),'linewidth',8)
plot(data.Y{7}(:,1),data.Y{7}(:,2),'o','markersize',30,'linewidth',6)
xlabel('[O_2] (\muM)')
ylabel('J_{H_2O_2} (nmol/mg/min)')
set(gcf,'color','none'),set(gca,'color','none')
set(gcf, 'Position', [1 1 1920 1080]); % Maximize figure.


%% Simulate ODE model

p.T = 273.15 + 25; % 25 C

p.Etot=21e-9; % M (0.2 mg/ml, 100 pmol/mg = 20 nM SDH)

% define integration absolute tolerances for matrix and cytosol SVs
AbsTol_vec=1e-6.*ones(1,numel(fieldnames(p.in)));

% set global integrator options
options = odeset('RelTol',1e-3,'AbsTol',AbsTol_vec,'Vectorized','on','BDF','on');

% set ICs
ic=zeros(numel(fieldnames(p.in)),1);
ic(p.in.O2)=235e-6;
ic(p.in.H)=10^-7.5;
ic(p.in.Q) = 20e-3;
ic(p.in.QH2) = 0;
ic(p.in.Succinate) =5000e-6;
ic(p.in.Fumarate) = 0e-6;

[T,X] = ode15s(@sdh_ode,[0 12*60],ic,options,p);

[J,JSO,JH2O2,JQH2,~,d] = main_complexII(X',p);

figure
set(gcf,'defaultAxesColorOrder',[[0 0 0]; [0 0 0]]);
axes('FontName','Times New Roman','FontSize',55)
hold on
box on
yyaxis left
plot(G_SMP_O2(1:3:end,1),G_SMP_O2(1:3:end,2),'o','markersize',30,'linewidth',6,'color',[0 0 0])
plot(T/60,X(:,p.in.O2)*1e6,'linewidth',8,'color',[0 0.447 0.741])
ylabel('[O_2] (\muM)','color',[0 0.447 0.741])
yyaxis right
plot(T/60,X(:,p.in.QH2)*1e3,'linewidth',8,'color',[0.85 0.325 0.098])
ylabel('[QH_2] (mM)','color',[0.85 0.325 0.098])
xlabel('Time (min)')
set(gcf,'color','none'),set(gca,'color','none')
set(gcf, 'Position', [1 1 1920 1080]); % Maximize figure.

figure
set(gcf,'defaultAxesColorOrder',[[0 0 0]; [0 0 0]]);
axes('FontName','Times New Roman','FontSize',55)
hold on
box on
yyaxis left
plot(T/60,(JH2O2+JSO/2)*1e12.*60/0.2/1e3,'linewidth',8,'color',[0 0.447 0.741])
ylabel('J_{H_2O_2} (pmol/mg/min)','color',[0 0.447 0.741])
yyaxis right
plot(T/60,X(:,p.in.H2O2)*1e9,'linewidth',8,'color',[0.85 0.325 0.098])
ylabel('[H_2O_2] (nM)','color',[0.85 0.325 0.098])
xlabel('Time (min)')
set(gcf,'color','none'),set(gca,'color','none')
set(gcf, 'Position', [1 1 1920 1080]); % Maximize figure.

figure
axes('FontName','Times New Roman','FontSize',55)
hold on
box on
plot(T/60,d.E0,T/60,d.E1,T/60,d.E2,T/60,d.E3,T/60,d.E4,'linewidth',8)
ylabel('Electronic State Fraction')
xlabel('Time (min)')
set(gcf,'color','none'),set(gca,'color','none')
set(gcf, 'Position', [1 1 1920 1080]); % Maximize figure.
               
               

%% Simulate Slater et al. 1968 Fig. 1A
% The SDH is isolated from the pig heart. The kinitics measurements are carried 
% at 25C, with 0.1 M phosphate buffer (pH 7.8), 1mM EGTA and 1mg/ml serum albumin.
% partially activated SDH is used in the experiments.
%In this experiment the rate of the oxidation of succinate is seen at
%different acceptor (PMS) concentrations. 90uM of malonate is present
%the Enzyme concentration is 0.0027uM

function model = S1968_F1A1(fun,p,model)
p = updateparams(p,'S1968');
% define model output according to data
labels = {'Slater_Fig1Aset1a.txt' 'Slater_Fig1Aset1b.txt' 'Slater_Fig1Aset1c.txt' 'Slater_Fig1Aset1d.txt' 'Slater_Fig1Aset1e.txt'};

% simulate independent variables 
Succinate = linspace(0,2.5,101).*1e-3;  % M
PMS = [0.067 0.1 0.167 0.30 2.0].*1e-3; %M

% create arrays for SV and fluxes
x = zeros(numel(fieldnames(p.in)),numel(Succinate));
J = zeros(numel(PMS),numel(Succinate));

% specify experimental conditions
x(p.in.O2,:)        = 200e-6;           %M  
x(p.in.Q,:)         = 1e-99;            %M
x(p.in.Succinate,:) = Succinate;        %M
x(p.in.Malonate,:)  = 90e-6;            %M
x(p.in.H,:)         = 10^(-7.8);        %pH = 7.8
p.T                 = 273.15 + 25;      % 25 C
p.atpenin           = 0;
p.Etot              = 27e-9;            %M (given)

% simulate experiments 
for i = 1:numel(PMS)
    x(p.in.GEA,:) = PMS(i);             %M
    [~,~,~,~,J(i,:),details,] = fun(x,p);
end

% format model output
for i = 1:numel(PMS)
    model.label{numel(model.label)+1} = labels{i};
    model.Y{numel(model.Y)+1}         = [Succinate.*1e3; J(i,:)*1e3*60*21]'; % [mM dA600/min] %detla absorbance at 600 index is 21 mM^-1 cm^-1 (King JBC 1963)
    model.g{numel(model.g)+1}         = details;
end


%% Simulate Vinogradov et al 1976 Fig 2
%SDH is isolated from bovine heart.
%The reaction mixture contain 10mM Succinate, 10mM Tris_sulphate buffer,
%pH 7.8, 0.1mM EGTA and experiments are calculate at 22C. No activation of
%enzyme was observed.
%In this experiment the rate of the oxidation of succinate is seen at
%different acceptor (WB) concentrations. 100uM of malonate is present.The 
%enzyme concentration is 5.5 ug per ml of assay. (may be 2-3 ml of assay
%they used)

function model = V1979_F21(fun,p,model)
p = updateparams(p,'V1979');

% define model output according to data
labels = {'Vinog_Fig2Aset1a.txt' 'Vinog_Fig2Aset1b.txt' 'Vinog_Fig2Aset1c.txt' 'Vinog_Fig2Aset1d.txt'};

% simulate independent variables 
Succinate = linspace(0,8,101).*1e-3;    % mM
WB = [0.05 0.066 0.1 0.33]*1e-3;        % M  

% create arrays for SV and fluxes 
x = zeros(numel(fieldnames(p.in)),numel(Succinate));
J = zeros(numel(WB),numel(Succinate));

% experimental details
x(p.in.O2,:)        = 200e-6;           % M  
x(p.in.Q,:)         = 1e-99;            % M
x(p.in.Succinate,:) = Succinate;        % M
x(p.in.Malonate,:)  = 100e-6;           % M      
x(p.in.H,:)         = 10^(-7.8);        % pH = 7.8
p.T                 = 273.15 + 22;      % 22 C
p.atpenin           = 0;
p.Etot              = 44.7e-9*p.xVinog; % M 5.5 ug enzyme/ml (using MW of 123 kD to convert)

for i = 1:numel(WB)
    x(p.in.GEA,:) = WB(i);              %M
    [~,~,~,~,J(i,:),details] = fun(x,p);
    model.label{numel(model.label)+1} = labels{i};
    model.Y{numel(model.Y)+1}         = [Succinate.*1e3; J(i,:)*1e6*60/5.5]'; % umol/min/mg protein (M/s x 60 s/min x 1e6 uM/M x 1/5.5 L/mg)
    model.g{numel(model.g)+1}         = details;
end

%% Simulate Vinogradov et al 1976 Fig 7
%SDH is isolated from bovine heart.
%The experiments are conducted at 22C, 20mM Succinate is present.
%No activation of enzyme was observed.
%The enzyme concentration is 0.8 - 2 ug per ml of assay.
%In these experiments the effect of varing pH on the rate of succinate
%oxidation is seen.(may be 2-3 ml of assay
%they used)

function model = V1979_F71(fun,p,model)
p = updateparams(p,'V1979');

% define model output according to data
labels = {'Vinog_Fig7Aset1e.txt' 'Vinog_Fig7Aset1d.txt' 'Vinog_Fig7Aset1c.txt' 'Vinog_Fig7Aset1b.txt' 'Vinog_Fig7Aset1a.txt'};

% simulate independent variables
pH = linspace(6,10,101);
WB = [50 66 100 200 1e3]*1e-6;              % M

% create arrays for SV and fluxes
x = zeros(numel(fieldnames(p.in)),numel(pH));
J = zeros(numel(WB),numel(pH));

% experimental details
x(p.in.O2,:)            = 200e-6;           % M  
x(p.in.Succinate,:)     = 20e-3;            % M
x(p.in.H,:)             = 10.^-pH;          % pH 
x(p.in.Q,:)             = 1e-99;            % M
p.T                     = 273.15 + 22;      % 22 C
p.atpenin               = 0;
p.Etot                  = 8.45e-9*p.xVinog; % M 0.8-2 ug enzyme/ml (using MW of 123 kD to convert)

for i = 1:numel(WB)
    x(p.in.GEA,:) = WB(i);                  % M 
    [~,~,~,~,J(i,:),details] = fun(x,p);
    model.label{numel(model.label)+1} = labels{i};
    model.Y{numel(model.Y)+1}         = [pH; J(i,:)*1e6*60/1.4]'; % umol/min/mg protein
    model.g{numel(model.g)+1}         = details;
end


%% Simulate Hirst 2013, Figure 3C
% The submitochondrial particles (SMP)are obtaied from the
% bovine heart mitochondria. All assays were carried out in
% 10 mM Tris�SO4 (pH 7.4 at 32 C),250 mM sucrose, 2 mM MgSO4, and 1 mM K2SO4 at 32 C
% In this experiment the rate of succinate oxidation is measured at
% different succinate concentrations.

function model = H2013_F3A(fun,p,model)

% simulate independent variables
Succinate = linspace(0,20,201)*1e-3; % M

% create array for SV
x = zeros(numel(fieldnames(p.in)),numel(Succinate));

% specify experimental conditions 
p.T                 = 273.15 + 37;       % 32 C
p.atpenin           = 0;
p.Etot              = 3e-9*p.xHirst;     % M (going with 100 pmol/mg, 30 ug/mL x 1000 mL/L x 100 pmol/mg x 1e-12 mol/pmol)
x(p.in.O2,:)        = 200e-6;            % M
x(p.in.Succinate,:) = Succinate;         % M
x(p.in.QH2,:)       = 20e-3*(1./(1+ p.Q./Succinate)).*(1./(1+ p.atpenin./p.Q_atpenin));        % M
x(p.in.Q,:)         =  20e-3 - x(p.in.QH2,:);
x(p.in.H,:)         = 10^(-7.2);         % pH = 7.4

% simulate experiments 
[~,~,~,J,~,details] = fun(x,p);

% format model output
model.label{numel(model.label)+1} = 'Hirst_Fig3Aset1a.txt';
model.Y{numel(model.Y)+1}         = [Succinate.*1e3; J*1e6.*60/30]'; % umol.min^-1.mg.protein, assuming 30 ug/ml
model.g{numel(model.g)+1}         = details;


%% Simulate Hirst 2013, Figure 3C
% The submitochondrial particles (SMP)are obtaied from the
% bovine heart mitochondria. All assays were carried out in
% 10 mM Tris�SO4 (pH 7.4 at 32 C),250 mM sucrose, 2 mM MgSO4, and 1 mM K2SO4 at 32 C
% In this experiment the rate of succinate oxidation is measured at
% different A5 concentrations.

function model = H2013_F3B(fun,p,model)

% simulate independent variables 
atpenin = linspace(0,100,501).*1e-9; %M

% create array for SV
x = zeros(numel(fieldnames(p.in)),numel(atpenin));

Succinate = 5e-3; % M

% specify experimental conditions 
p.T                 = 273.15 + 32;      % 32 C
p.Etot              = 3e-9*p.xHirst;    % M (going with 100 pmol/mg, 30 ug/mL x 1000 mL/L x 100 pmol/mg x 1e-12 mol/pmol)
p.atpenin           = atpenin;
x(p.in.O2,:)        = 200e-6;           % M
x(p.in.Succinate,:) = Succinate;        % M
x(p.in.QH2,:)       = 20e-3*(1./(1+ p.Q./Succinate)).*(1./(1+ p.atpenin./p.Q_atpenin));        % M
x(p.in.Q,:)         =  20e-3 - x(p.in.QH2,:);
x(p.in.H,:)         = 10^(-7.4);        % pH = 7.4

% simulate experiments 
[~,~,~,J,~,details] = fun(x,p);

% format model output
model.label{numel(model.label)+1} = 'Hirst_Fig3Bset1a.txt';
model.Y{numel(model.Y)+1}         = [atpenin.*1e9; (J/J(1))*100]'; % percent of initial
model.g{numel(model.g)+1}         = details;


%% Simulate Hirst 2013, Figure 3C
% The submitochondrial particles (SMP)are obtaied from the
% bovine heart mitochondria. All assays were carried out in
% 10 mM Tris�SO4 (pH 7.4 at 32 C),250 mM sucrose, 2 mM MgSO4, and 1 mM K2SO4 at 32 C
% In this experiment the rate of succinate oxidation is measured at
% different Malonate concentrations.

function model = H2013_F3C(fun,p,model)

% simulate independent variables 
Malonate = linspace(0,5,101).*1e-3; %M

Succinate = 5e-3; % M

% create array for SVs
x = zeros(numel(fieldnames(p.in)),numel(Malonate));

p.T                 = 273.15 + 32;   % 32 C
p.Etot              = 3e-9*p.xHirst; % M (going with 100 pmol/mg, 30 ug/mL x 1000 mL/L x 100 pmol/mg x 1e-12 mol/pmol)
p.atpenin           = 0;

% specify experimental conditions 
x(p.in.O2,:)        = 200e-6;        % M
x(p.in.Succinate,:) = Succinate;     % M
x(p.in.QH2,:)       = 20e-3*(1./(1+ p.Q./Succinate)).*(1./(1+ p.atpenin./p.Q_atpenin)).*(1./(1+ Malonate./p.Q_MALO));        % M
x(p.in.Q,:)         = 20e-3 - x(p.in.QH2,:);
x(p.in.H,:)         = 10^(-7.4);     % pH = 7.4
x(p.in.Malonate,:)  = Malonate;      % M 

% simulate experiments 
[~,~,~,J,~,details] = fun(x,p);

% format model output
model.label{numel(model.label)+1} = 'Hirst_Fig3Cset1a.txt';
model.Y{numel(model.Y)+1}         = [Malonate.*1e3; J/J(1)*100]'; % percent of initial
model.g{numel(model.g)+1}         = details;

%% Simulate Siebels and Drose, 2013, Figure 1
% The submitochondrial particles (SMP)are obtaied from the
% bovine heart mitochondria (0.12 mg protien /ml).
% SMP 24mg protein per cavity; total assay volume 200ul were dissolved in a
% buffer contating 75 mM sodium phosphate, 1mM EGTA, and 1mM Mgcl2 (pH
% 7.2 & 8.0). The experiments may conducted at 37c. In these experiments at the 
% different A5 concentrations, the ROS production and succinate oxidation
% activity is seen. Stigmatellin = 1 uM

function model = S2013_F1A1(fun,p,model)
p = updateparams(p,'s2013');

p.T = 273.15 + 37;                   % 37 C
p.Etot = 12e-9*p.xSiebe;             % M (100 pmol/mg x 0.12 mg/ml x 1e-12 mol/pmol x 1000 mL/L)

% simulate independent variables 
Succinate = linspace(0,10,201)*1e-3; % M

% create array for SV
x = zeros(numel(fieldnames(p.in)),numel(Succinate));

% specify experimental conditions 
x(p.in.O2,:)        = 200e-6;        % M
x(p.in.Succinate,:) = Succinate;     % M 
x(p.in.H,:)         = 10^(-7.2);     % M

%% Stigmatellin
p.atpenin           = 0;        % M
Malonate            = 0;        % M
x(p.in.Malonate,:)  = Malonate; % M

x(p.in.QH2,:)       = 20e-3*(1./(1+ p.Q_stig./Succinate)).*(1./(1+ Malonate./p.Q_MALO));        %M
x(p.in.Q,:)         =  20e-3 - x(p.in.QH2,:);                                                   %M 

% simulate experiment
[~, JSO,JH2O2,~,~,details]= fun(x,p);
JH2O2tot = JH2O2 + JSO/2;

% formate data into structure 
model.label{numel(model.label)+1} = 'Siebels13_fig1Aset1a.txt';
model.Y{numel(model.Y)+1}         = [Succinate*1e3; JH2O2tot*1e12.*60/0.12/1e3]'; % pmol.min^-1.mg.protein
model.g{numel(model.g)+1}         = details;

%% Stigmatellin + Malonate
p.atpenin           = 0;        % M
Malonate            = 1.5e-3;   % M
x(p.in.Malonate,:)  = Malonate; % M

x(p.in.QH2,:)   = 20e-3*(1./(1+ p.Q_stig./Succinate)).*(1./(1+ Malonate./p.Q_MALO));        %M
x(p.in.Q,:)     =  20e-3 - x(p.in.QH2,:);                                                   %M 

% simulate experiment
[~, JSO,JH2O2,~,~,details]= fun(x,p);
JH2O2tot = JH2O2 + JSO/2;

% formate data into structure 
model.label{numel(model.label)+1} = 'Siebels13_fig1Aset1b.txt';
model.Y{numel(model.Y)+1}         = [Succinate*1e3; JH2O2tot*1e12.*60/0.12/1e3]'; % pmol.min^-1.mg.protein
model.g{numel(model.g)+1}         = details;


%% Atpenin
p.atpenin           = 50e-9;    % M
Malonate            = 0;        % M
x(p.in.Malonate,:)  = Malonate; % M

x(p.in.QH2,:) = 20e-3*(1./(1+ p.Q./Succinate)).*(1./(1+ p.atpenin./p.Q_atpenin)).*(1./(1+ Malonate./p.Q_MALO));        %M
x(p.in.Q,:)   = 20e-3 - x(p.in.QH2,:);                                                                                 %M 

% simulate experiment
[~, JSO,JH2O2,~,~,details]= fun(x,p);
JH2O2tot = JH2O2 + JSO/2;

% formate data into structure 
model.label{numel(model.label)+1} = 'Siebels13_fig1Aset1c.txt';
model.Y{numel(model.Y)+1}         = [Succinate*1e3; JH2O2tot*1e12.*60/0.12/1e3]'; % pmol.min^-1.mg.protein
model.g{numel(model.g)+1}         = details;

%% Atpenin + Malonate
p.atpenin           = 50e-9;        % M
Malonate            = 1.5e-3;       % M
x(p.in.Malonate,:)  = Malonate;     % M

x(p.in.QH2,:)       = 20e-3*(1./(1+ p.Q./Succinate)).*(1./(1+ p.atpenin./p.Q_atpenin)).*(1./(1+ Malonate./p.Q_MALO));        %M
x(p.in.Q,:)         = 20e-3 - x(p.in.QH2,:);                                                                                 %M 

% simulate experiment
[~, JSO,JH2O2,~,~,details]= fun(x,p);
JH2O2tot = JH2O2 + JSO/2;

% formate data into structure 
model.label{numel(model.label)+1} = 'Siebels13_fig1Aset1d.txt';
model.Y{numel(model.Y)+1}         = [Succinate*1e3; JH2O2tot*1e12.*60/0.12/1e3]'; % pmol.min^-1.mg.protein
model.g{numel(model.g)+1}         = details;

%% Malonate
p.atpenin           = 0;        % M
Malonate            = 1.5e-3;   % M
x(p.in.Malonate,:)  = Malonate; % M

x(p.in.QH2,:)       = 20e-3*(1./(1+ p.Q./Succinate)).*(1./(1+ p.atpenin./p.Q_atpenin)).*(1./(1+ Malonate./p.Q_MALO));        %M
x(p.in.Q,:)         =  20e-3 - x(p.in.QH2,:);                                                                                %M 

% simulate experiment
[~, JSO,JH2O2,~,~,details]= fun(x,p);
JH2O2tot = JH2O2 + JSO/2;

% formate data into structure 
model.label{numel(model.label)+1} = 'Siebels13_fig1Aset1e.txt';
model.Y{numel(model.Y)+1} = [Succinate*1e3; JH2O2tot*1e12.*60/0.12/1e3]'; % pmol.min^-1.mg.protein
model.g{numel(model.g)+1}  = details;


%% Simulate Siebels and Drose, 2013, Figure 1b
% The submitochondrial particles (SMP)are obtaied from the
% bovine heart mitochondria (0.12 mg protien /ml). SDH conc 1.2-12 nM.
% SMP 24mg protein per cavity; total assay volume 200ul were dissolved in a
% buffer contating 75 mM sodium phosphate, 1mM EGTA, and 1mM Mgcl (pH
% 7.2). The experiments may conducted at 37c. In this experiment at
% different succinate concentrations ROS production is seen. The atpenin 
% concentration is (50nM).

function model = S2013_F1B(fun,p,model)
p = updateparams(p,'s2013');

p.T         = 273.15 + 37;      % 37 C
p.Etot      = 12e-9*p.xSiebe;   % M (100 pmol/mg x 0.12 mg/ml x 1e-12 mol/pmol * 1000 mL/L)
p.atpenin   = 50e-9;            % M 

% simulate independent variables 
Succinate = [linspace(0,1,501) linspace(1.1,8,101)].*1e-3; %M

% create array for SV
x = zeros(numel(fieldnames(p.in)),numel(Succinate));

% specify experimental conditions 
x(p.in.O2,:)        = 200e-6;                                                                  % M
x(p.in.Succinate,:) = Succinate;                                                               % M
x(p.in.QH2,:)       = 20e-3*(1./(1+ p.Q./Succinate)).*(1./(1+ p.atpenin./p.Q_atpenin));        % M
x(p.in.Q,:)         = 20e-3 - x(p.in.QH2,:);                                                   % M
x(p.in.H,:)         = 10^(-7.2);                                                               % ROS data could be at 7.2

% simulate experiment 
[~,JSO,JH2O2,~,~,details] = fun(x,p);
JH2O2tot = JH2O2 + JSO/2;

% format data for packing into structure 
model.label{numel(model.label)+1} = 'Siebels13_fig1Bset1a.txt';
model.Y{numel(model.Y)+1}         = [Succinate.*1e3; JH2O2tot.*1e12.*60/.12/1e3]'; % pmol.min^-1.mg.protein assuming .12 mg/ml SMP (Succinate is in mM)
model.g{numel(model.g)+1}         = details;

%% Simulate Siebels and Drose, 2013, Figure 1c
% The submitochondrial particles (SMP)are obtaied from the
% bovine heart mitochondria (0.12 mg protien /ml).
% SMP 24mg protein per cavity; total assay volume 200ul were dissolved in a
% buffer contating 75 mM sodium phosphate, 1mM EGTA, and 1mM Mgcl2 (pH
% 7.2). The experiments may conducted at 37c. In these experiments at the 
% different A5 concentrations, the ROS production and succinate oxidation
% activity is seen. succinate is 100uM. 

function model = S2013_F1C(fun,p,model)
p = updateparams(p,'s2013');

p.T         = 273.15 + 37;                  % 37 C
p.Etot      = 12e-9*p.xSiebe;               % M (100 pmol/mg x 0.12 mg/ml x 1e-12 mol/pmol * 1000 mL/L)
p.atpenin   = linspace(0,250,501).*1e-9;    % M

% simulate independent variables 
Succinate   = 100e-6;                       % M

% create array for SV
x = zeros(numel(fieldnames(p.in)),numel(p.atpenin));

% specify experimental conditions 
x(p.in.O2,:)        = 200e-6;                                                                  % M
x(p.in.Succinate,:) = Succinate;                                                               % M
x(p.in.QH2,:)       = 20e-3*(1./(1+ p.Q./Succinate)).*(1./(1+ p.atpenin./p.Q_atpenin));        % M
x(p.in.Q,:)         = 20e-3 - x(p.in.QH2,:);
x(p.in.H,:)         = 10^(-7.2);                                                               % ROS data could be at 7.2

% simulate experiment
[~,JSO,JH2O2,JQH2,~,details] = fun(x,p);
JH2O2tot = JH2O2 + JSO/2; 

% format model output
model.label{numel(model.label)+1} = 'Siebels13_fig1Cset1a.txt';
model.Y{numel(model.Y)+1}         = [p.atpenin.*1e9; JH2O2tot*1e12.*60/0.12/1e3]'; % [nM pmol.min^-1.mg.protein]
model.g{numel(model.g)+1}         = details;

model.label{numel(model.label)+1} = 'Siebels13_fig1Cset2a.txt';
model.Y{numel(model.Y)+1}         = [p.atpenin.*1e9; JQH2/JQH2(1)*100]'; % [nM %]
model.g{numel(model.g)+1}         = details;

model.label{numel(model.label)+1} = 'Siebels13_fig1Cset3a.txt';
model.Y{numel(model.Y)+1}         = [[1 2];[1 JQH2(1)*1e9.*60/0.12/1e3/2]]'; % max rate is 78.9 nmol O2/mg/min
model.g{numel(model.g)+1}         = details;


%% Simulate Siebels and Drose, 2013, Figure 1
% The submitochondrial particles (SMP)are obtaied from the
% bovine heart mitochondria (0.12 mg protien /ml).
% SMP 24mg protein per cavity; total assay volume 200ul were dissolved in a
% buffer contating 75 mM sodium phosphate, 1mM EGTA, and 1mM Mgcl2 (pH
% 7.2 & 8.0). The experiments may conducted at 37c. In these experiments at the 
% different A5 concentrations, the ROS production and succinate oxidation
% activity is seen.

function model = S2013_F1D(fun,p,model)
p = updateparams(p,'s2013');

p.T             = 273.15 + 37;             % 37 C
p.Etot          = 12e-9*p.xSiebe;          % M (100 pmol/mg x 0.12 mg/ml x 1e-12 mol/pmol x 1000 mL/L)
p.atpenin       = 250e-9;                  % M

% simulate independent variable
Succinate       = linspace(0,10,101)*1e-3; % M

% create array for SV
x = zeros(numel(fieldnames(p.in)),numel(Succinate));

% experimental details
x(p.in.O2,:)        = 200e-6;              % M
x(p.in.Succinate,:) = Succinate;           % M 
x(p.in.H,:)         = 10^(-7.2);           % ROS data could be at 7.2


%% Atpenin
p.atpenin           = 250e-9;    % M
Malonate            = 0;               % M
x(p.in.Malonate,:)  = Malonate;        % M

x(p.in.QH2,:)       = 20e-3*(1./(1+ p.Q./Succinate)).*(1./(1+ p.atpenin./p.Q_atpenin)).*(1./(1+ Malonate./p.Q_MALO));  % M
x(p.in.Q,:)         = 20e-3 - x(p.in.QH2,:);                                                                           % M 

% simulate experiment
[~, JSO,JH2O2,~,~,details]= fun(x,p);
JH2O2tot = JH2O2 + JSO/2;

% formate data into structure 
model.label{numel(model.label)+1} = 'Siebels13_fig1Dset1a.txt';
model.Y{numel(model.Y)+1}         = [Succinate*1e3; JH2O2tot*1e12.*60/0.12/1e3]'; % pmol.min^-1.mg.protein
model.g{numel(model.g)+1}         = details;

%% Atpenin + Malonate
p.atpenin           = 250e-9;     % M
Malonate            = 1.5e-3;           % M
x(p.in.Malonate,:)  = Malonate;         % M

x(p.in.QH2,:) = 20e-3*(1./(1+ p.Q./Succinate)).*(1./(1+ p.atpenin./p.Q_atpenin)).*(1./(1+ Malonate./p.Q_MALO));     %M
x(p.in.Q,:) =  20e-3 - x(p.in.QH2,:);                                                                               %M 

% simulate experiment
[~, JSO,JH2O2,~,~,details]= fun(x,p);
JH2O2tot = JH2O2 + JSO/2;

% formate data into structure 
model.label{numel(model.label)+1} = 'Siebels13_fig1Dset1b.txt';
model.Y{numel(model.Y)+1}         = [Succinate*1e3; JH2O2tot*1e12.*60/0.12/1e3]'; % pmol.min^-1.mg.protein
model.g{numel(model.g)+1}         = details;


%% Simulate Siebels and Drose, 2013, Figure 2
% The effect of pH on superoxide (SO) and total (SO and H2O2) was investigated by measuring SO and total ROS
% at 2 different pH (7.2 and 8.0). 
% Bovine heart mitochondria = 0.12 mg protein /ml
% Succinate concentration = 100 uM
% Stigmatellin = 1 uM (no atpenin A5)

function model = S2013_F2(fun,p,model)
p = updateparams(p,'s2013');

p.T         = 273.15 + 37;      % 37 C
p.Etot      = 12e-9*p.xSiebe;   % M (100 pmol/mg x 0.12 mg/ml x 1e-12 mol/pmol / 1000 mL/L)

Succinate   = 100e-6;           % M
pH          = linspace(7,8.5,101);

% create array for SV
x = zeros(numel(fieldnames(p.in)),numel(pH));

% specify experimental conditions 
x(p.in.O2,:)        = 200e-6;        % M
x(p.in.Succinate,:) = Succinate;     % M
x(p.in.H,:)         = 10.^(-pH);

%% Stigmatellin Experiment
p.atpenin           = 0; % M
Malonate            = 0; % M
x(p.in.Malonate,:)  = Malonate; % M

% simulate experiments
x(p.in.QH2,:)   = 20e-3*(1./(1+ p.Q_stig./Succinate)).*(1./(1+ Malonate./p.Q_MALO));  % M
x(p.in.Q,:)     = 20e-3 - x(p.in.QH2,:);                                              % M  
[~, JSO, JH2O2,~,~,details] = fun(x,p);
JSO             = JSO*(1-p.xSOD);
JH2O2           = JH2O2+p.xSOD*JSO/2;
JH2O2eq         = 2*JH2O2 + JSO;                                                      % ROS per O2

% format model output
% supeoxide
model.label{numel(model.label)+1} = 'Siebels13_fig2set1a.txt';
model.Y{numel(model.Y)+1}         = [pH; [JSO*1e12.*60/0.12/1e3]]';      % pH pmol.min^-1.mg.protein
model.g{numel(model.g)+1}         = details;
% hydrogen peroxide
model.label{numel(model.label)+1} = 'Siebels13_fig2set2a.txt';
model.Y{numel(model.Y)+1}         = [pH; [JH2O2eq*1e12.*60/0.12/1e3]]';  % pH pmol.min^-1.mg.protein
model.g{numel(model.g)+1}         = details;

%% Stigmatellin and Malonate
Malonate            = 1.5e-3;   % M
x(p.in.Malonate,:)  = Malonate; % M

% simulate experiments under pH = 7.2
x(p.in.QH2,:)       = 20e-3*(1./(1+ p.Q_stig./Succinate)).*(1./(1+ Malonate./p.Q_MALO));  % M
x(p.in.Q,:)         =  20e-3 - x(p.in.QH2,:);                                             % M  
[~, JSO, JH2O2,~,~,details] = fun(x,p);
JSO                 = JSO*(1-p.xSOD);
JH2O2               = JH2O2+p.xSOD*JSO/2;
JH2O2eq             = 2*JH2O2 + JSO;                                                      % ROS per O2

% format model output
% supeoxide
model.label{numel(model.label)+1} = 'Siebels13_fig2set1b.txt';
model.Y{numel(model.Y)+1}         = [pH; [JSO*1e12.*60/0.12/1e3]]';      % pH pmol.min^-1.mg.protein
model.g{numel(model.g)+1}         = details;
% hydrogen peroxide
model.label{numel(model.label)+1} = 'Siebels13_fig2set2b.txt';
model.Y{numel(model.Y)+1}         = [pH; [JH2O2eq*1e12.*60/0.12/1e3]]';  % pH pmol.min^-1.mg.protein
model.g{numel(model.g)+1}         = details;

%% Atpenin Experiment
p.atpenin           = 250e-9;     % M
Malonate            = 0;                % M
x(p.in.Malonate,:)  = Malonate;         % M

x(p.in.QH2,:)       = 20e-3*(1./(1+ p.Q./Succinate)).*(1./(1+ p.atpenin./p.Q_atpenin)).*(1./(1+ Malonate./p.Q_MALO));  %M
x(p.in.Q,:)         = 20e-3 - x(p.in.QH2,:);
[~, JSO, JH2O2,~,~,details] = fun(x,p);
JSO                 = JSO*(1-p.xSOD);
JH2O2               = JH2O2+p.xSOD*JSO/2;
JH2O2eq             = 2*JH2O2 + JSO;    % ROS per O2

% format model output
% supeoxide
model.label{numel(model.label)+1} = 'Siebels13_fig2set1c.txt';
model.Y{numel(model.Y)+1}         = [pH; [JSO*1e12.*60/0.12/1e3]]';  % pH pmol.min^-1.mg.protein
model.g{numel(model.g)+1}         = details;
% hydrogen peroxide
model.label{numel(model.label)+1} = 'Siebels13_fig2set2c.txt';
model.Y{numel(model.Y)+1}         = [pH; [JH2O2eq*1e12.*60/0.12/1e3]]';  % pH pmol.min^-1.mg.protein
model.g{numel(model.g)+1}         = details;

%% Atpenin and Malonate
p.atpenin           = 250e-9;     % M
Malonate            = 1.5e-3;           % M
x(p.in.Malonate,:)  = Malonate;         % M

% simulate experiments under pH = 7.2
x(p.in.QH2,:)   = 20e-3*(1./(1+ p.Q./Succinate)).*(1./(1+ p.atpenin./p.Q_atpenin)).*(1./(1+ Malonate./p.Q_MALO));  % M
x(p.in.Q,:)     = 20e-3 - x(p.in.QH2,:);                                                                           % M  
[~, JSO, JH2O2,~,~,details] = fun(x,p);
JSO             = JSO*(1-p.xSOD);
JH2O2           = JH2O2+p.xSOD*JSO/2;
JH2O2eq         = 2*JH2O2 + JSO;        % ROS per O2

% format model output
% supeoxide
model.label{numel(model.label)+1} = 'Siebels13_fig2set1d.txt';
model.Y{numel(model.Y)+1}         = [pH; [JSO*1e12.*60/0.12/1e3]]';      % pH pmol.min^-1.mg.protein
model.g{numel(model.g)+1}         = details;
% hydrogen peroxide
model.label{numel(model.label)+1} = 'Siebels13_fig2set2d.txt';
model.Y{numel(model.Y)+1}         = [pH; [JH2O2eq*1e12.*60/0.12/1e3]]';  % pH pmol.min^-1.mg.protein
model.g{numel(model.g)+1}         = details;

%% Malonate
p.atpenin           = 0;                    % M
x(p.in.Malonate,:)  = 1.5e-3;               % M

% simulate experiments under pH = 7.2
x(p.in.QH2,:)       = 20e-3*(1./(1+ p.Q./Succinate)).*(1./(1+ p.atpenin./p.Q_atpenin)).*(1./(1+ Malonate./p.Q_MALO));  % M
x(p.in.Q,:)         =  20e-3 - x(p.in.QH2,:);                                                                          % M  
[~, JSO, JH2O2,~,~,details] = fun(x,p);
JSO                 = JSO*(1-p.xSOD);
JH2O2               = JH2O2+p.xSOD*JSO/2;
JH2O2eq             = 2*JH2O2 + JSO;        % ROS per O2

% format model output
% supeoxide
model.label{numel(model.label)+1} = 'Siebels13_fig2set1e.txt';
model.Y{numel(model.Y)+1}         = [pH; [JSO*1e12.*60/0.12/1e3]]';  % pH pmol.min^-1.mg.protein
model.g{numel(model.g)+1}         = details;
% hydrogen peroxide
model.label{numel(model.label)+1} = 'Siebels13_fig2set2e.txt';
model.Y{numel(model.Y)+1}         = [pH; [JH2O2eq*1e12.*60/0.12/1e3]]';  % pH pmol.min^-1.mg.protein
model.g{numel(model.g)+1}         = details;

%% Simulate Siebels and Drose, 2013, Figure 3A
% Effect of fumarate titration (fumarate/ succinate ratio) on H2O2 production in the presenve of atpenin A5 was investigated.
% Bovine heart mitochondria = 0.12 mg protein /ml
% atpenin A5 = 50 nM 

function model = S2013_F3A(fun,p,model)
p = updateparams(p,'s2013');

p.T       = 273.15 + 37;     % 37 C
p.Etot    = 12e-9*p.xSiebe;  % M (100 pmol/mg x 0.12 mg/ml x 1e-12 mol/pmol / 1000 mL/L)
p.atpenin = 50e-9;           % M

% create labels for output 
labels = {'Siebels13_fig3Aset1a.txt' 'Siebels13_fig3Aset1b.txt' 'Siebels13_fig3Aset1c.txt'};

% simulate independent variables
Fumarate  = linspace(0,6,101).*1e-3;   % M
Succinate = [0.1 0.5 1].*1e-3;         % M

% create arrays for SV and fluxes
x = zeros(numel(fieldnames(p.in)),numel(Fumarate));

% specify experimental conditions
x(p.in.O2,:)       = 200e-6;           % M 
x(p.in.H,:)        = 10^(-7.2);        % ROS data could be at 7.2
x(p.in.Fumarate,:) = Fumarate;         % M

JH2O2    = zeros(numel(Succinate),numel(Fumarate));
JSO      = zeros(numel(Succinate),numel(Fumarate));
JH2O2tot = zeros(numel(Succinate),numel(Fumarate));

% simulate experiment
for i=1:numel(Succinate)
    x(p.in.Succinate,:) = Succinate(i);                                                                                     % M
    x(p.in.QH2,:)       = 20e-3*(1./(1+ p.Q./Succinate(i))).*(1./(1+ p.atpenin./p.Q_atpenin)).*(1./(1+ Fumarate./p.Q_FUM)); % M
    x(p.in.Q,:)         =  20e-3 - x(p.in.QH2,1);                                                                           % M
    [~, JSO(i,:),JH2O2(i,:),~,~,details]= fun(x,p);
    JH2O2tot(i,:)       = JH2O2(i,:) + JSO(i,:)/2; 
    
    model.label{numel(model.label)+1} = labels{i};
    model.Y{numel(model.Y)+1}         = [Fumarate.*1e3; JH2O2tot(i,:)*1e12.*60/0.12/1e3]'; % pmol.min^-1.mg.protein (Fumarate is in mM)
    model.g{numel(model.g)+1}         = details;
end


%% Simulate Siebels and Drose, 2013, Figure 3A
% Effect of fumarate titration (fumarate/ succinate ratio) on H2O2 production in the presenve of stigmatellin was investigated.
% Bovine heart mitochondria = 0.12 mg protein /ml
% Stigmatellin = 1 uM  

function model = S2013_F3B(fun,p,model) 
p = updateparams(p,'s2013');
labels = {'Siebels13_fig3Bset2a.txt' 'Siebels13_fig3Bset2b.txt' 'Siebels13_fig3Bset2c.txt'};

p.T         = 273.15 + 37;              % 37 C
p.Etot      = 12e-9*p.xSiebe;           % M (100 pmol/mg x 0.12 mg/ml x 1e-12 mol/pmol / 1000 mL/L)
p.atpenin   = 0;                        % M

% simulate independent variables
Fumarate    = linspace(0,6,101).*1e-3;  % M
Succinate   = [0.1 0.5 1].*1e-3;        % M

% create arrays for SV and fluxes
x           = zeros(numel(fieldnames(p.in)),numel(Fumarate));
JH2O2       = zeros(numel(Succinate),numel(Fumarate));
JSO         = zeros(numel(Succinate),numel(Fumarate));
JH2O2tot    = zeros(numel(Succinate),numel(Fumarate));

% specify experimental conditions
x(p.in.O2,:)        = 200e-6;           % M
x(p.in.H,:)         = 10^(-7.2);        % ROS data could be at 7.2,
x(p.in.Fumarate,:)  = Fumarate;         % M

H                   = 10^-7.2;
Em.Q_QH2            = p.Em0.Q_QH2 + (p.R*p.T./p.F).*log(H);
Em.Fum_Suc          = p.Em0.Fum_Suc + (p.R*p.T./p.F).*log(H);
Keq                 = exp(2*p.F*(Em.Q_QH2-Em.Fum_Suc)/p.R/p.T);

% simulate experiments
for i=1:numel(Succinate)
    x(p.in.Succinate,:) = Succinate(i);                                         % M
    x(p.in.QH2,:)       = 20e-3*Succinate(i)*Keq./(Fumarate+Succinate(i)*Keq);
    x(p.in.Q,:)         = 20e-3 - x(p.in.QH2,:) + eps;
    x(p.in.QH2,1)       = 20e-3*(1./(1+ p.Q_stig./Succinate(i)));               % M
    x(p.in.Q,1)         =  20e-3 - x(p.in.QH2,1);                               % M
    [~, JSO(i,:),JH2O2(i,:),~,~,details]= fun(x,p);
    JH2O2tot(i,:)       = JH2O2(i,:) + JSO(i,:)/2;
    
    model.label{numel(model.label)+1} = labels{i};
    model.Y{numel(model.Y)+1}         = [Fumarate.*1e3; JH2O2tot(i,:)*1e12.*60/0.12/1e3]'; % pmol.min^-1.mg.protein (Fumarate is in mM)
    details.data                      = 'Siebels13_fig3B';
    model.g{numel(model.g)+1}         = details;
end


%% Simulate Siebels and Drose, 2013, Figure 4b
% The submitochondrial particles (SMP)are obtaied from the
% bovine heart mitochondria (0.12 mg protien /ml).
% SMP 24mg protein per cavity; total assay volume 200ul were dissolved in a
% buffer contating 75 mM sodium phosphate, 1mM EGTA, and 1mM Mgcl2 (pH
% 7.2). The experiments may conducted at 37c. In these experiments at the 
% different Malate concentrations, the ROS production and succinate oxidation
% activity is seen. Succinate is 100uM, and A5 = 250nM is used.

function model = S2013_F4B(fun,p,model)
p = updateparams(p,'s2013');

% experimental details
p.T         = 273.15 + 37;              % 37 C
p.Etot      = 12e-9*p.xSiebe;           % M (100 pmol/mg x 0.12 mg/ml x 1e-12 mol/pmol / 1000 mL/L)
p.atpenin   = 250e-9;                   % M

Malate      = linspace(0,10,101).*1e-3; % M
Succinate   = 100e-6;                   % M

x = zeros(numel(fieldnames(p.in)),numel(Malate));
x(p.in.O2,:)        = 200e-6;             % M
x(p.in.Succinate,:) = Succinate;          % M
x(p.in.H,:)         = 10^(-7.2);          % ROS data could be at 7.2
x(p.in.Malate,:)    = Malate;             % M
x(p.in.QH2,:)       = 20e-3*(1./(1+ p.Q./Succinate)).*(1./(1+ p.atpenin./p.Q_atpenin)).*(1./(1+ Malate./p.Q_MAL));  % M
x(p.in.Q,:)         = 20e-3 - x(p.in.QH2,:);

[~,JSO,JH2O2,~,~,details] = fun(x,p);
JH2O2tot = JH2O2 + JSO/2;

% format model output
model.label{numel(model.label)+1} = 'Siebels13_fig4Bset1a.txt';
model.Y{numel(model.Y)+1}         = [Malate*1e3; JH2O2tot*1e12.*60/0.12/1e3]'; % pmol.min^-1.mg.protein
model.g{numel(model.g)+1}         = details;

p.atpenin = 0;           %M
% Malate inhibits QH2 production, so QH2 should not be high when malate is high
x(p.in.QH2,:) = 20e-3*(1./(1+ p.Q./Succinate)).*(1./(1+ p.atpenin./p.Q_atpenin)).*(1./(1+ Malate./p.Q_MAL));        % M
x(p.in.Q,:)   =  20e-3 - x(p.in.QH2,:);
[~,~,~,JQH2,~,details] = fun(x,p);

% format model output
model.label{numel(model.label)+1} = 'Siebels13_fig4Bset2a.txt';
model.Y{numel(model.Y)+1}         = [Malate*1e3; JQH2/JQH2(1)*100]';
model.g{numel(model.g)+1}         = details;


%% Simulate Siebels and Drose, 2013, Figure 4c
% The submitochondrial particles (SMP)are obtaied from the
% bovine heart mitochondria (0.12 mg protien /ml).
% SMP 24mg protein per cavity; total assay volume 200ul were dissolved in a
% buffer contating 75 mM sodium phosphate, 1mM EGTA, and 1mM Mgcl2 (pH
% 7.2). The experiments may conducted at 37c. In these experiments at the 
% different Oxaloacetate concentrations, the ROS production and succinate oxidation
% activity is seen. Succinate is 100uM, and A5 = 250nM is used.

function model = S2013_F4C(fun,p,model)
p = updateparams(p,'s2013');

% experimental details
p.T             = 273.15 + 37;              % 37 C
p.Etot          = 12e-9*p.xSiebe;           % M (100 pmol/mg x 0.12 mg/ml x 1e-12 mol/pmol / 1000 mL/L)
p.atpenin       = 250e-9;                   % M

Oxaloacetate    = linspace(0,30,101).*1e-6; % M
Succinate       = 100e-6;                   % M

x = zeros(numel(fieldnames(p.in)),numel(Oxaloacetate)); 
x(p.in.O2,:)            = 200e-6;           % M
x(p.in.Succinate,:)     = Succinate;        % M
x(p.in.H,:)             = 10^(-7.2);        % ROS data could be at 7.2, details are unclear
x(p.in.Oxaloacetate,:)  = Oxaloacetate;     % M
x(p.in.QH2,:)           = 20e-3*(1./(1+ p.Q./Succinate)).*(1./(1+ p.atpenin./p.Q_atpenin)).*(1./(1+ Oxaloacetate./p.Q_OAA));  % M
x(p.in.Q,:)             = 20e-3 - x(p.in.Q,:);                                                                                % M 

[~,JSO,JH2O2,~,~,details] = fun(x,p);
JH2O2tot = JH2O2 + JSO/2;

% format model output
model.label{numel(model.label)+1} = 'Siebels13_fig4Cset1a.txt';
model.Y{numel(model.Y)+1}         = [Oxaloacetate.*1e6; JH2O2tot*1e12.*60/0.12/1e3]'; % pmol.min^-1.mg.protein
model.g{numel(model.g)+1}         = details;

p.atpenin       = 0;                                                                                                  % M
% Oxaloacetate inhibits QH2 production, so QH2 should not be high when malate is high
x(p.in.QH2,:)   = 20e-3*(1./(1+ p.Q./Succinate)).*(1./(1+ p.atpenin./p.Q_atpenin)).*(1./(1+ Oxaloacetate./p.Q_OAA));  % M
x(p.in.Q,:)     = 20e-3 - x(p.in.Q,:);                                                                                % M 

[~,~,~,JQH2,~,details] = fun(x,p);

% format model output
model.label{numel(model.label)+1} = 'Siebels13_fig4Cset2a.txt';
model.Y{numel(model.Y)+1}         = [Oxaloacetate.*1e6; JQH2/JQH2(1)*100]';
model.g{numel(model.g)+1}         = details;


%% Simulate Grivennikota, et al. 2017
% SMPs are obtaied from the bovine heart mitochondria. The assay mixture
% contain 0.25 M sucrose, 10 mM potssium chloride, 0.1 mM EDTA, 
% 5mM potassium phosphate, pH(7.5). 0.25mg/ml of mitochondria were added to
% the assay mixture. The experiments were conducted at 30 C

function model = G2017_F2(fun,p,model)
p = updateparams(p,'G2017');

% experimental details
p.T             = 273.15 + 30;                              % 30 C
p.Etot          = 10e-9*p.xGrivennikova;                    % M (0.1 mg/mL x 1000 mL/L x 100 pmol/mg x 1e-12 mol/pmol)
p.atpenin       = 0;

Succinate   = logspace(-9,log10(25),201).*1e-3;             % M
x = zeros(numel(fieldnames(p.in)),numel(Succinate));

x(p.in.O2,:)        = 200e-6;                               % M
x(p.in.Succinate,:) = Succinate;                            % M
x(p.in.QH2,:)       = 20e-3*(1./(1+ p.Q_stig./Succinate));  % M
x(p.in.Q,:)         = 20e-3 - x(p.in.QH2,:);                % M 
x(p.in.H,:)         = 10^(-7.5);                            % pH = 7.5

[~,JSO,JH2O2,~,~,details] = fun(x,p);
JH2O2tot = JH2O2 + JSO/2; 

% format model output
model.label{numel(model.label)+1} = 'Grivennikova17_fig2Bset1a.txt';
model.Y{numel(model.Y)+1}         = [Succinate.*1e6; JH2O2tot.*1e9.*60/0.1/1e3]'; % nmol/mg/min (Succinate is in uM) 
model.g{numel(model.g)+1}         = details;

%% Simulate Grivennikota, et al. 2017
% SMPs are obtaied from the bovine heart mitochondria. The assay mixture
%(0.25 M sucrose, 50 mM Tris-Cl (pH 8.0), 0.2 mM EDTA, 10 uM rotenone, 1.6 uM
% myxothiazol,SOD (6 units/ml) was added. Atpenin A5 (1 uM) is also present. 
% (0.1 mg of protein per ml). The experiments were conducted at 30 C

function model = G2017_F5(fun,p,model)
p = updateparams(p,'G2017');

p.T         = 273.15 + 30;              % 30 C
p.Etot      = 10e-9*p.xGrivennikova;    % M (0.1mg/mL x 1000 mL/L x 100 pmol/mg x 1e-12 mol/pmol)
p.atpenin   = 1e-6;                     % M

labels = {'Grivennikova17_fig5set1a.txt' 'Grivennikova17_fig5set1b.txt'};

% simulate independent variable(s)
Fumarate = linspace(0,60,101)*1e-6;
Succinate = [2 5].*1e-6; % M

% create arrays for state variables and fluxes  
x = zeros(numel(fieldnames(p.in)),numel(Fumarate));

% specify experimental details
x(p.in.O2,:)        = 200e-6;      % M
x(p.in.Fumarate,:)  = Fumarate;    % M
x(p.in.H,:)         = 10^(-8);
x(p.in.Malonate,:)  = 0.5e-6;      % M (10 mg/ml stock SMP contained 50 uM, diluted 100x in assay buffer)

JH2O2       = zeros(numel(Succinate),numel(Fumarate));
JSO         = zeros(numel(Succinate),numel(Fumarate));
JH2O2tot    = zeros(numel(Succinate),numel(Fumarate));

for i=1:numel(Succinate)
    x(p.in.Succinate,:) = Succinate(i);                                                                                     % M
    x(p.in.QH2,:) = 20e-3*(1./(1+ p.Q_stig./Succinate(i))).*(1./(1+ p.atpenin./p.Q_atpenin)).*(1./(1+ Fumarate./p.Q_FUM));  % M
    x(p.in.Q,:) =  20e-3 - x(p.in.QH2,1);                                                                                   % M
    
    [~, JSO(i,:),JH2O2(i,:),~,~,details{i}]= fun(x,p);
    JH2O2tot(i,:) = JH2O2(i,:) + JSO(i,:)/2; 
    
    % format model output
    model.label{numel(model.label)+1} = labels{i};
    model.Y{numel(model.Y)+1} = [Fumarate.*1e6; JSO(i,:)*1e9.*60/0.1/1e3]'; % nmol.min^-1.mg.protein
    model.g{numel(model.g)+1}  = details{i};
end

%% Simulate Grivennikota, et al. 2017
%The reaction mixture was composed of 20 mM potassium phosphate (pH 7.8),
%0.2 mM EDTA,50 uM malonate, potassium succinate (1�20 mM), 
%(0.25 ug of protein per ml). The experiments were conducted at 30 C

function model = G2017_F6B(fun,p,model)
p = updateparams(p,'G2017F6B');

% experimental details
p.T         = 273.15 + 30;                  % 30 C
p.Etot      = 1.87e-9*p.xGrivennikovaSQR;   % M (0.25 ug SQR/mL x 1000 mL/L x 1e-6 g/ug / 134,000 g/mol (MW))
p.atpenin   = 0;

% define model output according to data
labels = {'Grivennikova17_fig6Bset1a.txt' 'Grivennikova17_fig6Bset1b.txt' 'Grivennikova17_fig6Bset1c.txt'};

Succinate   = linspace(0,30,101).*1e-3;     % M
PMS         = [0.05 0.2 2]*1e-3;            % M

x = zeros(numel(fieldnames(p.in)),numel(Succinate));
J = zeros(numel(PMS),numel(Succinate));

x(p.in.O2,:)        = 200e-6;               % M 
x(p.in.Q,:)         = 1e-99;                % M
x(p.in.Succinate,:) = Succinate;            % M
x(p.in.Malonate,:)  = 50e-6;                % M      
x(p.in.H,:)         = 10^(-7.8);            % pH = 7.8

for i = 1:numel(PMS)
    x(p.in.GEA,:)       = PMS(i);           % M
    [~,~,~,~,J(i,:),d]  = fun(x,p);
    
    model.label{numel(model.label)+1} = labels{i};
    model.Y{numel(model.Y)+1}         = [Succinate.*1e3; J(i,:)*1e6*60/0.25e-3/1e3]'; % umol/min/mg SQR protein
    model.g{numel(model.g)+1}         = d;
end


%% Mitochondria and SMP were assayed in the standard 
% reaction mixture composed of 0.25 M sucrose, 5 mM 
% potassium phosphate,10 mM KCl, and 0.1 mM EDTA (pH 7.5).
% succinate (50uM) in the presence of 5 uM rotenone and 1.6 uM myxothiazol. 
% SMP (40ug/ml) ug of protein per ml 
% The experiments were conducted at 30 C

function model = G2018_F2C(fun,p,model)

% experimental details
p.T         = 273.15 + 30; % 30 C
p.Etot      = 8.5e-10; % M based on comparing '17 and '18 papers
p.atpenin   = 0;

% define model output according to data
labels = {'Grivennikova18_fig2Cset1a.txt'};

Succinate   = 50e-6;                                        % M
O2          = linspace(0,800,801)*1e-6;

x = zeros(numel(fieldnames(p.in)),numel(O2));

x(p.in.O2,:)        = O2;                                   % M 
x(p.in.QH2,:)       = 20e-3*(1./(1+ p.Q_stig./Succinate));  % M
x(p.in.Q,:)         = 20e-3 - x(p.in.QH2,:);                % M 
x(p.in.Succinate,:) = Succinate;                            % M
x(p.in.H,:)         = 10^(-7.5);

[~,JSO,JH2O2,~,~,d] = fun(x,p);
JH2O2tot = JH2O2 + JSO/2;
    
model.label{numel(model.label)+1} = labels{1};
model.Y{numel(model.Y)+1}         = [O2.*1e6; JH2O2tot.*1e9.*60/0.04/1e3]'; % nmol/min/mg SMP
model.g{numel(model.g)+1}         = d;


%% Update kinetic constates when exogenous electron acceptor is used

function p = updateparams(p,flag)

switch flag
    case 'S1968'
         p.kf_GEA = p.kf_PMS;
         p.Em0.GEA = 358; % mV
    case 'V1979'
          p.kf_GEA = p.kf_WB;
          p.Em0.GEA = 270; % mV
    case 'G2017F6B'
         p.kf_GEA = p.kf_PMS;
         p.Em0.GEA = 358; % mV
end

%% Wrapper function to simulate experimental protocols

function model = getoutput(fun,p,fittingfunctions)

%% Initialize model cell array
model.label = {}; % string used to identify model outputs
model.Y = {}; % model output used for fitting
model.g = {}; % misc model outputs set by user

%% Loop through fittingfunctions
for i=1:numel(fittingfunctions)
    model = fittingfunctions{i}(fun,p,model);
end

%% ODE model used for Grivennikova 2018 simulations

function f = sdh_ode(t,x,p)

% initialize LHS
f = zeros(size(x,1),size(x,2));

[J,JSO,JH2O2,JQH2] = main_complexII(x,p);


% Evaluate the right hand side.
f(p.in.Succinate,:) = -J; % conc wrt to aqueous phase
f(p.in.Fumarate,:)  = J;  % conc wrt to aqueous phase

% succinate oxidase
JO2 = 30e-6/60*(x(p.in.QH2,:)./(x(p.in.QH2,:) + 1e-6)).*(x(p.in.O2,:)./(x(p.in.O2,:) + 0.5e-6)); % M/s

% Qpool volume much less than aqueous phase
% 250 nl lipid volume per mg mito(SMP), 0.2 mg/ml SMP
f(p.in.Q,:)    = (2*JO2 - JQH2)/(50e-6);
f(p.in.QH2,:)  = (JQH2 - 2*JO2)/(50e-6);

f(p.in.O2,:)   = -(JSO+JH2O2+JO2) + 1e-9; % O2 consumption plus small background influx to avoid negative values
f(p.in.H2O2,:) = (JSO/2+JH2O2);
```
