## Supplementary material for "Analysis of Mammalian Succinate Dehydrogenase Kinetics and Reactive Oxygen Species Production": Source code: main_complexII.m

```
%% Complex II
% model of kinetics and free radical generation
%
% Copyright:
% Jason Bazil November 2019
% Bazil Laboratory
% Department of Physiology
% Michigan State University

function varargout = main_complexII(x,p)

n = size(x,2); % Need this for substate array construction

%% Solution of the matrix 
O2            = x(p.in.O2,:);            
SO            = x(p.in.SO,:);            
H2O2          = x(p.in.H2O2,:);           
Q             = x(p.in.Q,:);                
QH2           = x(p.in.QH2,:); 
H             = x(p.in.H,:);
Succinate     = x(p.in.Succinate,:);      
Fumarate      = x(p.in.Fumarate,:);       
Malonate      = x(p.in.Malonate,:);
Oxaloacetate  = x(p.in.Oxaloacetate,:);
GEA           = x(p.in.GEA,:);
Malate        = x(p.in.Malate,:);

GED           = 0; % no data set has any non-zero general electron donor concentrations

%% Proton dependency

xH            = 1./(1+H./p.KD_H);
p.kf_GEA      = p.kf_GEA;

%% Binding Polynomials for enzyme, substrates, products and regulators

BP_Q     = (1 + Q./p.KD_Q + QH2./p.KD_QH2 + p.atpenin./p.KD_atpenin);

p.kf_QH2 = p.kf_QH2.*(1 + p.atpenin./p.KD_atpenin2)./(1 + p.beta_atpenin*p.atpenin./p.KD_atpenin2);
p.kf_SUC = p.kf_SUC.*(1 + p.atpenin./p.KD_atpenin2)./(1 + p.beta_atpenin*p.atpenin./p.KD_atpenin2).*xH;
 
BP_FAD   = (1 + Succinate./p.KD_SUC + Fumarate./p.KD_FUM + Malate./p.KD_MAL + Malonate./p.KD_MALO  + Oxaloacetate./p.KD_OXA); 

%% FMN Occupancy for SO and H2O2 reactions

p.kf_FADH  = p.kf_FADH./BP_FAD./(1+H./p.K_FADH);
p.kf_FADH2 = p.kf_FADH2./BP_FAD./(1+H./p.K_FADH2);
p.kf_H2O2  = p.kf_H2O2./BP_FAD./(1+H./p.K_FADH2);

%% Adjust redox midpoint potentials

% FAD potential as a pH dependence
Em.FAD_FADH     = p.Em0.FAD_FADH + (p.R.*p.T./p.F).*log(H.*(1+p.K_FADH./H)); 
Em.FADH_FADH2   = p.Em0.FADH_FADH2 + (p.R.*p.T./p.F) *log(H.*(1+H./p.K_FADH2)./(1+H./p.K_FADH));
Em.FAD_FADH2    = p.Em0.FAD_FADH2 + (p.R.*p.T./p.F./2)*log(H.^2.*(1+H./p.K_FADH2).*(1+p.K_FADH./H)./(1+H./p.K_FADH));

Em.Q_SQ         = p.Em0.Q_SQ.*ones(1,n); % pH independent
Em.SQ_QH2       = 2*p.Em0.Q_QH2 + 2*(p.R*p.T./p.F).*log(H) - Em.Q_SQ; % Em of SQ./QH2 couple at given pH

Em.Q_QH2        = p.Em0.Q_QH2 + (p.R*p.T./p.F).*log(H);
Em.Fum_Suc      = p.Em0.Fum_Suc + (p.R*p.T./p.F).*log(H);

% general electron acceptor (phenazine or TMPD in studies used)
p.Em.GEA        = p.Em0.GEA  + (p.R*p.T/2/p.F) * log(H);

% Midpoint potentials for electron transfer of bound states
Emb.SQ_QH2      = Em.SQ_QH2 - (p.R.*p.T./p.F).*log(p.KD_QH2); % electron transfer free energy (mV)
Emb.Q_SQ        = Em.Q_SQ + (p.R.*p.T./p.F).*log(p.KD_Q); % electron transfer free energy (mV)
Emb.Fum_Suc     = Em.Fum_Suc - (p.R.*p.T./p.F/2).*log(p.KD_SUC) + (p.R.*p.T./p.F/2).*log(p.KD_FUM); % electron transfer free energy (mV)

% Equilibrium constants, Keq
Keq.SO_ISC3     = exp(p.F.*(p.Em.O2_SO  - p.Em.ISC3)./(p.R*p.T));
Keq.SO_SQ       = exp(p.F.*(p.Em.O2_SO  - Emb.Q_SQ)./(p.R*p.T));
Keq.SO_FADH     = exp(p.F.*(p.Em.O2_SO - Em.FAD_FADH)./(p.R*p.T));
Keq.SO_FADH2    = exp(p.F.*(p.Em.O2_SO - Em.FADH_FADH2)./(p.R*p.T));
Keq.H2O2_FADH2  = exp(2*p.F.*(p.Em.O2_H2O2 - Em.FAD_FADH2)./(p.R*p.T));
Keq.QH2_ISC3    = exp(p.F.*(Emb.SQ_QH2  - p.Em.ISC3)./(p.R*p.T)); % should be bound state Em for SQ_QH2
Keq.FUM_FADH2   = exp(2*p.F.*(Em.FAD_FADH2 - Emb.Fum_Suc)./(p.R*p.T)); % should be Em.FAD - Emb.Suc
Keq.GEA_ISC     = exp(2*p.F.*(p.Em.GEA - (p.Em.ISC3 + p.Em.ISC1))./( p.R*p.T)); % since Em of ISC2 so low, using ISC1 as second e- source

%% Boltzman redox poise potentials
% compute Eh for Q./SQ reaction
Eh.FAD_FADH     = Em.FAD_FADH;
Eh.FADH_FADH2   = Em.FADH_FADH2;
Eh.ISC1         = p.Em.ISC1*ones(1,n);
Eh.ISC2         = p.Em.ISC2*ones(1,n);
Eh.ISC3         = p.Em.ISC3*ones(1,n);    
Eh.Q_SQ         = Emb.Q_SQ + (p.R.*p.T./p.F).*log(Q./p.KD_Q./BP_Q);

%% RT normalized DeltaG's of all substrates
% FAD_FADH, FADH_FADH2,FAD_FADH2,ISC1,ISC2,ISC3,Q_SQ;
% Eh.FAD_FADH2
DfG1 = -p.F.* [ Eh.FAD_FADH;...
                Eh.FADH_FADH2;...
                Eh.ISC1;...
                Eh.ISC2;...
                Eh.ISC3;...
                Eh.Q_SQ ]./p.R./p.T;

%% The fractional substrates of redox centers for each elecrtonic states
% E0
s0 = 1;
% E1
E1i = DfG1(p.substates.s1,:);
s1 = exp(-E1i)./(ones(size(p.substates.s1,1),1)*sum(exp(-E1i),1));
% E2
E2i = squeeze(sum(reshape(DfG1([p.substates.s2],:),size(p.substates.s2,1),size(p.substates.s2,2),n),2));          
s2 = exp(-E2i)./(ones(size(p.substates.s2,1),1)*sum(exp(-E2i),1));
% E3
E3i = squeeze(sum(reshape(DfG1([p.substates.s3],:),size(p.substates.s3,1),size(p.substates.s3,2),n),2)); 
s3 = exp(-E3i)./(ones(size(p.substates.s3,1),1)*sum(exp(-E3i),1));
% E4
E4i=squeeze(sum(reshape(DfG1([p.substates.s4],:),size(p.substates.s4,1),size(p.substates.s4,2),n),2)); 
s4=exp(-E4i)./(ones(size(p.substates.s4,1),1)*sum(exp(-E4i),1));

%% State Transition Rates
s1_FADH             = sum(s1(p.s1.FADH,:),1);
s1_FAD              = sum(s1(p.s1.FADox,:),1);
s1_SQ               = sum(s1(p.s1.SQ,:),1);
s1_SQempty          = sum(s1(p.s1.SQempty,:),1);
s1_SQempty_ISC3ox   = sum(s1(p.s1.SQempty_ISC3ox,:),1);
s1_ISC1ox_ISC3ox    = sum(s1(p.s1.ISC1ox_ISC3ox,:),1);
s1_ISC3             = sum(s1(p.s1.ISC3,:),1);
s1_ISC3ox           = sum(s1(p.s1.ISC3ox,:),1);

s2_FAD              = sum(s2(p.s2.FADox,:),1);
s2_FADH             = sum(s2(p.s2.FADH,:),1);
s2_FADH2            = sum(s2(p.s2.FADH2,:),1);
s2_SQ               = sum(s2(p.s2.SQ,:),1);
s2_SQempty          = sum(s2(p.s2.SQempty,:),1);
s2_ISC3             = sum(s2(p.s2.ISC3,:),1);
s2_ISC3ox           = sum(s2(p.s2.ISC3ox,:),1);
s2_ISC1_ISC3        = sum(s2(p.s2.ISC1_ISC3,:),1); 
s2_SQ_ISC3          = sum(s2(p.s2.SQ_ISC3,:),1);
s2_SQempty_ISC3ox   = sum(s2(p.s2.SQempty_ISC3ox,:),1);
s2_ISC1ox_ISC3ox    = sum(s2(p.s2.ISC1ox_ISC3ox,:),1);

s3_FAD              = sum(s3(p.s3.FADox,:),1);
s3_FADH             = sum(s3(p.s3.FADH,:),1);
s3_FADH2            = sum(s3(p.s3.FADH2,:),1);
s3_SQ               = sum(s3(p.s3.SQ,:),1);
s3_SQempty          = sum(s3(p.s3.SQempty,:),1);
s3_ISC3             = sum(s3(p.s3.ISC3,:),1);
s3_ISC3ox           = sum(s3(p.s3.ISC3ox,:),1);
s3_ISC1_ISC3        = sum(s3(p.s3.ISC1_ISC3,:),1);
s3_SQ_ISC3          = sum(s3(p.s3.SQ_ISC3,:),1);

s4_FADH             = sum(s4(p.s4.FADH,:),1);
s4_FADH2            = sum(s4(p.s4.FADH2,:),1);
s4_SQ               = sum(s4(p.s4.SQ,:),1);
s4_ISC3             = sum(s4(p.s4.ISC3,:),1);
s4_ISC1_ISC3        = sum(s4(p.s4.ISC1_ISC3,:),1);
s4_SQ_ISC3          = sum(s4(p.s4.SQ_ISC3,:),1);

%% Unidirectional state transition rates
% 1) E0->E1
% SO + Q -> O2 + SQ
% FAD + H + SO -> FADH + O2

k01_SO_ISC3   = (p.kf_SO2./Keq.SO_ISC3).* 1./BP_Q .* SO .* s0;
k01_SO_Q      = (p.kf_SO./Keq.SO_SQ).* Q./p.KD_Q./BP_Q .* SO .* s0;
k01_SO_FAD    = (p.kf_FADH./Keq.SO_FADH).* SO .* s0;
k0_1          = k01_SO_Q + k01_SO_FAD + k01_SO_ISC3;

% 2) E1->E0
% O2 + SQ -> SO + Q
% FADH + O2 -> FAD + H + SO

k10_O2_ISC3   = p.kf_SO2 .* O2 .* s1_ISC3 ./ BP_Q;
k10_O2_SQ     = p.kf_SO .* O2 .* s1_SQ;
k10_O2_FADH   = p.kf_FADH .* O2 .* s1_FADH;
k1_0          = k10_O2_SQ  + k10_O2_FADH + k10_O2_ISC3;

% 3) E0->E2                       
% H2O2 + FAD -> O2 + FADH2
% FAD + Succinate -> FADH2 + fumarate
% QH2 + [4Fe_3S]ox -> SQ + [4Fe_3S]red + 2H
% GED + [4Fe_3S]ox + [3Fe_4S]ox -> GEA  + [4Fe_3S]red + [3Fe_4S]red

k02_H2O2_FAD           = (p.kf_H2O2./Keq.H2O2_FADH2) .* H2O2 .* s0;
k02_SUC_FAD            =  p.kf_SUC.* Succinate./p.KD_SUC./BP_FAD .* s0;
k02_QH2_SQempty_ISC3ox =  (p.kf_QH2./ Keq.QH2_ISC3).* QH2./p.KD_QH2./BP_Q .* s0;
k02_GED                = (p.kf_GEA./Keq.GEA_ISC).*GED.*s0;
k0_2                   = k02_H2O2_FAD + k02_SUC_FAD + k02_QH2_SQempty_ISC3ox +  k02_GED; 

% 4) E2->E0
% O2 + FADH2 ->  H2O2 + FAD
% FADH2 + fumarate -> FAD + Succinate
% SQ-[4Fe_3S]red + 2H -> QH2 + [4Fe_3S]ox
% GEA + [4Fe_3S]red + [3Fe_4S]red = GED + [4Fe_3S]ox + [3Fe_4S]ox

k20_O2_FADH2        = p.kf_H2O2 .* O2 .* s2_FADH2;
k20_FUM_FADH2       = (p.kf_SUC./ Keq.FUM_FADH2).* Fumarate./p.KD_FUM./BP_FAD .* s2_FADH2; 
k20_SQ_ISC3_QH2     =  p.kf_QH2 .* s2_SQ_ISC3;
k20_GEA             = p.kf_GEA.*GEA.*s2_ISC1_ISC3;
k2_0                = k20_O2_FADH2 + k20_FUM_FADH2 + k20_SQ_ISC3_QH2 + k20_GEA; 

% 5) E1->E2 
% SO + Q -> O2 + SQ
% SO + FAD + H -> FADH + O2
% SO + FADH + H -> FADH2 + O2

k12_SO_ISC3        = (p.kf_SO2./Keq.SO_ISC3) .* 1./BP_Q .* SO .* s1_SQempty;
k12_SO_Q           = (p.kf_SO./Keq.SO_SQ) .* Q./p.KD_Q./BP_Q .* SO .* s1_SQempty;
k12_SO_FAD         = (p.kf_FADH./Keq.SO_FADH).* SO .* s1_FAD;
k12_SO_FADH        = (p.kf_FADH./Keq.SO_FADH2).* SO .* s1_FADH;
k1_2               = k12_SO_Q + k12_SO_FAD + k12_SO_FADH + k12_SO_ISC3;

% 6) E2->E1
% O2 + SQ -> SO + Q
% O2 + FADH -> FAD + H + SO
% O2 + FADH2 -> FADH + H+ SO

k21_O2_ISC3        = p.kf_SO2 .* O2 .* s2_ISC3 ./  BP_Q;
k21_O2_SQ          = p.kf_SO .* O2 .* s2_SQ;
k21_O2_FADH        = p.kf_FADH .* O2 .* s2_FADH;
k21_O2_FADH2       = p.kf_FADH2 .* O2 .* s2_FADH2;
k2_1               = k21_O2_SQ  + k21_O2_FADH + k21_O2_FADH2 + k21_O2_ISC3;

% 7) E1->E3
% H2O2 + FAD -> O2 + FADH2
% FAD + Succinate -> FADH2 + fumarate
% QH2 + [4Fe_3S]ox -> SQ + [4Fe_3S]red + 2H
% GED + [4Fe_3S]ox + [3Fe_4S]ox -> GEA  + [4Fe_3S]red + [3Fe_4S]red

k13_H2O2_FAD           = (p.kf_H2O2./Keq.H2O2_FADH2) .* H2O2 .* s1_FAD;
k13_SUC_FAD            =  p.kf_SUC.* Succinate./p.KD_SUC./BP_FAD .* s1_FAD; 
k13_QH2_SQempty_ISC3ox = (p.kf_QH2./ Keq.QH2_ISC3).* QH2./p.KD_QH2./BP_Q .* s1_SQempty_ISC3ox;
k13_GED                = (p.kf_GEA./Keq.GEA_ISC).*GED.*s1_ISC1ox_ISC3ox; 
k1_3                   = k13_H2O2_FAD + k13_SUC_FAD + k13_QH2_SQempty_ISC3ox + k13_GED; 

% 8) E3->E1
% O2 + FADH2 ->  H2O2 + FAD
% FADH2 + fumarate -> FAD + Succinate
% SQ-[4Fe_3S]red + 2H -> QH2 + [4Fe_3S]ox
% GEA + [4Fe_3S]red + [3Fe_4S]red = GED + [4Fe_3S]ox + [3Fe_4S]ox

k31_O2_FADH2       = p.kf_H2O2 .* O2 .* s3_FADH2;
k31_FUM_FADH2      = (p.kf_SUC./ Keq.FUM_FADH2).* Fumarate./p.KD_FUM./BP_FAD .* s3_FADH2;
k31_SQ_ISC3_QH2    = p.kf_QH2.* s3_SQ_ISC3;
k31_GEA            = p.kf_GEA.*GEA.*s3_ISC1_ISC3;
k3_1               = k31_O2_FADH2 + k31_FUM_FADH2 + k31_SQ_ISC3_QH2 + k31_GEA;

% 9) E2->E3
% SO + ISC3ox -> O2 + ISC3red
% SO + Q -> O2 + SQ
% SO + FAD + H -> FADH + O2
% SO + FADH + H -> FADH2 + O2

k23_SO_ISC3        = (p.kf_SO2./Keq.SO_ISC3) .* 1./BP_Q .* SO .* s2_SQempty;
k23_SO_Q           = (p.kf_SO./Keq.SO_SQ) .* Q./p.KD_Q./BP_Q .* SO .* s2_SQempty;
k23_SO_FAD         = (p.kf_FADH./Keq.SO_FADH).* SO .* s2_FAD;
k23_SO_FADH        = (p.kf_FADH./Keq.SO_FADH2).* SO .* s2_FADH;
k2_3               = k23_SO_Q + k23_SO_FADH + k23_SO_FAD + k23_SO_ISC3;

% 10) E3->E2
% O2 + SQ -> SO + Q
% O2 + FADH -> FAD + H+ SO
% O2 + FADH2 -> FADH + H + SO

k32_O2_ISC3        = p.kf_SO2 .* O2 .* s3_ISC3 ./ BP_Q;
k32_O2_SQ          = p.kf_SO .* O2 .* s3_SQ;
k32_O2_FADH        = p.kf_FADH .* O2 .* s3_FADH;
k32_O2_FADH2       = p.kf_FADH2 .* O2 .* s3_FADH2;
k3_2               = k32_O2_SQ + k32_O2_FADH + k32_O2_FADH2;

% 11) E2->E4
% H2O2 + FAD -> O2 + FADH2
% FAD + Succinate -> FADH2 + fumarate
% QH2 + [4Fe_3S]ox -> SQ + [4Fe_3S]red + 2H
% GED + [4Fe_3S]ox + [3Fe_4S]ox -> GEA  + [4Fe_3S]red + [3Fe_4S]red

k24_H2O2_FAD           = (p.kf_H2O2./Keq.H2O2_FADH2) .* H2O2 .* s2_FAD;
k24_SUC_FAD            =  p.kf_SUC.* Succinate./p.KD_SUC./BP_FAD .* s2_FAD; 
k24_QH2_SQempty_ISC3ox = (p.kf_QH2./Keq.QH2_ISC3) .* QH2./p.KD_QH2./BP_Q .* s2_SQempty_ISC3ox;
k24_GED                = (p.kf_GEA./Keq.GEA_ISC).*GED.*s2_ISC1ox_ISC3ox; 
k2_4                   = k24_H2O2_FAD + k24_SUC_FAD + k24_QH2_SQempty_ISC3ox + k24_GED;

% 12) E4->E2
% O2 + FADH2 ->  H2O2 + FAD
% FADH2 + fumarate -> FAD + Succinate
% SQ-[4Fe_3S]red + 2H -> QH2 + [4Fe_3S]ox
% GEA + [4Fe_3S]red + [3Fe_4S]red = GED + [4Fe_3S]ox + [3Fe_4S]ox

k42_O2_FADH2           = p.kf_H2O2 .* O2 .* s4_FADH2;
k42_FUM_FADH2          = (p.kf_SUC./ Keq.FUM_FADH2).* Fumarate./p.KD_FUM./BP_FAD .* s4_FADH2; 
k42_SQ_ISC3_QH2        =  p.kf_QH2.* s4_SQ_ISC3;
k42_GEA                = p.kf_GEA.*GEA.*s4_ISC1_ISC3; 
k4_2                   = k42_O2_FADH2 + k42_FUM_FADH2 + k42_SQ_ISC3_QH2 + k42_GEA;

% 13) E3->E4
% SO + Q -> O2 + SQ
% SO + FAD + H -> FADH + O2
% SO + FADH + H -> FADH2 + O2

k34_SO_ISC3            = (p.kf_SO2./Keq.SO_ISC3) .* 1./BP_Q .* SO .* s3_SQempty;
k34_SO_Q               = (p.kf_SO./Keq.SO_SQ) .* Q./p.KD_Q./BP_Q .* SO .* s3_SQempty;
k34_SO_FAD             = (p.kf_FADH./Keq.SO_FADH).* SO .* s3_FAD;
k34_SO_FADH            =  p.kf_FADH2 .*SO .* s3_FADH;
k3_4                   =  k34_SO_Q + k34_SO_FAD + k34_SO_FADH + k34_SO_ISC3;

% 14) E4->E3
% O2 + SQ -> SO + Q
% O2 + FADH -> FAD + H + SO
% O2 + FADH2 -> FADH + H+ SO

k43_O2_ISC3            = p.kf_SO2 .* O2 .* s4_ISC3 ./ BP_Q;
k43_O2_SQ              = p.kf_SO .* O2 .* s4_SQ;
k43_O2_FADH            = p.kf_FADH .* O2 .* s4_FADH;
k43_O2_FADH2           = p.kf_FADH2 .* O2 .* s4_FADH2;
k4_3                   = k43_O2_SQ + k43_O2_FADH + k43_O2_FADH2;

% electron occupancy steady states
D = (k0_2.*k1_0.*k2_4.*k3_1 + k0_1.*k1_2.*k2_4.*k3_1 + k0_1.*k1_3.*k2_0.*k3_4 + k0_2.*k1_0.*k2_4.*k3_2 + k0_1.*k1_2.*k2_4.*k3_2 + k0_1.*k1_3.*k2_1.*k3_4 + k0_2.*k1_0.*k2_3.*k3_4 + k0_2.*k1_2.*k2_4.*k3_1 + k0_1.*k1_2.*k2_3.*k3_4 + k0_1.*k1_3.*k2_4.*k3_2 + k0_2.*k1_0.*k2_4.*k3_4 + k0_2.*k1_2.*k2_4.*k3_2 + k0_2.*k1_3.*k2_1.*k3_4 + k0_1.*k1_2.*k2_4.*k3_4 + k0_1.*k1_3.*k2_3.*k3_4 + k0_2.*k1_2.*k2_3.*k3_4 + k0_2.*k1_3.*k2_4.*k3_2 + k0_1.*k1_3.*k2_4.*k3_4 + k0_2.*k1_2.*k2_4.*k3_4 + k0_2.*k1_3.*k2_3.*k3_4 + k0_2.*k1_3.*k2_4.*k3_4 + k0_1.*k1_3.*k2_0.*k4_2 + k0_1.*k1_3.*k2_0.*k4_3 + k0_1.*k1_3.*k2_1.*k4_2 + k0_2.*k1_0.*k2_3.*k4_2 + k0_1.*k1_2.*k2_3.*k4_2 + k0_1.*k1_3.*k2_1.*k4_3 + k0_2.*k1_0.*k2_3.*k4_3 + k0_2.*k1_3.*k2_1.*k4_2 + k0_1.*k1_2.*k2_3.*k4_3 + k0_1.*k1_3.*k2_3.*k4_2 + k0_2.*k1_0.*k2_4.*k4_3 + k0_2.*k1_2.*k2_3.*k4_2 + k0_2.*k1_3.*k2_1.*k4_3 + k0_1.*k1_2.*k2_4.*k4_3 + k0_1.*k1_3.*k2_3.*k4_3 + k0_2.*k1_2.*k2_3.*k4_3 + k0_2.*k1_3.*k2_3.*k4_2 + k0_1.*k1_3.*k2_4.*k4_3 + k0_2.*k1_2.*k2_4.*k4_3 + k0_2.*k1_3.*k2_3.*k4_3 + k0_2.*k1_3.*k2_4.*k4_3 + k0_2.*k1_0.*k3_1.*k4_2 + k0_1.*k1_2.*k3_1.*k4_2 + k0_2.*k1_0.*k3_1.*k4_3 + k0_2.*k1_0.*k3_2.*k4_2 + k0_1.*k1_2.*k3_1.*k4_3 + k0_1.*k1_2.*k3_2.*k4_2 + k0_2.*k1_0.*k3_2.*k4_3 + k0_2.*k1_2.*k3_1.*k4_2 + k0_1.*k1_2.*k3_2.*k4_3 + k0_1.*k1_3.*k3_2.*k4_2 + k0_2.*k1_0.*k3_4.*k4_2 + k0_2.*k1_2.*k3_1.*k4_3 + k0_2.*k1_2.*k3_2.*k4_2 + k0_1.*k1_2.*k3_4.*k4_2 + k0_1.*k1_3.*k3_2.*k4_3 + k0_2.*k1_2.*k3_2.*k4_3 + k0_2.*k1_3.*k3_2.*k4_2 + k0_1.*k1_3.*k3_4.*k4_2 + k0_2.*k1_2.*k3_4.*k4_2 + k0_2.*k1_3.*k3_2.*k4_3 + k0_2.*k1_3.*k3_4.*k4_2 + k0_1.*k2_0.*k3_1.*k4_2 + k0_1.*k2_0.*k3_1.*k4_3 + k0_1.*k2_0.*k3_2.*k4_2 + k0_1.*k2_1.*k3_1.*k4_2 + k0_1.*k2_0.*k3_2.*k4_3 + k0_1.*k2_1.*k3_1.*k4_3 + k0_1.*k2_1.*k3_2.*k4_2 + k0_2.*k2_1.*k3_1.*k4_2 + k0_1.*k2_0.*k3_4.*k4_2 + k0_1.*k2_1.*k3_2.*k4_3 + k0_1.*k2_3.*k3_1.*k4_2 + k0_2.*k2_1.*k3_1.*k4_3 + k0_2.*k2_1.*k3_2.*k4_2 + k0_1.*k2_1.*k3_4.*k4_2 + k0_1.*k2_3.*k3_1.*k4_3 + k0_2.*k2_1.*k3_2.*k4_3 + k0_2.*k2_3.*k3_1.*k4_2 + k0_1.*k2_4.*k3_1.*k4_3 + k0_2.*k2_1.*k3_4.*k4_2 + k0_2.*k2_3.*k3_1.*k4_3 + k0_2.*k2_4.*k3_1.*k4_3 + k1_0.*k2_0.*k3_1.*k4_2 + k1_0.*k2_0.*k3_1.*k4_3 + k1_0.*k2_0.*k3_2.*k4_2 + k1_0.*k2_1.*k3_1.*k4_2 + k1_0.*k2_0.*k3_2.*k4_3 + k1_0.*k2_1.*k3_1.*k4_3 + k1_0.*k2_1.*k3_2.*k4_2 + k1_2.*k2_0.*k3_1.*k4_2 + k1_0.*k2_0.*k3_4.*k4_2 + k1_0.*k2_1.*k3_2.*k4_3 + k1_0.*k2_3.*k3_1.*k4_2 + k1_2.*k2_0.*k3_1.*k4_3 + k1_2.*k2_0.*k3_2.*k4_2 + k1_0.*k2_1.*k3_4.*k4_2 + k1_0.*k2_3.*k3_1.*k4_3 + k1_2.*k2_0.*k3_2.*k4_3 + k1_3.*k2_0.*k3_2.*k4_2 + k1_0.*k2_4.*k3_1.*k4_3 + k1_2.*k2_0.*k3_4.*k4_2 + k1_3.*k2_0.*k3_2.*k4_3 + k1_3.*k2_0.*k3_4.*k4_2);
E0= (k1_0.*k2_0.*k3_1.*k4_2 + k1_0.*k2_0.*k3_1.*k4_3 + k1_0.*k2_0.*k3_2.*k4_2 + k1_0.*k2_1.*k3_1.*k4_2 + k1_0.*k2_0.*k3_2.*k4_3 + k1_0.*k2_1.*k3_1.*k4_3 + k1_0.*k2_1.*k3_2.*k4_2 + k1_2.*k2_0.*k3_1.*k4_2 + k1_0.*k2_0.*k3_4.*k4_2 + k1_0.*k2_1.*k3_2.*k4_3 + k1_0.*k2_3.*k3_1.*k4_2 + k1_2.*k2_0.*k3_1.*k4_3 + k1_2.*k2_0.*k3_2.*k4_2 + k1_0.*k2_1.*k3_4.*k4_2 + k1_0.*k2_3.*k3_1.*k4_3 + k1_2.*k2_0.*k3_2.*k4_3 + k1_3.*k2_0.*k3_2.*k4_2 + k1_0.*k2_4.*k3_1.*k4_3 + k1_2.*k2_0.*k3_4.*k4_2 + k1_3.*k2_0.*k3_2.*k4_3 + k1_3.*k2_0.*k3_4.*k4_2)./D;
E1= (k0_1.*k2_0.*k3_1.*k4_2 + k0_1.*k2_0.*k3_1.*k4_3 + k0_1.*k2_0.*k3_2.*k4_2 + k0_1.*k2_1.*k3_1.*k4_2 + k0_1.*k2_0.*k3_2.*k4_3 + k0_1.*k2_1.*k3_1.*k4_3 + k0_1.*k2_1.*k3_2.*k4_2 + k0_2.*k2_1.*k3_1.*k4_2 + k0_1.*k2_0.*k3_4.*k4_2 + k0_1.*k2_1.*k3_2.*k4_3 + k0_1.*k2_3.*k3_1.*k4_2 + k0_2.*k2_1.*k3_1.*k4_3 + k0_2.*k2_1.*k3_2.*k4_2 + k0_1.*k2_1.*k3_4.*k4_2 + k0_1.*k2_3.*k3_1.*k4_3 + k0_2.*k2_1.*k3_2.*k4_3 + k0_2.*k2_3.*k3_1.*k4_2 + k0_1.*k2_4.*k3_1.*k4_3 + k0_2.*k2_1.*k3_4.*k4_2 + k0_2.*k2_3.*k3_1.*k4_3 + k0_2.*k2_4.*k3_1.*k4_3)./D;
E2= (k0_2.*k1_0.*k3_1.*k4_2 + k0_1.*k1_2.*k3_1.*k4_2 + k0_2.*k1_0.*k3_1.*k4_3 + k0_2.*k1_0.*k3_2.*k4_2 + k0_1.*k1_2.*k3_1.*k4_3 + k0_1.*k1_2.*k3_2.*k4_2 + k0_2.*k1_0.*k3_2.*k4_3 + k0_2.*k1_2.*k3_1.*k4_2 + k0_1.*k1_2.*k3_2.*k4_3 + k0_1.*k1_3.*k3_2.*k4_2 + k0_2.*k1_0.*k3_4.*k4_2 + k0_2.*k1_2.*k3_1.*k4_3 + k0_2.*k1_2.*k3_2.*k4_2 + k0_1.*k1_2.*k3_4.*k4_2 + k0_1.*k1_3.*k3_2.*k4_3 + k0_2.*k1_2.*k3_2.*k4_3 + k0_2.*k1_3.*k3_2.*k4_2 + k0_1.*k1_3.*k3_4.*k4_2 + k0_2.*k1_2.*k3_4.*k4_2 + k0_2.*k1_3.*k3_2.*k4_3 + k0_2.*k1_3.*k3_4.*k4_2)./D;
E3= (k0_1.*k1_3.*k2_0.*k4_2 + k0_1.*k1_3.*k2_0.*k4_3 + k0_1.*k1_3.*k2_1.*k4_2 + k0_2.*k1_0.*k2_3.*k4_2 + k0_1.*k1_2.*k2_3.*k4_2 + k0_1.*k1_3.*k2_1.*k4_3 + k0_2.*k1_0.*k2_3.*k4_3 + k0_2.*k1_3.*k2_1.*k4_2 + k0_1.*k1_2.*k2_3.*k4_3 + k0_1.*k1_3.*k2_3.*k4_2 + k0_2.*k1_0.*k2_4.*k4_3 + k0_2.*k1_2.*k2_3.*k4_2 + k0_2.*k1_3.*k2_1.*k4_3 + k0_1.*k1_2.*k2_4.*k4_3 + k0_1.*k1_3.*k2_3.*k4_3 + k0_2.*k1_2.*k2_3.*k4_3 + k0_2.*k1_3.*k2_3.*k4_2 + k0_1.*k1_3.*k2_4.*k4_3 + k0_2.*k1_2.*k2_4.*k4_3 + k0_2.*k1_3.*k2_3.*k4_3 + k0_2.*k1_3.*k2_4.*k4_3)./D;
E4= 1 -(E0+E1+E2+E3);

%% Compute steady-state rates

% succinate oxidation rate
J = p.Etot.*(k02_SUC_FAD.*E0 + k13_SUC_FAD.*E1 + k24_SUC_FAD.*E2 ...
           - k20_FUM_FADH2.*E2 - k31_FUM_FADH2.*E3 - k42_FUM_FADH2.*E4);

% Superoxide formation
JSO = p.Etot.*((k10_O2_ISC3 + k10_O2_SQ + k10_O2_FADH).*E1 ...
             + (k21_O2_ISC3 + k21_O2_SQ + k21_O2_FADH + k21_O2_FADH2).*E2 ...
             + (k32_O2_ISC3 + k32_O2_SQ + k32_O2_FADH + k32_O2_FADH2).*E3 ...
             + (k43_O2_ISC3 + k43_O2_SQ + k43_O2_FADH + k43_O2_FADH2).*E4 ...
             - (k01_SO_ISC3 + k01_SO_Q + k01_SO_FAD).*E0 ...
             - (k12_SO_ISC3 + k12_SO_Q + k12_SO_FAD + k12_SO_FADH).*E1 ...
             - (k23_SO_ISC3 + k23_SO_Q + k23_SO_FAD + k23_SO_FADH).*E2 ...
             - (k34_SO_ISC3 + k34_SO_Q + k34_SO_FAD + k34_SO_FADH).*E3 );
            
% Hydrogen peroxide formation
JH2O2 = p.Etot.*(k20_O2_FADH2.*E2 + k31_O2_FADH2.*E3 + k42_O2_FADH2.*E4 ...
               - k02_H2O2_FAD.*E0 - k13_H2O2_FAD.*E1 - k24_H2O2_FAD.*E2);
               
%QH2 reduction              
JQH2 = p.Etot.*(k20_SQ_ISC3_QH2.*E2 + k31_SQ_ISC3_QH2.*E3 + k42_SQ_ISC3_QH2.*E4...
              - k02_QH2_SQempty_ISC3ox.*E0 - k13_QH2_SQempty_ISC3ox.*E1 - k24_QH2_SQempty_ISC3ox.*E2);
% GEA reduction
JGEA = p.Etot .*(k20_GEA.*E2 + k31_GEA.*E3 + k42_GEA.*E4... 
               - k02_GED.*E0 - k13_GED.*E1 - k24_GED.*E2);

%%  the stuff might be interested  
           
JISC3_SO = p.Etot.*(k10_O2_ISC3.*E1 + k21_O2_ISC3.*E2  + k32_O2_ISC3.*E3 + k43_O2_ISC3.*E4);

% SO formation from Q site
JQSO = p.Etot.*(k10_O2_SQ.*E1 + k21_O2_SQ.*E2  + k32_O2_SQ.*E3 + k43_O2_SQ.*E4);

% Superoxide formation from FADH site 
JFADH_SO = p.Etot.*(k10_O2_FADH.*E1 + k21_O2_FADH.*E2 + k32_O2_FADH .*E3 + k43_O2_FADH.*E4); 

%Superoxide formation from FADH2 site 
JFADH2_SO = p.Etot.*((k21_O2_FADH2).*E2 + (k32_O2_FADH2).*E3 + (k43_O2_FADH2).*E4);

% Hydrogen peroxide formation
JFADH2_H2O2 = p.Etot.*(k20_O2_FADH2.*E2 + k31_O2_FADH2.*E3 + k42_O2_FADH2.*E4);
                   

%% Collect model details and package into structure

% midpoint potentials
details.Em=Em;
details.Emb=Emb;

% equilibrium constants
details.Keq=Keq;

% substate occupancies
details.s0=s0;
details.s1=s1;
details.s2=s2;
details.s3=s3;
details.s4=s4;

% substate redox fractions
details.s1_FADH               = s1_FADH;
details.s1_FAD                = s1_FAD;
details.s1_SQ                 = s1_SQ;
details.s1_SQempty            = s1_SQempty;
details.s1_SQempty_ISC3ox     = s1_SQempty_ISC3ox;
details.s1_ISC3               = s1_ISC3;
details.s1_ISC3ox             = s1_ISC3ox;
details.s2_FAD                = s2_FAD;
details.s2_FADH               = s2_FADH;
details.s2_FADH2              = s2_FADH2;
details.s2_SQ                 = s2_SQ;
details.s2_SQempty            = s2_SQempty;
details.s2_ISC3               = s2_ISC3;
details.s2_ISC3ox             = s2_ISC3ox;
details.s2_SQ_ISC3            = s2_SQ_ISC3;
details.s2_SQempty_ISC3ox     = s2_SQempty_ISC3ox;
details.s3_FAD                = s3_FAD;
details.s3_FADH               = s3_FADH;
details.s3_FADH2              = s3_FADH2;
details.s3_SQ                 = s3_SQ;
details.s3_SQempty            = s3_SQempty;
details.s3_ISC3               = s3_ISC3;
details.s3_ISC3ox             = s3_ISC3ox;
details.s3_SQ_ISC3            = s3_SQ_ISC3;
details.s4_FADH               = s4_FADH;
details.s4_FADH2              = s4_FADH2;
details.s4_SQ                 = s4_SQ;
details.s4_ISC3               = s4_ISC3;
details.s4_SQ_ISC3            = s4_SQ_ISC3;

% state occupancies
details.E0                    = E0;
details.E1                    = E1;
details.E2                    = E2;
details.E3                    = E3;
details.E4                    = E4;

%total redox state fractions
details.E1_FADH               = E1.*s1_FADH;
details.E1_SQ                 = E1.*s1_SQ;
details.E2_FADH               = E2.*s2_FADH;
details.E2_FADH2              = E2.*s2_FADH2;
details.E2_SQ                 = E2.*s2_SQ;
details.E3_FADH               = E3.*s3_FADH;
details.E3_FADH2              = E3.*s3_FADH2;
details.E3_SQ                 = E3.*s3_SQ;
details.E4_FADH               = E4.*s4_FADH;
details.E4_FADH2              = E4.*s4_FADH2;
details.E4_SQ                 = E4.*s4_SQ;

%total redox fractions
details.FADH                  = E1.*s1_FADH + E2.*s2_FADH + E3.*s3_FADH + E4.*s4_FADH;
details.SQ                    = E1.*s1_SQ + E2.*s2_SQ + E3.*s3_SQ + E4.*s4_SQ;
details.FADH2                 = E2.*s2_FADH2 + E3.*s3_FADH2 + E4.*s4_FADH2;

% turnover at each site
details.J                      = J;
details.JSO                    = JSO;
details.JH2O2                  = JH2O2;
details.JQH2                   = JQH2;
details.JGEA                   = JGEA;

% Partial reactions 
% 1)E0->E1
details.k01_SO_Q               = k01_SO_Q;
details.k01_SO_FAD             = k01_SO_FAD;
details.k0_1                   = k0_1;

% 2)E1->E0
details.k10_O2_SQ              = k10_O2_SQ;
details.k10_O2_FADH            = k10_O2_FADH;
details.k1_0                   = k1_0;

% 3)E0->E2
details.k02_H2O2_FAD           = k02_H2O2_FAD;
details.k02_SUC_FAD            = k02_SUC_FAD ;
details.k02_QH2_SQempty_ISC3ox = k02_QH2_SQempty_ISC3ox;
details.k02_GED                = k02_GED; 
details.k0_2                   = k0_2;

% 4)E2->E0
details.k20_O2_FADH2           = k20_O2_FADH2;
details.k20_FUM_FADH2          = k20_FUM_FADH2; 
details.k20_SQ_ISC3_QH2        = k20_SQ_ISC3_QH2; 
details.k20_GED                = k20_GEA; 
details.k2_0                   = k2_0; 

% 5) E1->E2 
details.k12_SO_Q               = k12_SO_Q;
details.k12_SO_FAD             = k12_SO_FAD;
details.k12_SO_FADH            = k12_SO_FADH;
details.k1_2                   = k1_2;

% 6)E2->E1
details.k21_O2_SQ              = k21_O2_SQ;
details.k21_O2_FADH            = k21_O2_FADH;
details.k21_O2_FADH2           = k21_O2_FADH2;
details.k2_1                   = k2_1;

% 7)E1->E3
details.k13_H2O2_FAD           = k13_H2O2_FAD;
details.k13_SUC_FAD            = k13_SUC_FAD; 
details.k13_QH2_SQempty_ISC3ox = k13_QH2_SQempty_ISC3ox;
details.k13_GED                = k13_GED; 
details.k1_3                   = k1_3;

% 8)E3->E1
details.k31_O2_FADH2           = k31_O2_FADH2;
details.k31_FUM_FADH2          = k31_FUM_FADH2;
details.k31_SQ_ISC3_QH2        = k31_SQ_ISC3_QH2;
details.k31_GED                = k31_GEA;
details.k3_1                   = k3_1;

% 9)E2->E3
details.k23_SO_Q               = k23_SO_Q;
details.k23_SO_FAD             = k23_SO_FAD;
details.k23_SO_FADH            = k23_SO_FADH;
details.k2_3                   = k2_3;

% 10)E3->E2
details.k32_O2_SQ              = k32_O2_SQ;
details.k32_O2_FADH            = k32_O2_FADH;
details.k32_O2_FADH2           = k32_O2_FADH2;
details.k3_2                   = k3_2;

% 11)E2->E4
details.k24_H2O2_FAD           = k24_H2O2_FAD;
details.k24_SUC_FAD            = k24_SUC_FAD; 
details.k24_QH2_SQempty_ISC3ox = k24_QH2_SQempty_ISC3ox;
details.k24_GED                = k24_GED;
details.k2_4                   = k2_4;
% 12)E4->E2
details.k42_O2_FADH2           = k42_O2_FADH2;
details.k42_FUM_FADH2          = k42_FUM_FADH2; 
details.k42_SQ_ISC3_QH2        = k42_SQ_ISC3_QH2;
details.k42_GED                = k42_GEA;
details.k4_2                   = k4_2;
% 13)E3->E4
details.k34_SO_Q               = k34_SO_Q;
details.k34_SO_FAD             = k34_SO_FAD ;
details.k34_SO_FADH            = k34_SO_FADH;
details.k3_4                   = k3_4;
% 14)E4->E3
details.k43_O2_SQ              = k43_O2_SQ;
details.k43_O2_FADH            = k43_O2_FADH;
details.k43_O2_FADH2           = k43_O2_FADH2;
details.k4_3 = k4_3;

details.JISC3_SO               = JISC3_SO;
details.JQSO                   = JQSO;
details.JFADH_SO               = JFADH_SO;
details.JFADH2_SO              = JFADH2_SO;
details.JFADH2_H2O2            = JFADH2_H2O2;
details.x                      = x;

%% output
if nargout > 1
    varargout = {J, JSO, JH2O2, JQH2, JGEA,details,JQSO,JFADH_SO, JFADH2_SO,JFADH2_H2O2};
else
    varargout = {J};
end
```
